# Waiting for love but not forever: modelling the evolution of waiting time to selfing in hermaphrodites

**DOI:** 10.1101/2020.08.10.243626

**Authors:** Chantal Blüml, Steven A. Ramm, Koen J. van Benthem, Meike J. Wittmann

**Affiliations:** Department of Theoretical Biology, Bielefeld University, 33615 Bielefeld, Germany; Department of Evolutionary Biology, Bielefeld University, 33615 Bielefeld, Germany

**Keywords:** Waiting time, density-dependent selection, inbreeding depression, individual-based model, mating system, hermaphrodites

## Abstract

Although mixed mating systems involving both selfing and outcrossing are fairly common in hermaphrodites, it is unclear how they are maintained. In some species, individuals delay self-fertilization while they have not found a mating partner. The ‘waiting time’ is subject to two opposing selection pressures: waiting helps to avoid inbreeding depression in offspring by increasing the density-dependent probability to encounter a mate, but also increases the risk of dying before reproduction. In some species waiting time can vary between individuals and be heritable. We therefore used an individual-based model to explore how delayed selfing evolves in response to density and density fluctuations. We find that at high density, when individuals meet often, drift drives waiting time; at intermediate densities, strong inbreeding depression causes waiting time to increase; and at low densities, inbreeding depression is purged, and waiting time approaches zero. Positive feedback loops drive the system to either complete selfing or complete outcrossing. Fluctuating density can slow down convergence to these alternative stable states. However, mixed mating, in the sense of either a stable polymorphism in waiting times, or stable intermediate times, was never observed. Thus, additional factors need to be explored to explain the persistence of delayed selfing.

## Introduction

In many animal and plant species, successful reproduction requires encountering a mating partner or successful transfer of pollen, which can be difficult at low population or pollinator density. Under these conditions, having an alternative reproduction mechanism is potentially advantageous (Darwin 1877; Goodwillie et al. 2005; Jarne and Auld 2006), and in hermaphrodites, self-fertilization (selfing) is one such option. In addition to this reproductive assurance (Darwin 1877; Ramm et al. 2015),selfing also has a transmission advantage (Fisher 1941) and grants reproductive assurance. However, selfed offspring may suffer from inbreeding depression, causing high mortality, reduced fertility and reduced lifespan of offspring (Charlesworth and Charlesworth 1998; Jarne et al. 2000; Ramey et al. 2000; O’Grady et al. 2006; Escobar et al. 2007; Roman and Darling 2007; Dlugosch and Parker 2008; Charlesworth and Willis 2009; Ramm et al. 2012). Ideally, mating behaviour should be adapted to population density, since mixed mating systems encompassing both selfing and outcrossing have the potential both to grant reproductive assurance in low-density situations (Darwin 1877; Coates et al. 2006; Ramm et al. 2015), while also maintaining the advantages of outcrossing – such as the creation of new allele combinations and protection against the consequences of recessive deleterious mutations – when mating partners are available (Goodwillie et al. 2005).

One mechanism by which selfing and outcrossing can be combined in a mixed mating system is so-called ‘delayed selfing’. Here, individuals only begin to self after a certain waiting time has passed, during which they have not encountered other individuals with which to outcross. Mixed mating systems with delayed selfing are present in many species (Goodwillie et al. 2005; Jarne and Auld 2006). Ramm et al. (2012) observed the existence of a substantial waiting time in flatworms of the species *Macrostomum hystrix*. In their experiments, waiting time was heritable, and highly variable across individuals. Similarly, in the freshwater snail *Physa acuta* there is similarly substantial heritable variation among families for age at first reproduction in isolated individuals (Escobar et al. 2007). Other examples of species that express such a waiting time to selfing are the sea slug *Alderia willowi* (Smolensky et al. 2009), the cestode *Schistocephalus solidas (Schjørring 2004)* (Schärer and Wedekind 1999), the plant *Linaria cavanillesii* (Voillemot et al. 2019), and the androdioecious clam shrimp *Eulimnadia texana* (Weeks and Zucker 1999).

The high heritability of waiting times in some of these systems suggests that waiting times to selfing might evolve over time. This evolution should reflect a trade-off between the benefits of producing outcrossed offspring and the (mortality) risk of waiting too long for a potential mating partner. If a partner is encountered before death, having selfed beforehand was potentially a waste of resources that could otherwise have been more profitably invested in non-inbred progeny (Tsitrone et al. 2003b). Those that waited long enough to put all these resources into outcrossed offspring will have a higher fitness. If, additionally, resources not used while waiting can be allocated into increased future offspring number (i.e. if an individual invests in growth), there might be additional reason to delay reproduction (Tsitrone et al. 2003b). However, if an individual’s lifespan is shorter than its waiting time, it would not produce any offspring at all in the absence of a mate. Consequently, waiting may have a large pay-off if it goes well, but is also a risky strategy. Furthermore, individuals with a short waiting time may reproduce early and therefore also produce grand-offspring earlier, shortening their generation time. They then have a smaller lifetime reproductive success, but more descendants per time while minimizing the risk of death before reproduction (Levin et al. 1986; Kot 2012).

An influential body of theory suggests that populations will evolve either towards complete outcrossing or complete selfing, making these mating systems alternative stable states (Lande and Schemske 1985). The reason is that under frequent outcrossing, the recessive deleterious alleles causing inbreeding depression rise in frequency, making selfing more and more costly, whereas under frequent selfing these alleles are purged such that the cost of selfing decreases (Lande and Schemske 1985; Bataillon and Kirkpatrick 2000). Fisher (1941) argued that in a system where selfing is caused by a dominant allele, it should quickly become the only strategy present in the population. Jarne et al. (2000) proposed that the range of conditions leading to a mixed mating system might be quite narrow and the review by Igic and Busch (2013) suggested that inbreeding depression might have to be very large to favour a reversion to outcrossing from established selfing, especially when there are other advantages to selfing besides the transmission advantage. Previous theoretical and empirical studies (Lande and Schemske 1985; Schemske and Lande 1985; Tsitrone et al. 2003a; Goodwillie et al. 2005; Ramm et al. 2012) have identified several mechanisms by which mixed mating systems and stable intermediate selfing rates can be maintained (some specific to plants), but in general the maintenance of mixed mating systems remains puzzling, especially in animals (Jarne and Charlesworth 1993; Lehtonen and Kokko 2012).

In their classical paper, Lande and Schemske (1985) proposed temporal density fluctuations as a possible mechanism for the maintenance of mixed mating systems. In many natural populations, population density fluctuates over time, for example because of disturbances, predator-prey interactions, or seasonality. Thus, there might be times when the number of available mates is low and selfing is favoured and other times when there is no mate limitation and outcrossing is favoured. For fluctuations in pollinator density, stabilization of selfing rates has been shown in models for the evolution of plant mating systems (Morgan and Wilson 2005; Cheptou and Schoen 2007). However, fluctuating selection scenarios where each type is favoured during some time interval do not necessarily lead to the maintenance of variation, but could also cause the fixation of whichever type is best on average (Haldane and Jayakar 1963; Lande 2008; Wittmann et al. 2017). Evaluating whether density fluctuations, and thus fluctuations in mate availability, can maintain a mixed mating system with intermediate or variable waiting times to selfing thus requires explicit modelling, which to our knowledge has not been performed to date (but see Cheptou and Dieckmann (2002), who found intermediate selfing rates for fluctuating density when density affects inbreeding depression rather than mate availability without purging).

Here, we build and analyse an individual-based simulation model for the evolution of waiting times under constant or fluctuating population density. The model is inspired by the reproductive biology of the free-living flatworm *M. hystrix*, but should be applicable more generally to animals and plants with delayed selfing. We first study the evolution of waiting times under constant density to explore whether waiting times go to zero or infinity or to an intermediate optimum value depending on density. For a simplified version of this model, we also perform a mathematical analysis. We then include density fluctuations and again evaluate whether waiting times evolve to extreme or intermediate values and whether heritable variation in waiting times, as observed in species such as *M. hystrix* (Ramm et al. 2012), *P. acuta* (Jarne et al. 2000) and *Clarkia xantiana* (Briscoe Runquist et al. 2017), can be maintained.

## Material & Methods

### Model overview

The individual-based model is stochastic and operates in discrete time with overlapping generations. Each individual has a diploid genome that determines waiting time to selfing and may also carry recessive deleterious mutations that influence offspring viability. In each time unit, which can be interpreted as a day, first potential changes in density are executed by altering the size of the habitat. After that, individuals might die. Then the survivors produce eggs and “search” for a mating partner. The mate encounter probability depends on the current density. After that, reproduction via outcrossing, selfing or use of stored sperm from previous matings takes place and offspring viability is determined. In the following sections, we describe each of these processes in detail.

### Density and density fluctuations

The probability of encountering a mate in our model depends on density, i.e. the number of individuals divided by habitat size. The number of individuals, *N*, is variable, but is kept close to a carrying capacity of *K*=200 individuals, as described below. The habitat size is *g^2^* and we do not assume a particular spatial structure. Instead, *g* is used to determine the probability of encountering a mate as described below. To evaluate the effect of density and density fluctuations on the evolution of waiting time, we manipulate the habitat size rather than the population size, in order to avoid effects of population size on inbreeding depression.

### Death

Each day, individuals die with probability *d=*0.01, representing mortality from predation, old age, disease etc. Lifespan of individuals that have survived the recruitment phase is therefore geometrically distributed with mean 1/*d*.

### Mate search and reproduction

Each individual produces *c=*1 eggs per day, which are stored until the next reproductive event and then released. More generally, *c* can also be interpreted as resources stored for future reproduction. There are three ways to fertilize eggs: [1] outcrossing with a partner, [2] selfing when time since last sperm exchange (through outcrossing or selfing), or since birth for first reproduction, exceeds waiting time, and [3] using sperm from a previous selfing or outcrossing event. In each reproduction phase, first each of the 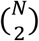 possible pairs of individuals independently meet with probability 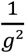. Meeting events are executed in random order and an individual can mate with multiple partners. Second, all individuals who have no sperm stored and not received sperm for longer than their individual waiting time, self-fertilize. Finally, individuals who have sperm stored from a previous outcrossing or selfing use that sperm to fertilize any eggs they may have. Since the model was inspired by flatworms, we assumed that sperm only lasted for *o=10* days (Peters and Michiles 1996; Ramm et al. 2015). Thus, individuals with longer waiting times could go through phases without reproduction, even after having outcrossed or selfed before.

Whenever an individual partakes in a reproductive event, it always releases all its stored eggs. This means selfing or sperm usage opportunities for that day may become obsolete, but individuals can transfer sperm and fertilise their partner’s eggs in multiple outcrossing events. Sperm is always replaced by that of the most recent reproductive event. Outcrossed sperm, however, might be more competitive than selfed sperm (Ramm 2017). This is represented by the fact that although stored sperm is always replaced when outcrossing, individuals will not attempt selfing while having outcrossed sperm stored.

To model density-dependent limits on reproduction, each egg initially has the same chance of surviving, which is calculated by 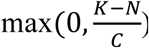, where *K* is the carrying capacity, *N* the population size, and *C* is the cumulative number of eggs laid by all individuals reproducing in that day.

### Genetics

Individuals are diploid. There are two types of loci: *W=5* loci that determine waiting time to selfing and *I=20* loci that can carry deleterious alleles and cause inbreeding depression. Each offspring receives for each locus one allele from each parent (or in the case of selfing, two from the same parent, but alleles are chosen independently).

The alleles at the waiting-time loci represent waiting time relative to the average life span and are represented by positive real numbers. When an offspring is created, mutations occur. Each allele value is multiplied by a random number independently drawn from a gamma distribution with mean 1 and standard deviation of *u*_*w*_=0.05 (shape=400, scale=0.0025). Assuming that the loci act additively, the average allele value across all loci represents an individual’s genotypic value for waiting time. This genotypic value can then be converted to the actual waiting time by multiplying it by the expected lifespan 1/*d*. Consequently, an individual with an average allele value above 1 waits longer than an expected lifespan before starting to self.

For each inbreeding-depression locus, there are *v*=10 alleles: a wild-type allele *w* and *v*-*1=9* alleles, *x*_*1*_,…,*x*_*9*_, that are deleterious when homozygous. Mutations happen with a probability *u*_*i*_=0.03 per allele copy in each new offspring and change an allele with equal probability to any of the other *v*-1 alleles. There are therefore four genotypes classes for each inbreeding-depression locus: *ww* (both wildtype), *wx*_*i*_ (wildtype and any deleterious allele), *x*_*i*_*x*_*j*_ (two different deleterious alleles), and *x*_*i*_*x*_*i*_ (two copies of the same deleterious allele); the first three genotypes have no negative effect on fitness, while the last one affects fitness by a factor 1 – *s*, where *s=0*.*8* is the selection coefficient. Biologically, this corresponds to each deleterious allele causing a different problem, but the other allele being able to compensate for this. The loci then act multiplicatively, so that offspring survival is determined by (1 – *s*)^χ^, and χ is the number of *x*_*i*_*x*_*i*_-loci homozygous for a recessive deleterious mutation.

### Starting conditions

The starting population consists of *N*_0_=*K=*200 individuals. This population size was chosen for computational efficiency, but small changes in *N*_*0*_=*K* had a minor effect (see supporting information Figure N & Figure O) and exploratory analyses with *N*_*0*_=*K*=1000 gave very similar results (not shown). Individuals start out with a time since last reproduction of zero. The number of initially stored eggs is drawn from a Poisson distribution with a mean of 10% of the average life span (i.e. 10) for each individual.

For waiting time in the initial population, first the genotypic values, i.e. average allele values, are determined. These values are chosen to range in evenly spaced steps from 0 to 1 across individuals. For each individual, all allele values are simply set to its genotypic value. This procedure was chosen to ensure consistent and high genetic variation across individuals in the initial population. At each inbreeding locus, each individual in the starting population carries two copies of the wildtype allele. The final “equilibrium” frequency of deleterious mutations depends on many factors such as the selfing frequency (Lande and Schemske 1985), which results from a combination of mate availability, waiting times, purging and the feedback of inbreeding depression. Equilibria usually emerge within ca. 10,000 days.

### Scenarios and simulations

First, we ran the model with various constant habitat sizes *g* in order to understand the effect of density on waiting time and inbreeding depression. To investigate the effect of density fluctuations, we then changed habitat size *g* over time using a step function. This meant that habitat size alternated between two values, remaining constant at each for several days (supporting information Figure A).

The parameter values for our default scenario are given in Table 1. To assess model robustness, we ran a set of additional scenarios, with each parameter altered by ±5%. Additionally, we used an approach where we randomly drew parameter values within a certain range (supporting information Table A), and one where we would increase or decrease waiting time after 30,000 days to see whether or average waiting time would systematically return to its initial state (see supporting information section 10, Figure N-O for results).

**Table 1:** Parameters and their values used for the model. Unless stated otherwise, default values apply.

| Parameter | Symbol | Default value |
| --- | --- | --- |
| Carrying capacity | $K$ | 200 |
| Starting population size | $N_0$ | $K$ |
| Habitat size | $g^2$ | |
| Mortality | $d$ | 0.01 |
| Sperm storage duration | $o$ | 10 |
| Number of eggs produced per day | $c$ | 1 |
| Number of loci for waiting time | $W$ | 5 |
| Number of loci for inbreeding depression | $I$ | 20 |
| Standard deviation for mutation of waiting time loci | $u_w$ | 0.05 |
| Mutation rate at inbreeding loci | $u_i$ | 0.03 |
| “Allelic diversity” Number of alleles at inbreeding loci | $v$ | 10 |
| Selection coefficient | $s$ | 0.8 |

The model was implemented in R 3.5.1 (R Core Team, 2019).

### Mathematical Approximation

To gain a first understanding of the conditions leading to the evolution of higher or lower waiting times, and specify conditions for the individual-based model we analysed a simplified continuous-time version of the model (see supporting information section 2 for details). Mathematical approximations were constructed with the help of Mathematica (Wolfram Research, Inc., Version 11.3, 2018). In brief, time to first meeting *T*_*M*_ and time to death *T*_*D*_ are exponentially distributed with 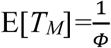, and 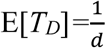, where *Φ* is the meeting probability per day, which depends on population size and habitat size, and *d* is the death rate. Population size *N*, habitat size *g*, and therefore density, are assumed to be constant for the mathematical approximation. We additionally assumed that individuals switch from selfing to outcrossing when meeting another individual, but never switch back to selfing after receiving outcrossed sperm. Sperm, once received, lasts for the entire lifespan of an individual. Offspring resulting from selfing have viability *z*, which we assumed constant, while outcrossed offspring is never affected by inbreeding depression. There was no feedback from inbreeding depression to population density.

This leads to four scenarios for an individual’s life trajectory (supporting information Figure B): 1. the individual dies before reproducing (0 offspring), 2. the individual outcrosses, then dies (number of offspring proportional to life span), 3. the individual selfs then dies (number of offspring proportional to lifespan, but proportional loss due to inbreeding), or 4. the individual selfs, then outcrosses and then dies (selfed portion of offspring with reduced survival due to inbreeding).

We followed an approach very similar to that by Tsitrone et al. (2003a) and focused on female lifetime reproductive success, assuming that reproductive success in the male function is independent of waiting time. We integrated over all possible times of death and meeting to obtain the expected lifetime number of viable offspring (see supporting information section 2 for details) as a function of waiting time τ:

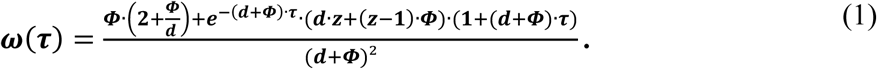

The main difference of our approach compared to Tsitrone et al. (2003a) is that they assume a different effect of waiting on resource allocation, growth, and future reproduction.

To study the direction of selection, we take the fitness gradient, that is, the derivative of fitness with respect to waiting time:

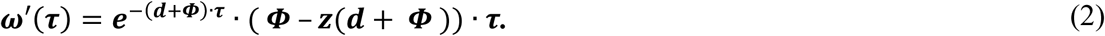

Since the exponential and waiting time are always positive, the direction of selection depends on *Φ* − *z* · (*d* + *Φ*). If this term is positive, i.e. if the meeting probability is above a critical value 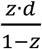, then there is selection towards increased waiting times, and waiting times are expected to increase to infinity. If the meeting probability is below 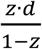, selection acts to reduce waiting times and waiting times are expected to converge to zero. Examples of fitness and fitness gradient under different conditions can be seen in Figure 1.

**Figure 1:**
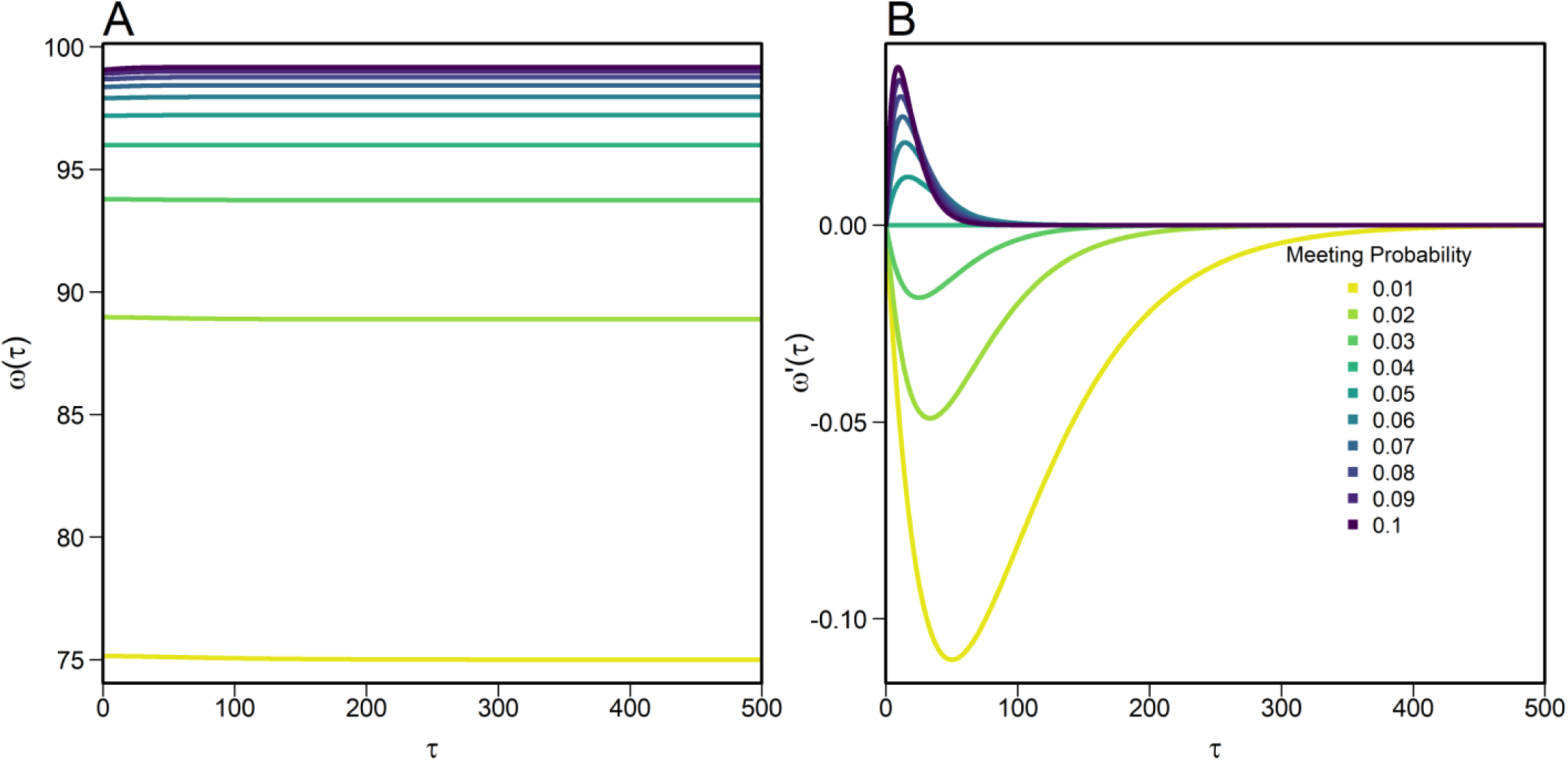
Results from the mathematical approximation, designed to clarify conditions which may favour long or short waiting times. It represents a simplification of the individual-based model. Effects of waiting time (*τ*) on selfing fitness (*ω(τ)*), calculated from formula (1), and its derivative (*ω’(τ)*), calculated from formula (2), under different mate encounter probabilities *Φ*. This mathematical approximation illustrates that fitness ω(τ) is highest at either τ=0 (with low mate availability) or *τ=*infinity (at high mate availability). The critical meeting probability for the parameters we used is *Φ=0.04*. Assumptions are exponentially distributed time of death and time of first meeting, acquired sperm can be used for the rest of an individual’s life, individuals will never self after having outcrossed, outcrossed sperm is used if present, and inbreeding depression is constant. In this example, viability of selfed offspring *z*=0.8 and probability of death h *d=*0.01.

In conclusion, if meeting probability, inbreeding depression, and/or average lifespan are low, waiting time will evolve towards zero, and individuals will immediately self. Meanwhile, high chance of meeting other individuals, strong inbreeding depression or very long lifespans may lead to selfing avoidance, namely an infinite waiting time. In natural scenarios, this may lead to losing the ability to outcross or self, respectively. These results therefore suggest that with constant meeting probability, a population with a range of intermediate waiting times to selfing cannot be explained.

The simplified assumptions of the mathematical model, however, ignore several aspects of the system, such as purging, or the advantage of shorter generation times through earlier reproduction. Moreover, it does not include changes in density that cause Φ to vary between being above or below the threshold. To incorporate these aspects and gain a fuller understanding of the system, including these processes, we now analyse the individual-based model.

## Results

### Waiting time evolution under constant density

We began by exploring how waiting time evolves under constant density. As in the mathematical approximation (see Material & Methods), the evolution of waiting time to selfing in the individual-based model depended on meeting probability, determined by the habitat size *g* (Figure 2). There were three distinct scenarios: low, intermediate and high density.

**Figure 2:**
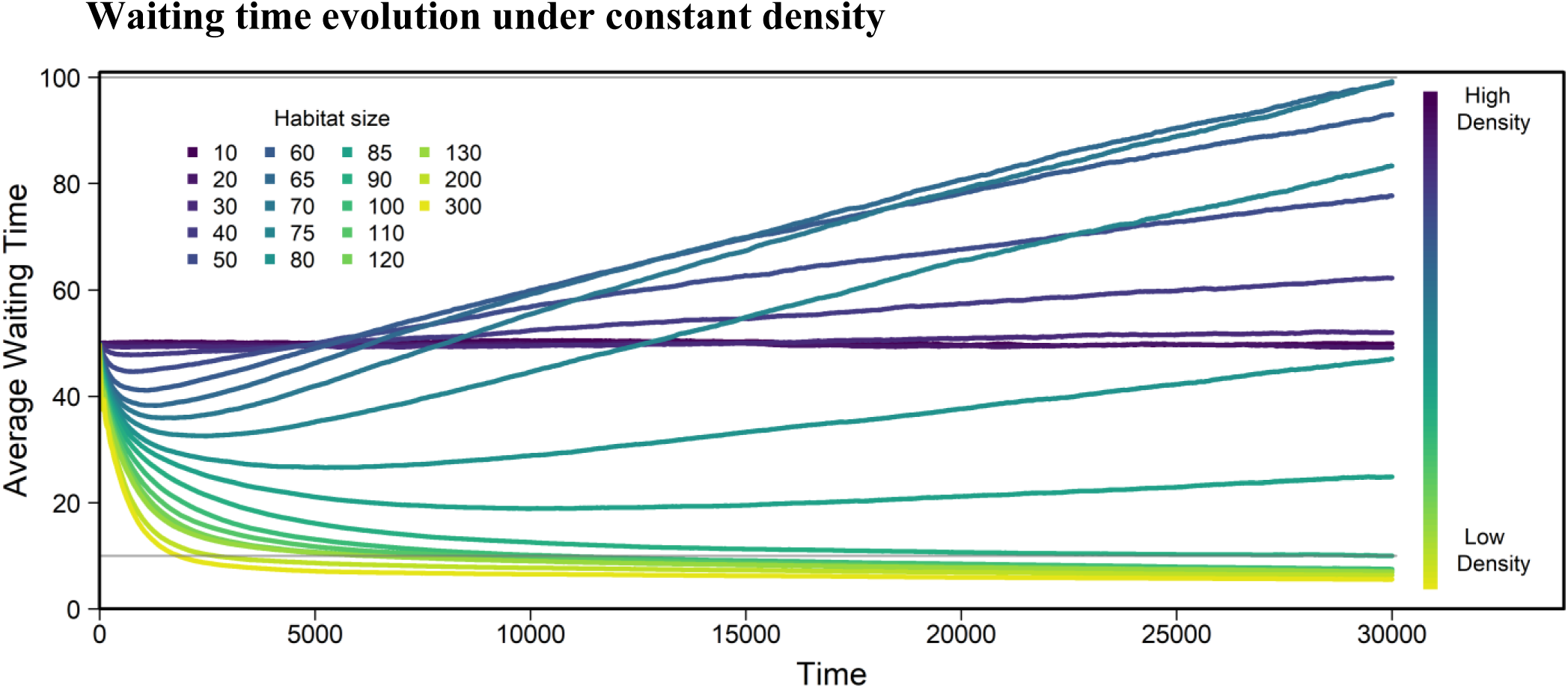
Evolution of average waiting time over all individuals and replicates using the default parameters described in Table 1 and various habitat sizes *g*. Note that habitat sizes are given in terms of area, *g*. The colour bar shows how *g* relates to density. Number of replicates=1000. Generations overlap, but average lifespan is 100 days, meaning the graph shows approximately 300 generations. At low densities (yellow), average waiting time decreases, at intermediate densities (turquoise) it increases, while it stays approximately constant on average for high densities (blue). The grey horizontal line indicates the sperm storage duration.

At low density, i.e. in large habitats, average waiting time decreases over time, converging towards zero or a small nonzero waiting time below the sperm storage time such that they would start selfing immediately once they run out of stored sperm. In such large habitats, individuals are relatively unlikely to ever meet a mating partner (e.g. 0.33 for g=200 in a continuous-time approximation, supporting information Figure C and Equation (6)). Without sperm and/or egg storage, waiting time always approaches zero (supporting information Figure D).

At intermediate density, average waiting time increases over time. Under these conditions, individuals are likely to meet a mating partner within their lifetime (e.g. with probability 0.8 for *g*=70, and 0.75 for *g*=80; supporting information Figure C and Equation (6), continuous time approximation). Importantly, in accordance with the mathematical approximation, no stable waiting time is reached (supporting information Figure E for long term behaviour).

At high density, i.e. in small habitats, average waiting times remain constant. Individuals here have essentially unlimited access to mating partners and individuals typically meet multiple potential mating partners each day. Accordingly, the proportion and number of offspring actually produced by selfing is close to zero (e.g. at *g=*10, the proportion of offspring generated among all replicates over the course of the entire simulation by selfing is 1.92 · 10^−7^).

The rate of change in average waiting time varies with population density (Figure 2), which indicates that the strength of selection differs between densities. Additionally, we observe transient effects in waiting time evolution, especially at intermediate habitat sizes where average waiting time initially decreased before increasing again. This is caused by the frequency of deleterious alleles initially being lower than at equilibrium, thereby reducing the cost of selfing, and favouring short waiting times (supporting information Figure F). In addition, at some point in time, some replicates fix at zero while others keep increasing in waiting time (supporting information Figure H). Those that increase in waiting time then cause the average to increase. Finally, there is a small initial effect on population size (supporting information Figure J), due to the many maladapted individuals present in the diverse starting population.

Results from the robustness analysis where each parameter was varied by ±5% of its default value (Table 1) suggest that small changes in most parameters cause no major changes in average waiting time at the end of the simulation (supporting information Figure N).

### Inbreeding depression and purging

We next examined the implications of waiting-time evolution for the frequency of deleterious alleles and thus inbreeding depression. The frequency of deleterious recessive alleles declines with habitat size (Figure 3), which is expected if in large habitats there is more selfing and thus increased exposure to selection (Lande and Schemske 1985). The transition from densities where deleterious alleles can spread through the population to those where strong selection effectively removes them nearly immediately is quite rapid (supporting information Figure F). Even when selection is strong, however, the total frequency of wildtype alleles only becomes 0.927 (with 0.073 deleterious alleles persisting because of mutation-selection balance). It takes about 10,000 time units for wildtype allele frequencies to stabilize (supporting information Figure F).

**Figure 3:**
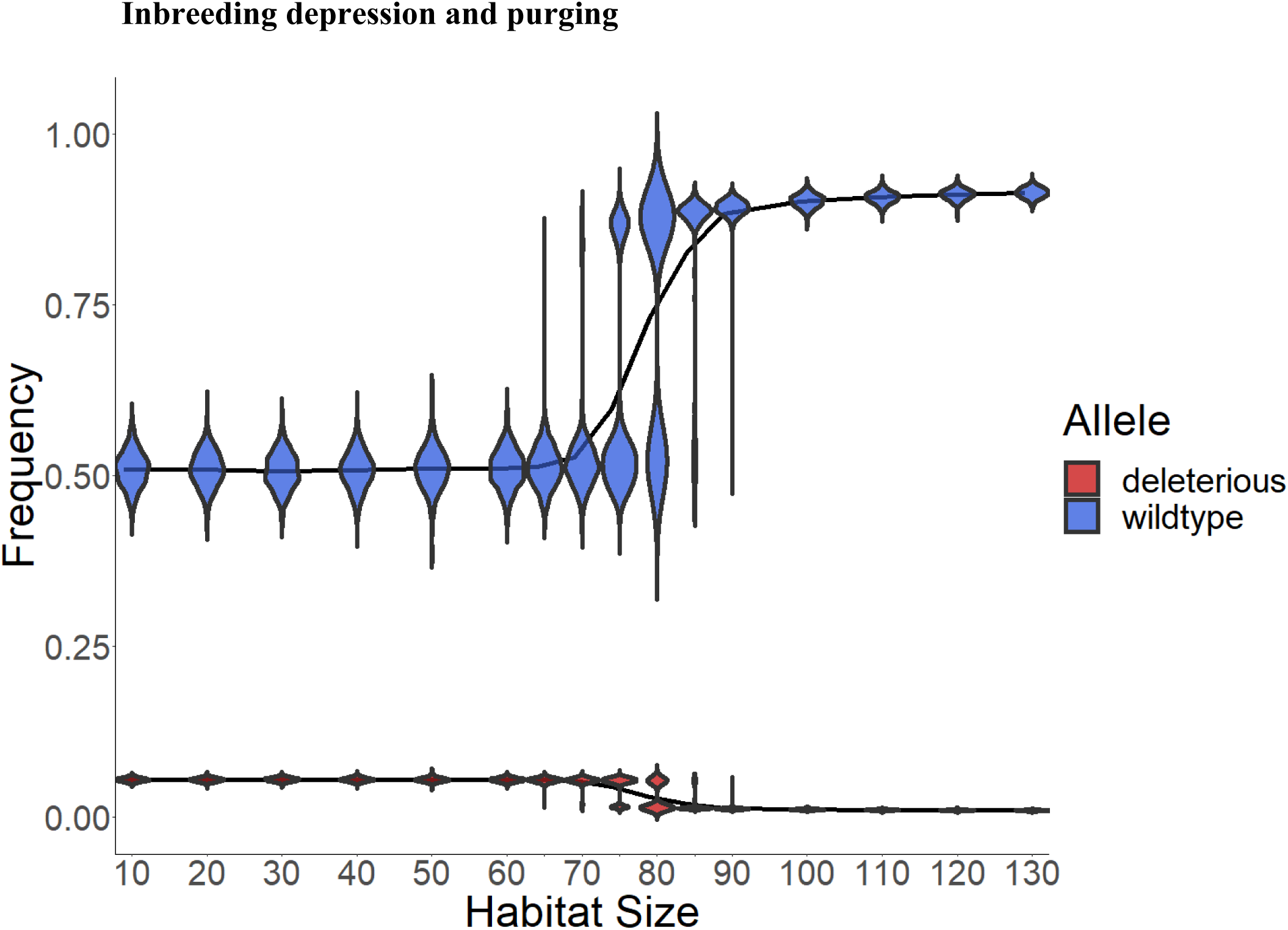
Frequency of wildtype and deleterious alleles at the end of the simulation. All 9 types of deleterious alleles (red) are summarized to average frequency of each mutation. Lines show the average frequencies of all (1000) replicates over the entire simulation time. Violin areas indicate the distribution of allele frequencies among replicates. At low densities (large habitat size *g*), the frequency of wildtype alleles (blue) is high. This is because at low density selfing happens more frequently, and deleterious alleles are purged. Data as in Figure 2. The frequencies of the underlying genotypes are shown in supplementary Figure 8, and the changes in wildtype frequency over time in supporting material Figure G.

In an altered model run, with only two types of alleles per locus (one wildtype and one deleterious rather than the nine in Figure 3), we found that the resulting deleterious allele frequency was smaller (0.065). Consequently, the difference in fitness between outcrossing and selfing individuals was not large enough for selfing avoidance to evolve (supporting information Figure M). To find out how strong selection against selfing needs to be in order for outcrossing to be favoured, we ran additional simulations with fixed mortality rates for selfed offspring and no mortality in mated offspring (supporting information Figure K). The level of inbreeding depression required to drive populations towards outcrossing, i.e. increased waiting times, is close to 60-70% and thus notably higher than the 50% suggested by Lande and Schemske (1985). This may explain why we could not find increasing waiting times with only two variants at each locus, i.e. with just one deleterious allele (supporting information Figure M), as in this case even outcrossed offspring die more often with decreasing wildtype frequency and the difference in viability between outcrossed and selfed offspring is not as large as with multiple deleterious alleles per locus.

### Variation & Trait Distributions

The magnitude of variance in waiting time was largely driven by the magnitude of waiting time itself, which is expected because of the multiplicative mutation process. We therefore used the coefficient of variation (CV) as a more meaningful measure for comparison. The CV was calculated from the variation between individuals within a population and then averaged over replicates. The CV is highest at intermediate densities and lowest for densities where selection favoured short waiting times, i.e. at large habitat sizes (Figure 4B). At high densities (small habitat size), the CV is intermediate.

**Figure 4:**
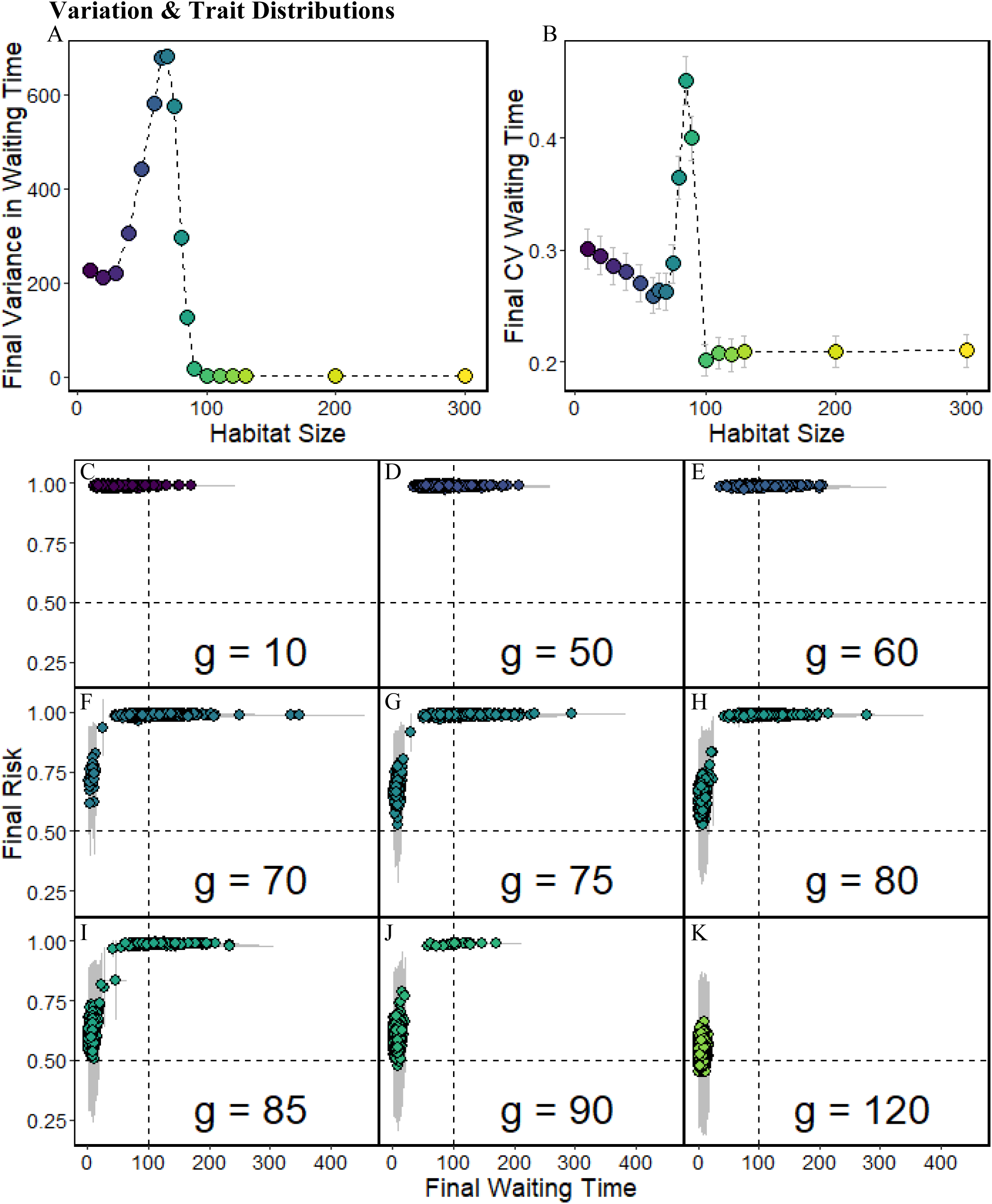
Variation in waiting time. Data as in Figure 2. A: Average variance in waiting time within replicates. B: Average coefficient of variation (CV) in waiting time within replicates. C-K: Points show the average waiting time in the final (last day) population plotted against average individual risk *r*, which is the probability of selfed offspring of an individual dying of inbreeding depression. Each point represents one replicate (note that many points overlap). The grey bars represent within-replicate standard deviations for both x and y-range. Each panel shows a different habitat size *g*.

In Figure 4C-K, we plotted average waiting time against average individual risk *r*. This measured the probability that an offspring resulting from selfing by a given parent dies due to inbreeding depression:

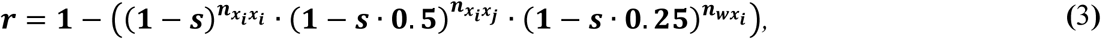

Where 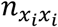 is the number of loci that are homozygous for one of the deleterious mutant alleles in the parent, 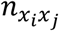 is the number of loci that carry two different deleterious mutations, and 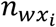 is the number of loci which carry one wildtype and one deleterious allele. Loci of the type *x*_*i*_*x*_*i*_ will always produce the genotype *x*_*i*_*x*_*i*_ in selfed offspring. The combination of *x*_*i*_*x*_*j*_ produces offspring that is homozygous for one of the deleterious alleles with probability 0.5 when selfing. For loci with genotype *wx*_*i*_, the probability of generating offspring with genotype *x*_*i*_*x*_*i*_ is 0.25 when selfing. Each homozygous deleterious locus then decreases the chance that offspring survives and acts multiplicatively with the other loci. Frequencies of genotypes can be found in supporting material Figure G.

In Figure 4C-K, we see that at low habitat sizes, individual risk is high for almost all replicates, while waiting time varies across replicates. For some habitat sizes (here, for example, g=75, and g=80), the waiting-time range is shifted to higher values in comparison to even higher densities (i.e. g=85, and g=90). Here selfing avoidance has evolved. At large habitat sizes, by contrast, most deleterious alleles cannot spread through the population: individual risk is therefore low, as is the waiting time. For a range of intermediate habitat sizes between these extremes, however, both possible outcomes occur in different replicates. Some populations evolve towards high individual risk with long waiting times, whereas others exhibit low individual risk and short waiting time. The larger the habitat is (i.e., the lower the density), the larger the proportion of replicates that follow the latter trajectory.

The temporal dynamics of waiting time for individual replicates (supporting information Figure H) illustrates that the apparent increase in waiting time at the end of the simulation for those densities with alternative stable states occurs because a subset of the trajectories evolve toward infinity, driving the mean upwards, while those trajectories with an average waiting time of zero cannot evolve towards even smaller waiting times. Temporal change in CV and variation in waiting time can be seen in Figure I in the supporting information.

### Fluctuating Density

Finally, we explored how fluctuating density impacts the mean and variance of waiting time. Results for simulations with density fluctuations where *g* alternated between 70 (high density, chance to encounter a mating partner within lifespan is 80.4%) and 300 (low density, chance to encounter a mating partner within lifespan is 18.2%), are shown in Figure 5. In these scenarios, there are long-term trends in waiting time. Evolution towards zero waiting time appears to be faster than evolution towards long waiting times (see also Figure 2). Although waiting time decreases under low density and increases under high density, we found no conditions with perfect balance between these (see also below). The coefficient of variation of waiting time becomes even smaller under fluctuating density than under constant density, as can be seen in Figure 5 B.

**Figure 5:**
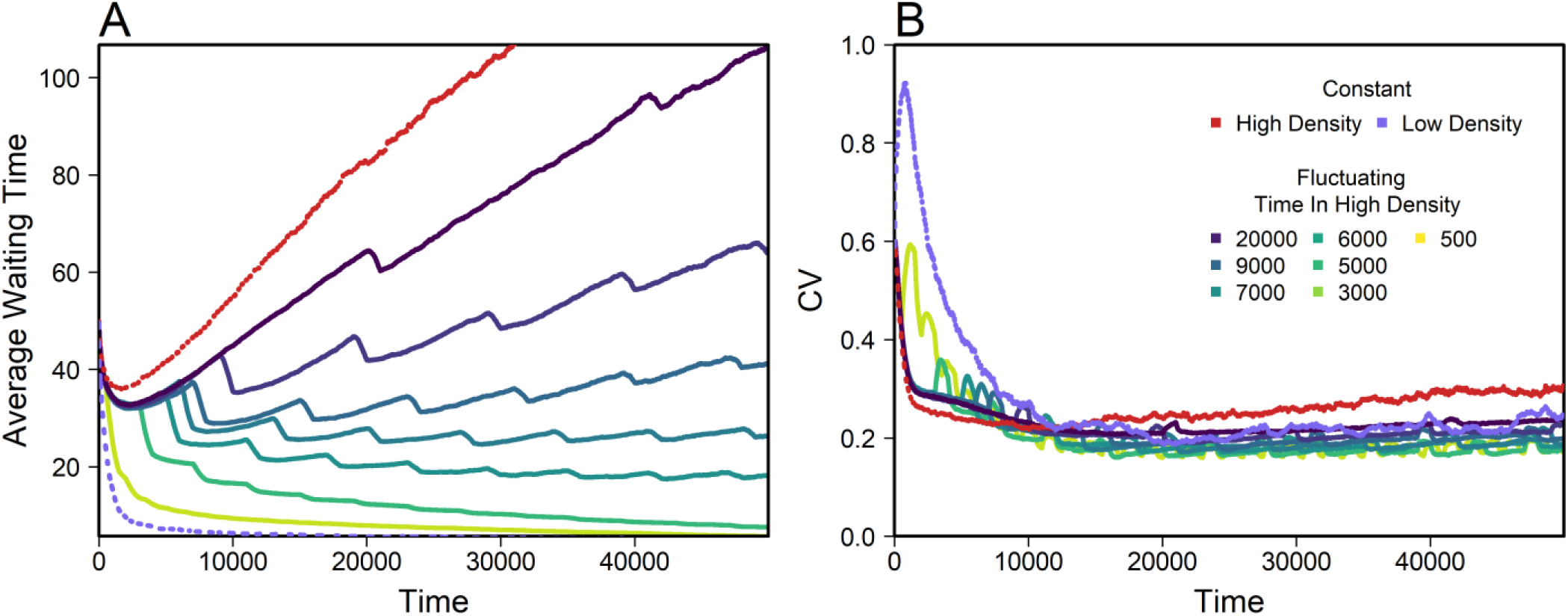
Example of average waiting time (A) and coefficient of variation (CV, B) for different patterns of density fluctuations. Average within replicate CV calculated over all replicates. The colour-scaled lines show evolution for density fluctuations, while the purple and red line show data for constant densities of *g*=70 and *g*=300, which were used as high and low densities. Fluctuations were assembled in a way that populations were exposed to high or low densities for a certain time, before switching to the other situation (example supporting information Figure A). All replicates spent 1000 days in low density and the time in high density was as stated in the legend. All replicates began at high density. Number of replicates=630.

Just as for constant density, we altered all parameters by 5%. Results can be seen in Figure O and show no unexpected or strong influence of any parameter. The ability to store sperm for longer made populations wait longer. Similarly, increases in parameters such as the mutation rate on inbreeding loci or number of inbreeding loci led to increased waiting time. Lastly, an increase in carrying capacity at a given habitat size led to larger densities and meeting probabilities and thus larger waiting times.

In search of parameter combinations that might cause stable intermediate waiting times, we ran 7000 replicates where all parameters were randomly drawn (of which 6577 could be evaluated, as the others went extinct). For the 10 parameter combinations where the result most strongly suggested stable intermediate waiting times (for criteria see supporting information section 10 and Figure P), 100 replicate simulations were run and we found that each replicate had a different waiting time by the end of the simulation (supporting information Figure Q). We tested whether their final values represented a stable equilibrium value by adding or subtracting 10 days from waiting time after an initialisation of 30000 days and then continuing the simulation with this perturbed value to see if they would return to their state before perturbation (supporting information Figure R), which was not the case within the observed timeframe (20000 days). This indicated that waiting time was driven by stochasticity, rather than selection. Closer inspection revealed that the chosen parameter combinations had low mutation rates, further supporting low evolvability.

We also explored the impact of fluctuations between three different densities in a regular or random pattern, but again did not observe stable variation or stable intermediate waiting times (supporting information Figure S).

In summary, fluctuating densities may slow down the evolution of extreme waiting times. However, we did not find evidence for selection favouring intermediate waiting times in the long term.

## Discussion

### Waiting time evolution under constant density

Under constant density, we observed four different evolutionary scenarios but no stable polymorphisms for waiting time to selfing. First, at low density, waiting time decreases towards zero, as waiting is too risky. Second, at high density, mating opportunities are frequent and so waiting time is rarely expressed, such that trait changes are mostly due to genetic drift, which also explains the relatively high final variation in waiting time. Third, at intermediate densities, we find an increase in waiting time towards infinity. Since meeting other individuals eventually is likely, waiting pays off.

Finally, there are also intermediate densities at which two alternative stable states can occur: low waiting times in combination with low (purged) inbreeding depression, or high waiting times combined with high frequency of deleterious alleles (and high inbreeding depression in rare selfing events). Populations that experience the same starting conditions can potentially go either way, depending on stochasticity at the beginning of the simulation. Once a state has established it becomes hard to invade. If most individuals have long waiting times and reproduce mostly via outcrossing, deleterious alleles can drift to higher frequencies, making selfing more costly and leading to even higher waiting times. A mutant with a short waiting time would have a large disadvantage in this system, as its offspring would likely be homozygous for some recessive deleterious mutations, and therefore express inbreeding depression. Even an invader with a short waiting time and purged deleterious alleles would potentially have a disadvantage in later generations, because it might meet a resident and produce offspring with intermediate waiting times and higher mutational load than the invader parent. At the other extreme, i.e. if most individuals have short waiting times and self frequently, recessive deleterious mutations are purged out of the population and selfing no longer has a cost. A mutant with a long waiting time will then not have any advantage and only has a higher risk of death before reproduction. Moreover, it will have a transmission disadvantage because the long-waiting mutant shares the saved-up eggs with its partner, which will have already released most of its eggs in previous selfing events. These results can thus be interpreted as a positive feedback loop (Lehtonen and Kokko 2012), but we emphasize that these alternative stable states (Lande and Schemske 1985; Kokko and Rankin 2006) were never observed in the same population. Thus, under constant density, we did not find evidence for a stable polymorphism of waiting times within populations.

However, waiting time polymorphism is not necessary for a mixed mating system. When waiting times are smaller than the lifespan and mating partners are rare, some individuals will self and others will outcross, even without stable variation in waiting times. The same individual might even self-fertilize and then later fertilize a mating partner’s eggs. When individuals at low density occasionally meet, they will use the opportunity to outcross, even when their waiting time is low. In both cases, a mixed mating system is technically expressed. At relatively high densities, on the other hand, variation in waiting time is high through drift. This does, however, not lead to a mixed mating system, as the time between two outcrossing opportunities will be lower than any waiting time.

Although intermediate waiting times could thus also maintain a mixed mating system, it appears that in our simulations waiting times will eventually either approach zero or exceed the average lifespan. Additionally, natural populations might even completely lose the ability to self or outcross when there are costs to maintaining these mechanisms and they are no longer used. We therefore conclude that in our model under constant density neither optimal intermediate waiting times nor variation in waiting time within a population caused by anything other than stochasticity can be found.

### Dependence on inbreeding depression

Our result that inbreeding depression and density can drive a population to complete selfing or complete outcrossing supports previous work (Lande and Schemske 1985; Jarne and Charlesworth 1993). Furthermore, both our mathematical approximation (Equation (1) & (2)) and the simulations support the conclusion that inbreeding depression needs to be relatively strong in order to systematically drive up waiting times in the intermediate density scenarios (supporting information Figure K). As shown in Figure 4, the inbreeding depression needed for an increase in waiting time seems to be much larger than the 0.5 proposed by Lande and Schemske (1985), which would only balance out the transmission advantage of selfing. It agrees with Tsitrone et al. (2003a) that inbreeding depression high enough to avoid selfing is hard to reach (supporting information Figure F &K), in particular when there are only few deleterious allele variants per locus. When having only two alleles per locus, it can be shown analytically that a frequency of entirely recessive lethal alleles of 50% can be purged down to 10% within three generations of selfing (supplementary information Figure L). Charlesworth et al. (1990) describes a similar model where deleterious alleles occur as closely linked loci, which could be comparable to multiple allele variants.

A positive effect of the level of inbreeding depression on waiting times, where large inbreeding depression is correlated with long waiting times, has been shown in *Physa acuta* (Jarne et al. 2000; Weeks et al. 2001; Escobar et al. 2009; Noël et al. 2016; Noël et al. 2018), and *Eulimnadia texana* (Weeks et al. 1999; Weeks et al. 2001). A recent study in plants (Brown and Kelly 2020) has shown that inbreeding depression caused by rare mutations can be severe enough to meet the criteria shown in Figure K in the supporting information. It should, however, be mentioned that other studies in the same systems did not find links between inbreeding depression and waiting time (Weeks et al. 1999; Escobar et al. 2007). Therefore it is not certain whether waiting time was a consequence of inbreeding depression or other factors, such as environmental stressors, that might potentially interact with inbreeding depression (Armbruster and Reed 2005; Cheptou and Schoen 2007). Alternatively density dependence of inbreeding depression could lead to negative instead of positive feedback loops, as shown by Cheptou and Dieckmann (2002), who did however not include purging and mate limitation, which are driving factors for positive feedback loops in our model.

In this study, we assumed reciprocal mating where both individuals transfer sperm and release eggs, if they have any. Outcrossing would release eggs from both individuals, so twice the amount from selfing. Each outcrossed egg, however, only contained half the genome of each parent. Selfing, on the other hand, generated eggs from only one individual, but with the all of the genes from that parent. Therefore, both strategies get the same genetic output from a reproductive event 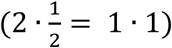 in a very simplified case. The threshold of an inbreeding depression of 0.5, introduced by Lande and Schemske (1985) assumes that half the genetic output is the cost of mating. Such a cost does not occur if reciprocal mating is obligatory. Macrostromum hystrix, the species that initially inspired the model, as well as *Physa acuta*, and *Eulimnadia texana*, do not always perform obligatory reciprocal mating (Weeks and Zucker 1999; Ramm et al. 2012; Noël et al. 2016). In practice, however, in *M. hystrix*, both mating partners can try to inseminate the other individual, so that mating often is reciprocal. In *Physa acuta* however individuals need to choose a role, often leading to conflict between mates (Noël et al. 2016). Moreover, if matings are limited as in our simulation, unmated individuals have an interest in receiving sperm. In order to fertilize their eggs, once two individuals encounter each other, they might stay in spatial proximity, and sequentially mate multiple times. It should be noted that we do not take sex allocation and the cost of sex into account here, which could alter the results as they increase the ability to develop a selfing syndrome (Sicard and Lenhard 2011).

### Waiting time evolution under fluctuating densities

Natural populations of species such as *Macrostomum hystrix* and *Physa acuta* can experience environments strongly fluctuating over time (Sassaman 1989; Henry et al. 2006). Density fluctuations due to changing habitat size, as in our simulation, likely also happen to natural populations, as ponds dry up or fill with rain water (Sassaman 1989). Here, we have mainly considered such temporal variation in population density.

Fluctuations in density lead to fluctuating selection on waiting times, causing them to increase during high-density phases and decrease during low-density phases (Figure 5). Low density affected average waiting time much faster than high density. The major reason for this is that while inbreeding depression that would lead to selfing avoidance at high densities needed to slowly build up again after having been purged out, a lack of mating partners was experienced immediately at low density.

Despite fluctuating density and the resulting fluctuations in selection on waiting time, we did not find patterns of fluctuations that would keep waiting times at an intermediate level for a long time. Instead, waiting time evolved towards one of the two alternative endpoints, similar to the pattern described by Lande and Schemske (1985), although the approach to these endpoints took longer than under constant density (Figure 5). This was especially the case when mutation rates were low (supporting information Figure P). Similarly, intraspecific variation in waiting times was initially lost more slowly under fluctuating density than under constant density, but in the long run fell to similar or lower values. Thus, contrary to the long-standing hypothesis that density fluctuations may promote mixed mating systems, we did not find evidence for permanently stable intermediate waiting times or increased variation in waiting times in our model.

Our results appear to be robust to small parameter changes as well as randomly chosen combinations of parameters. Although this suggests that the absence of stable intermediate waiting times and persistent variation in waiting times might be a general result under our model assumptions, our parameter space was twelve-dimensional and so logistic constraints meant we could only explore a limited part of that parameter space. We did find cases of apparent stable intermediate waiting times, which were, however, not repeatable (supporting information Figure Q), or stable against perturbation (supporting information Figure R), and therefore likely results of slow evolution (supporting information Figure P) and/or stochasticity. Thus, we cannot exclude the possibility that parameter combinations with stable intermediate waiting times or stable variation in waiting times exist somewhere in parameter space. However, if stable intermediate waiting times only occurred for few very specific parameter combinations, the biological relevance in natural populations would be questionable.

We thus conclude that density fluctuations and the resulting fluctuating selection on selfing waiting times are generally not a sufficient condition for the maintenance of mixed mating systems in hermaphrodites. We join Goodwillie et al. (2005) in concluding that reproductive assurance may happen, but nevertheless, the fluctuations may not provide a good basis for maintenance of intermediate waiting times. Even if reproductive assurance was advantageous once in a while, it would be hard to maintain the mechanism over longer timespans. Our results agree with multiple other studies not finding stable intermediate waiting times or selfing rates (Lande and Schemske 1985; Jarne and Charlesworth 1993; Kirkpatrick and Jarne 2000; Igic and Busch 2013). Morgan and Wilson (2005) and Cheptou and Schoen (2007) do find stable intermediate selfing rates with fluctuating pollination probabilities, but they do not include purging and thus appear to have weaker positive feedbacks. Holsinger (1988) also found stable intermediate selfing rates, but only when the locus determining the mating system was closely linked to a locus with heterozygote advantage. With recessive deleterious mutations at unlinked loci, the system went to either complete selfing or outcrossing, as in our study. However, mechanisms other than inbreeding depression, such as the Fisher-Muller-effect, where new allele combinations through sexual reproduction (with conspecifics) can lead to beneficial allele-combinations instead of inbreeding depression, theoretically lead to intermediate rates of sexual reproduction (Kokko 2020).

### Alternative explanations and future work

We speculate that sex allocation and the selfing syndrome (Noël et al. 2016) may further influence the strength of selection of waiting time in either direction. An alternative explanation for the maintenance of variation in waiting time is related to spatial structure (Ronfort and Couvet (1995). In our simulations, we found different final waiting times across replicates for the same density conditions. Migration between otherwise separated populations that have evolved in different directions could lead to intermediate waiting times in their offspring and yield an opportunity to evolve in either direction again. In fact, flatworms collected at different sites show substantial differences in waiting time. While some seem to delay selfing (Ramm et al. 2012), flatworms from other locations do not (Giannakara and Ramm 2020).

Another promising mechanism is pollen discounting, and allocation of resources towards (future) outcrossed reproduction (Gregorius 1982; Holsinger 1991; Tsitrone et al. 2003b; Goodwillie et al. 2005; Dlugosch and Parker 2008; Briscoe Runquist et al. 2017; Huang and Burd 2019; Voillemot et al. 2019). Resource allocation to future reproduction might be crucial for the emergence of intermediate waiting times from a combination of density and inbreeding depression (Tsitrone et al. 2003b). For example, *Physa acuta* uses life-history trade-offs to invest resources into growth instead of egg production (Tsitrone et al. 2003a). This mechanism leads to over-proportional gain in egg production for waiters, compared to the relatively equivalent output for waiters and non-waiters in our model. For plants, Huang and Burd (2019) similarly present an explanation for intermediate selfing rates, for the case that seed size can be adapted. Thereby offspring quality and quantity evolve in accordance with reproductive mode. Moreover, in their study, fertilization success differs between ovules. Similarly, avoiding hypodermic insemination, and the resulting cost of injury may be a reason to delay selfing (Smolensky et al. 2009). All the studies mentioned here have in common that they find intermediate waiting times when adding further life-history trade-offs and allocation of resources to the system. Our model in contrast had the aim to put more focus solely on the decision between outcrossing and selfing, and therefore excluded such mechanisms.

### Conclusions

Using a stochastic individual-based simulation and mathematical approximations, we could show that waiting times to selfing evolve to very small values at low density and to very large values at high density. At intermediate densities, we observed alternative stable states, but no stable polymorphism within populations. We did not find evidence that fluctuating population densities lead to maintenance of variation in waiting times or stable intermediate waiting times. Thus, our results question the hypothesis that density fluctuations can explain the maintenance of a mixed mating system. Variation could then only arise from either drift or variation in optimal time of reproduction. Not taking additional factors that would drive life-history evolution into account, evolution towards alternative stable states in different partially isolated subpopulations seem the most likely explanation of variation within a species.

## Supporting information

Simulation Scripts

## Author contributions

C.B., M.J.W. and S.A.R. designed the project and modelling framework. C.B. designed the computational framework, analysed the data, and drafted the paper. S.A.R. provided insight and discussion on the empirical motivation for the project. K.J.v.B. improved computational efficiency of the code. M.J.W. supervised the project. All authors provided critical feedback and helped shape the research, analysis and manuscript.

## Acknowledgements

We would like to thank the Theoretical Biology group, and in particular Matthias Spangenberg, for discussion and feedback on the project. Moreover, we are thankful for inspiring discussions with Hanna Kokko. This research was partly funded by the German Research Foundation (DFG) as part of the SFB TRR 212 (NC^3^) – project numbers 316099922 and 396782288.

## Data Accessibility

Supporting information containing further figures and simulation code files is provided as a supplement.

## Notes

### Competing Interest Statement

The authors have declared no competing interest.

