## Supporting Information for "Waiting for love but not forever: modelling the evolution of waiting time to selfing in hermaphrodites"

### 1. Step functions

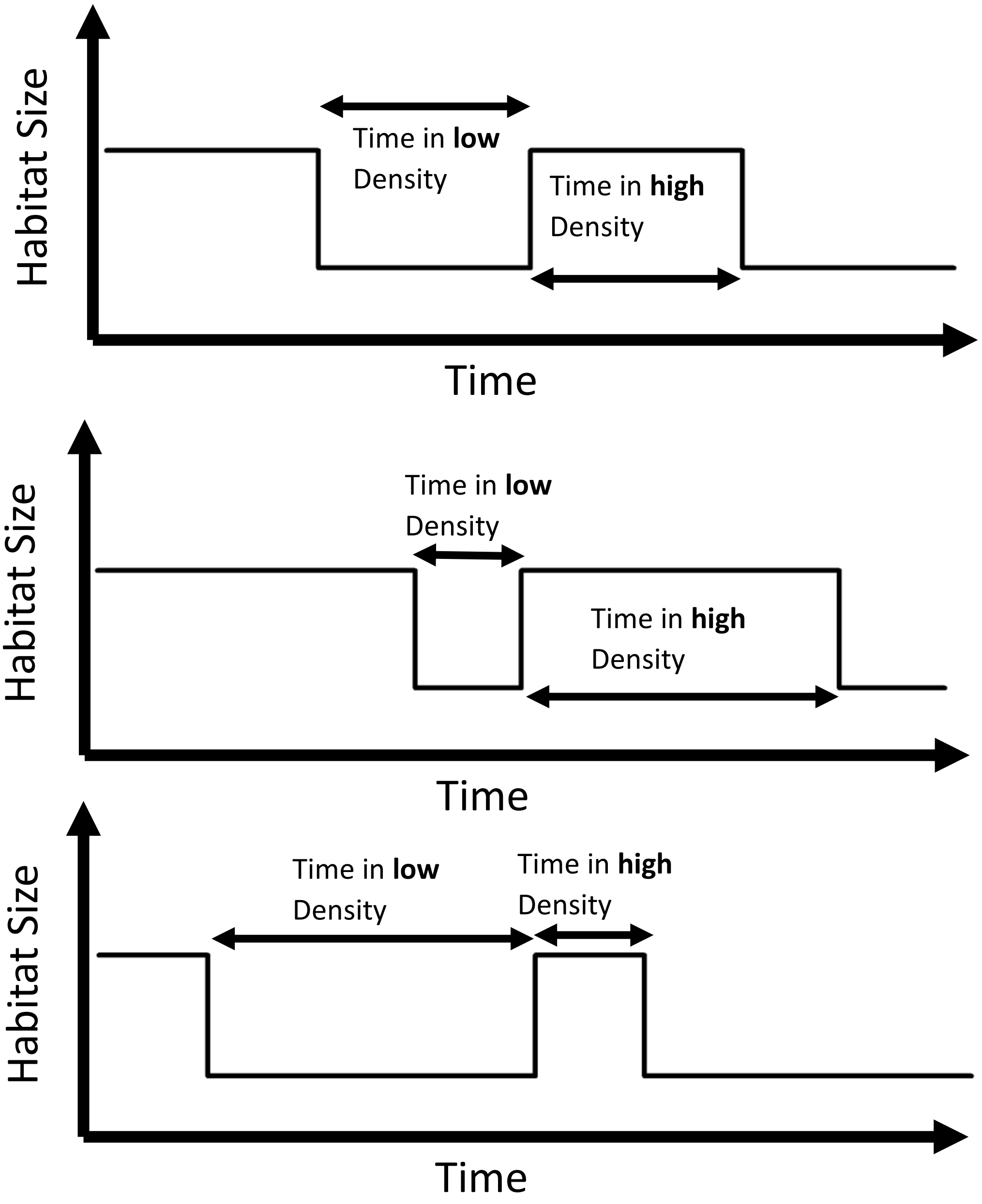


**Figure A: Exemplary illustrations on habitat size alterations for density fluctuations.**

### 2. Mathematical Approximation

We use a simplified mathematical model to calculate lifetime number of viable offspring, $\omega$, depending on waiting time $\tau$. We follow a similar approach as Tsitrone et al. (2003a) and focus on lifetime reproductive success in the female role, i.e. number of eggs that give rise to viable offspring, and assume that success in the male role is independent of waiting time. Although our simulation works in discrete time, we here assume continuous time because it simplifies the calculations. Individuals die at a rate *d* such that the time of death, $T_D$, is exponentially distributed with parameter *d*. Similarly, individuals meet mating partners at a rate

$$\boldsymbol{\Phi = 1 - \left(1 - \frac{1}{g^2}\right)^{N-1}}, \qquad \textbf{(1)}$$

where $g^2$ is the habitat size and *N* is the population size. This corresponds to the probability that at least one of the other *N*-1 individuals is found in the same grid cell as the focal individual on a particular day. The time to the first meeting, $T_M$, is then exponentially distributed with parameter $\Phi$. We draw $T_D$ and $T_M$ as two independent random variables. Meeting events that

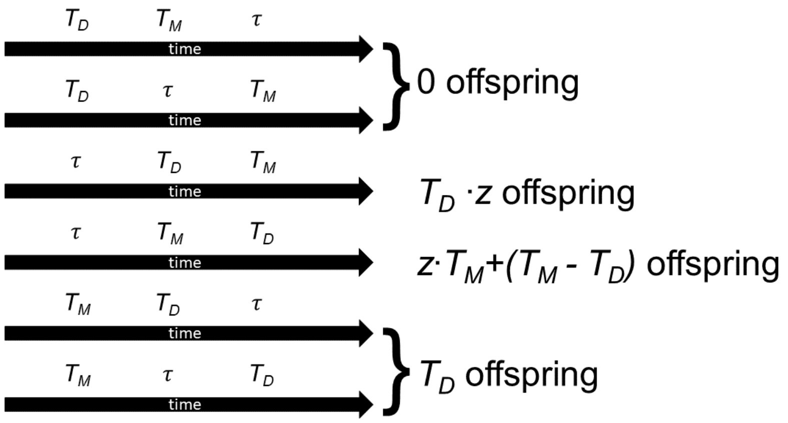


**Figure B: Possible life histories in the simplified model with their corresponding lifetime number of viable offspring. $T_D$ is the time of death, $T_M$ time to first meeting, $\tau$ is the waiting time, where an individual will start selfing, and *z* the viability of selfed offspring. In the first two cases, an individual dies before getting the chance to reproduce. The third case represents a case where an individual selfs at time $\tau$, and then dies before meeting a mating partner. All its offspring are affected by inbreeding depression, but it was able to lay all its eggs. In the fourth case, the individual started selfing after time $\tau$, but later met a mating partner. It therefore released all its eggs until time $T_M$ as selfed progeny that suffers from inbreeding depression, but then can release the rest as outcrossed eggs that are not affected by inbreeding depression. In the last two cases, the individual never selfed, because it met a mating partner early. It releases all of its eggs as outcrossed eggs without inbreeding depression.**

occur after death are of course discarded.

For given $T_D$, $T_M$ and $\tau$, we can then calculate lifetime number of viable offspring (Figure 2). Here we assume that individuals can use sperm they received for the rest of their lives. They will not self-fertilise after having outcrossed but will outcross even if they previously selfed. Individuals are able to produce one offspring per day, but results would be analogous with any other value since we are only interested in relative fitness. Resources can be fully stored until an opportunity to reproduce arises. Viability of selfed offspring $z$ was constant and did not depend on selfing proportion in the population.

To compute the expected number of viable offspring of an individual with waiting time $\tau$, we need to integrate over all possible times of death and mate encounter:

$$\boldsymbol{\omega(\tau) = \underbrace{\int_0^{\tau} f_d(l) \cdot \left\{ \int_0^{l} f_m(j) \cdot l \; dj \right\} dl}_{Case\ 5} + \int_{\tau}^{\infty} f_d(l) \left\{ \underbrace{\int_0^{\tau} f_m(j) \cdot l \, dj}_{Case\ 6} + \underbrace{\int_{\tau}^{l} f_m(j) \cdot (z \cdot j + l - j) \, dj}_{Case\ 4} + \underbrace{\int_{l}^{\infty} f_m(j) \cdot z \cdot l \, dj}_{Case\ 3} \right\} dl} \quad \textbf{(2)}$$

where $\boldsymbol{f_d(l)}$ is the probability density function of time to death and $\boldsymbol{f_m(l)}$ is the probability density function of meeting time. The case numbers refer to those in Fig. 12. Note that cases 1 and 2 do not have to be accounted for because they lead to 0 offspring. After substituting the parameters and integrating (using Mathematica), we obtain the fitness function

$$\boldsymbol{\omega(\tau) = \frac{\Phi(2+\frac{\Phi}{d}+e^{-(d+\Phi)\tau}(d \cdot z+\Phi(z-1))(1+\tau(d+\Phi))}{(d+\Phi)^2}} . \quad \textbf{(3)}$$

To predict the direction of selection, we then compute the derivative with respect to waiting time:

$$\boldsymbol{\omega'(\tau) = e^{-(d+\Phi)\tau} \cdot \big(\Phi - z(d+\Phi)\big)\tau.} \quad \textbf{(4)}$$

The direction of selection thus depends on the sign of $\Phi - z(d + \Phi)$, which is a constant under the assumptions of our mathematical approximation. If this term is positive, waiting times will increase towards infinity. If this term is negative, waiting times will evolve to approach zero.

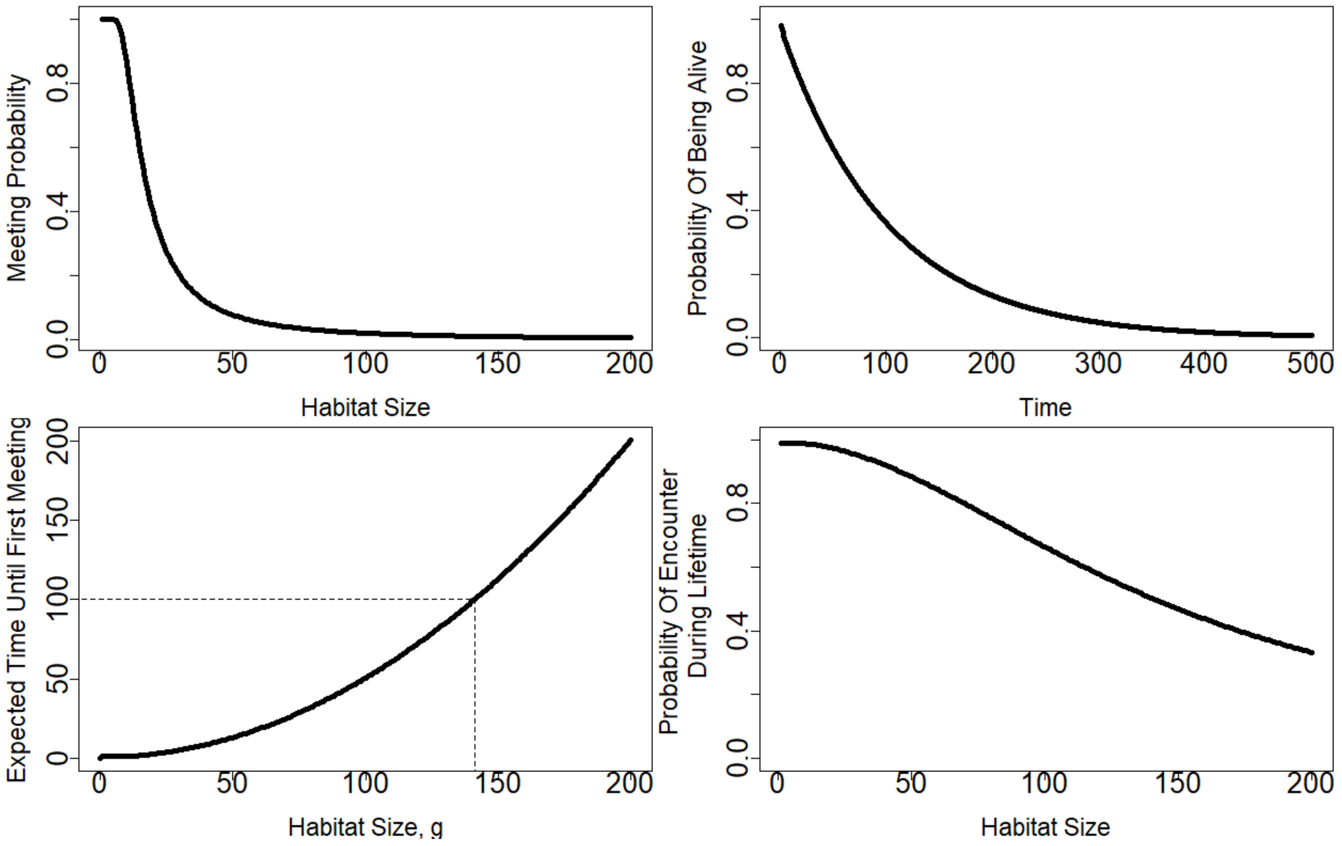


**Figure C: Plots showing basic measures such as A: The meeting probability per day $\Phi$, calculated by Formula (2), B: The probability to be alive after so many days, calculated by Formula (5) C: The expected time of first meeting as calculated by $E[T_m]=\frac{1}{\Phi(N,g)}$, indicated are average lifespan of 100 days and the habitat size at which expected time of first meeting equals average lifespan (*g*=141.07), and D: The combination of A and B: the probability for an individual to ever encounter a conspecific for reproduction within their lifetime as calculated by Formula (6).**

In Figure C we show expected probabilities for the habitat sizes used in the simulation. Meeting probability $\boldsymbol{\Phi}$ is calculated as in (1). And since individual lifespans are exponentially distributed, the probability to not have died at a certain time *t* is:

$$p_{alive} = e^{-d \cdot t}. \tag{5}$$

Because both time to first meeting and time of death are exponentially distributed, the probability for an individual to meet a mating partner within their lifetime is

$$p_{encounter\ during\ lifetime} = \frac{\Phi}{\Phi + d} \tag{6}$$

(see Pinsky and Karlin (2010) chapter 1.5.2 for proof).

### 3. Effects of Storage

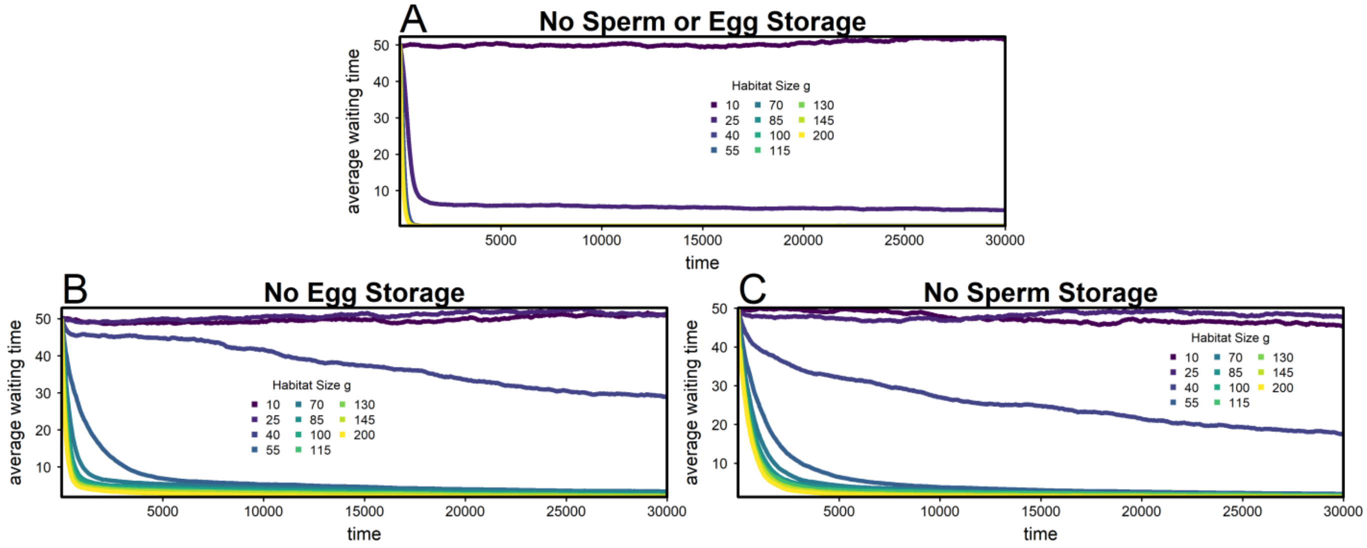


**Figure D: Simulation results without the ability to store resources for later usage. Number of replicates: 100. A: Sperm and eggs expired immediately on the day, after they were received/produced. B: Sperm lasted *o*=10 days, but eggs could not be stored. C: Eggs were stored, but sperm expired at the end of the day it was received.**

### 4. Long-term Behaviour

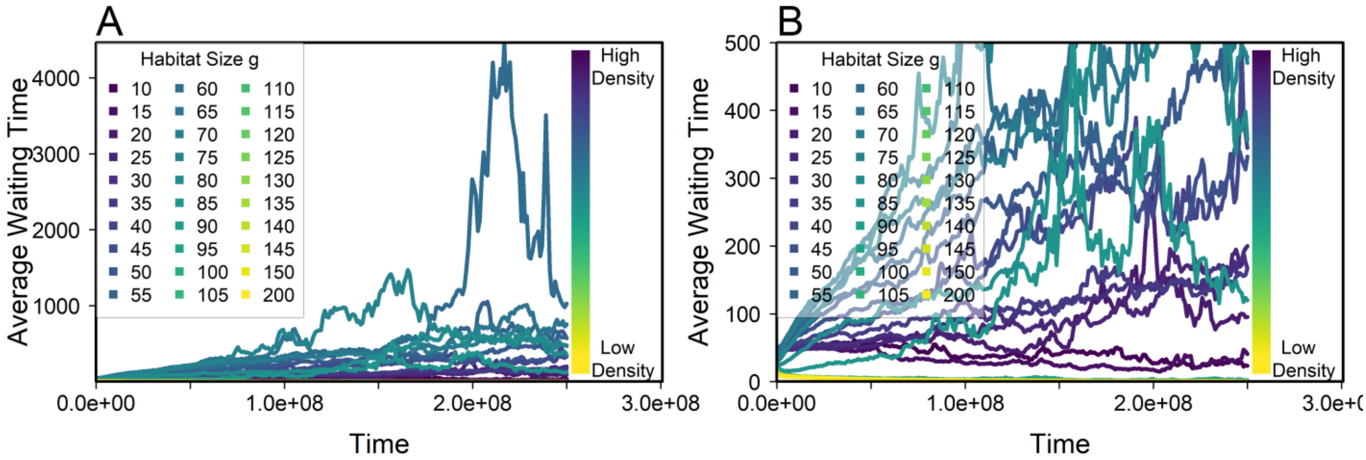


**Figure E: Waiting time evolution over a long time span. Simulation parameters equal to Figure 2, but with number of replicates=100. A: Automatically scaled y-axis. B: „zoomed“ y-axis with maximum 500.**

### 5. Purging and accumulation of deleterious alleles

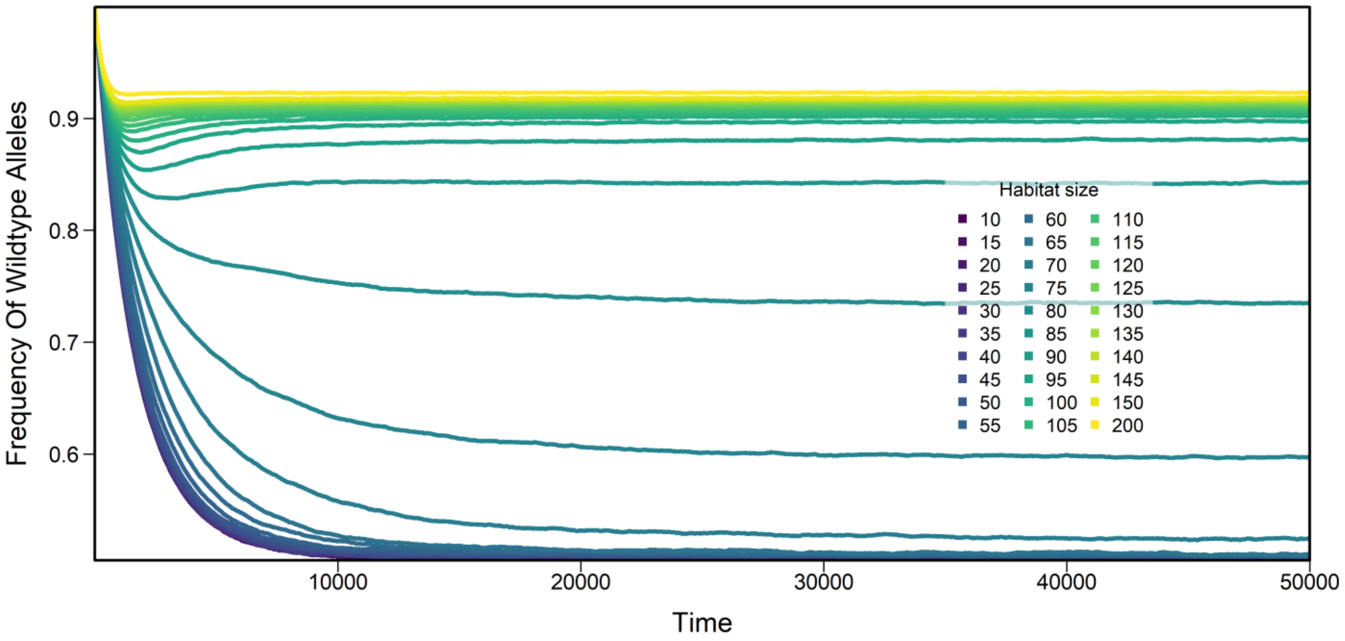


**Figure F: Frequency of wildtype alleles over time. Data from Figure 2.**

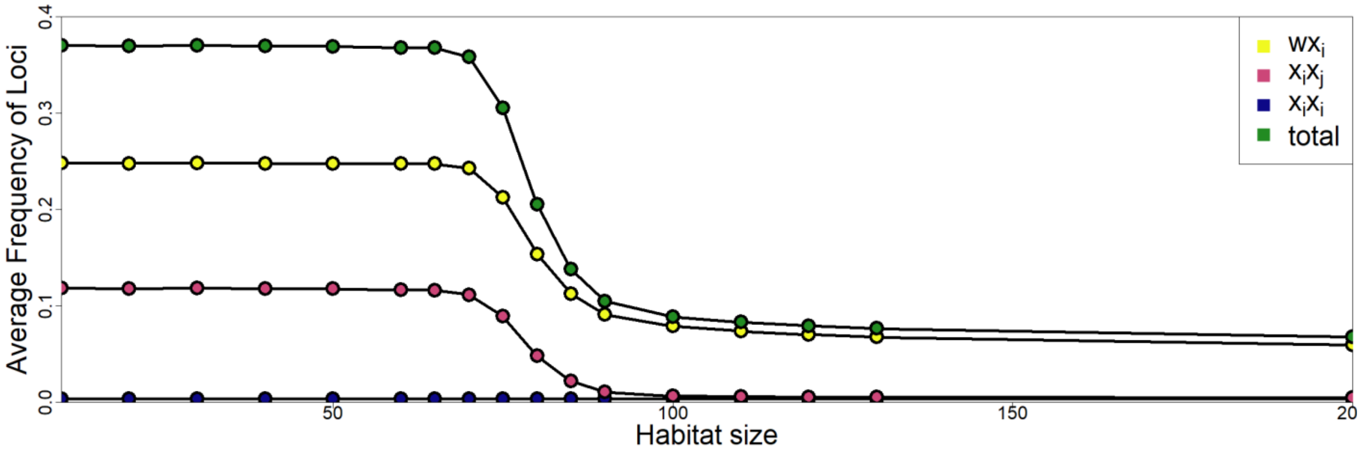


**Figure G: Proportion of loci being of each type (data from final 5% of simulation). The green "total" points indicate the frequency of all but the ww loci, therefore summarizing those that influence the calculation of o in Equation 3.**

### Individual Replicates

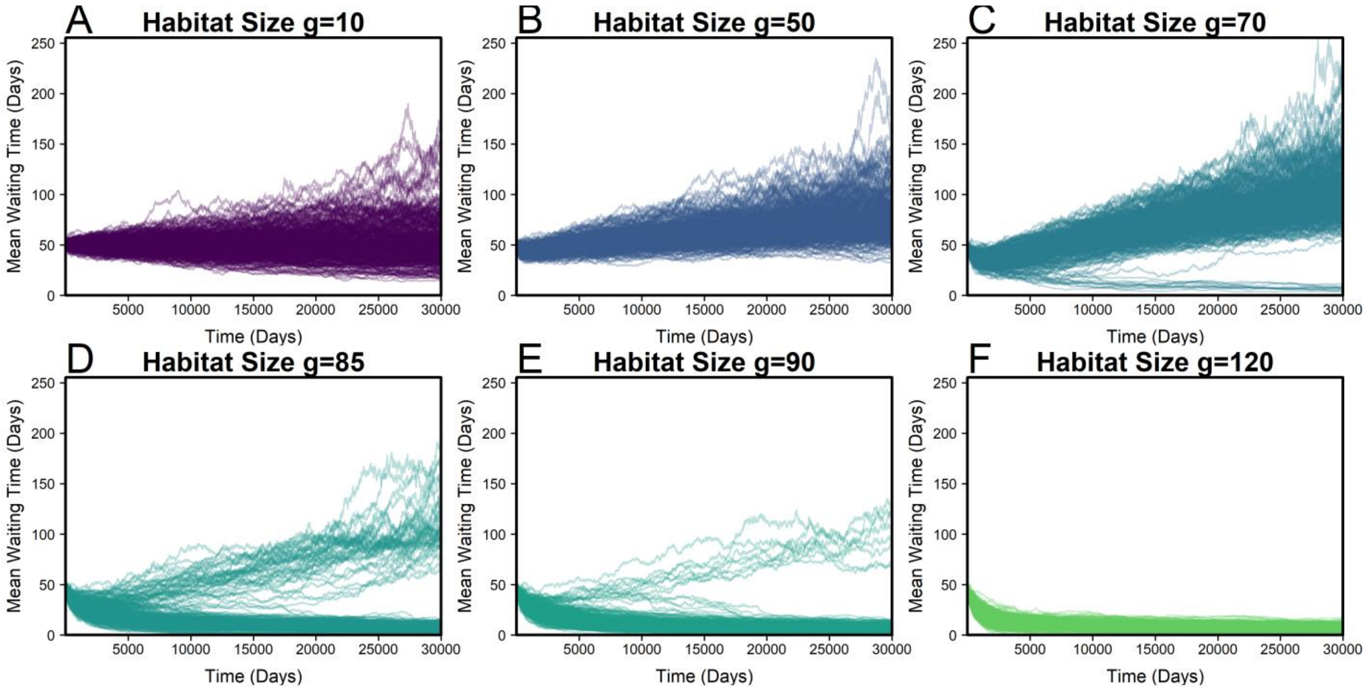


**Figure H: Trajectories for 200 replicates at different habitat sizes.**

### 6. Variation in Waiting Time

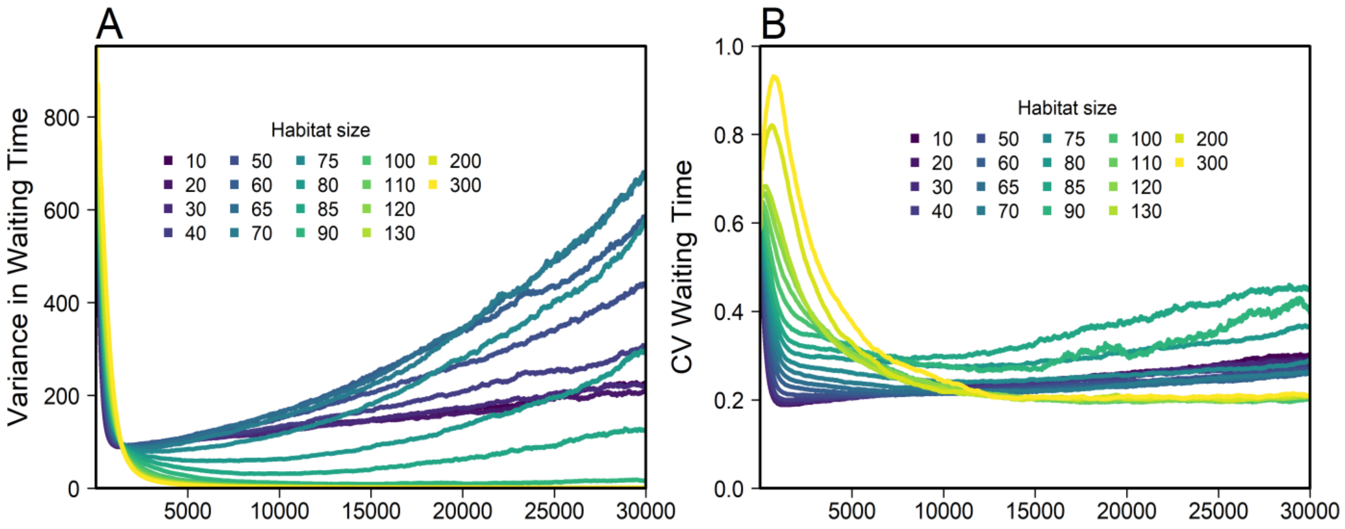


**Figure I: Temporal trajectories of variance and coefficient of variation of waiting time. Results as in Figure 2. A: Final (last day) within-replicate variance in waiting time, averaged over replicates, standard errors obscured by points. B: Final (last day) within-replicate coefficient of variation (CV) of waiting time, averaged over replicates with standard error.**

### 7. Population Size

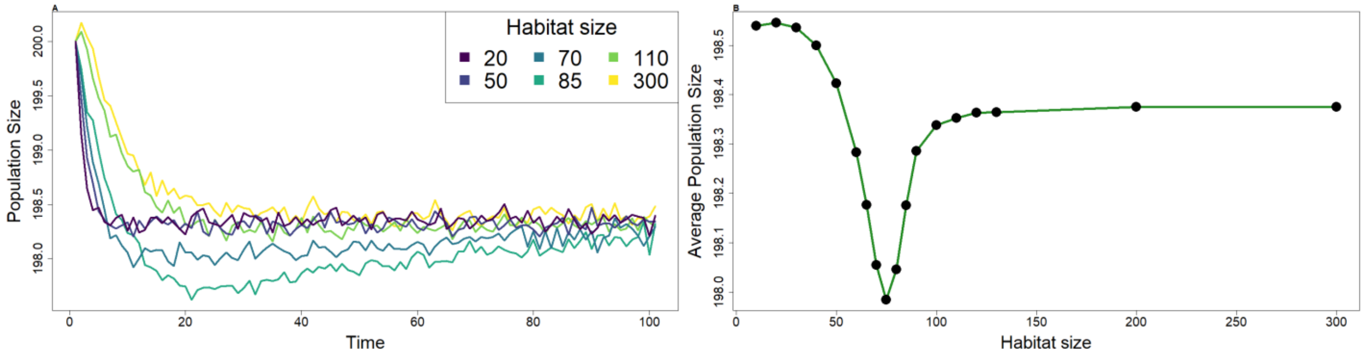


**Figure J: Consequences of waiting-time evolution for population size. Data as in Figure 2. A: Average population size over time, Data as in Figure 2, every 300th point is plotted to smooth lines. B: Average population size over the entire simulation over all (1000) replicates per habitat size *g*.**

### 8. Inbreeding depression necessary for outcrossing

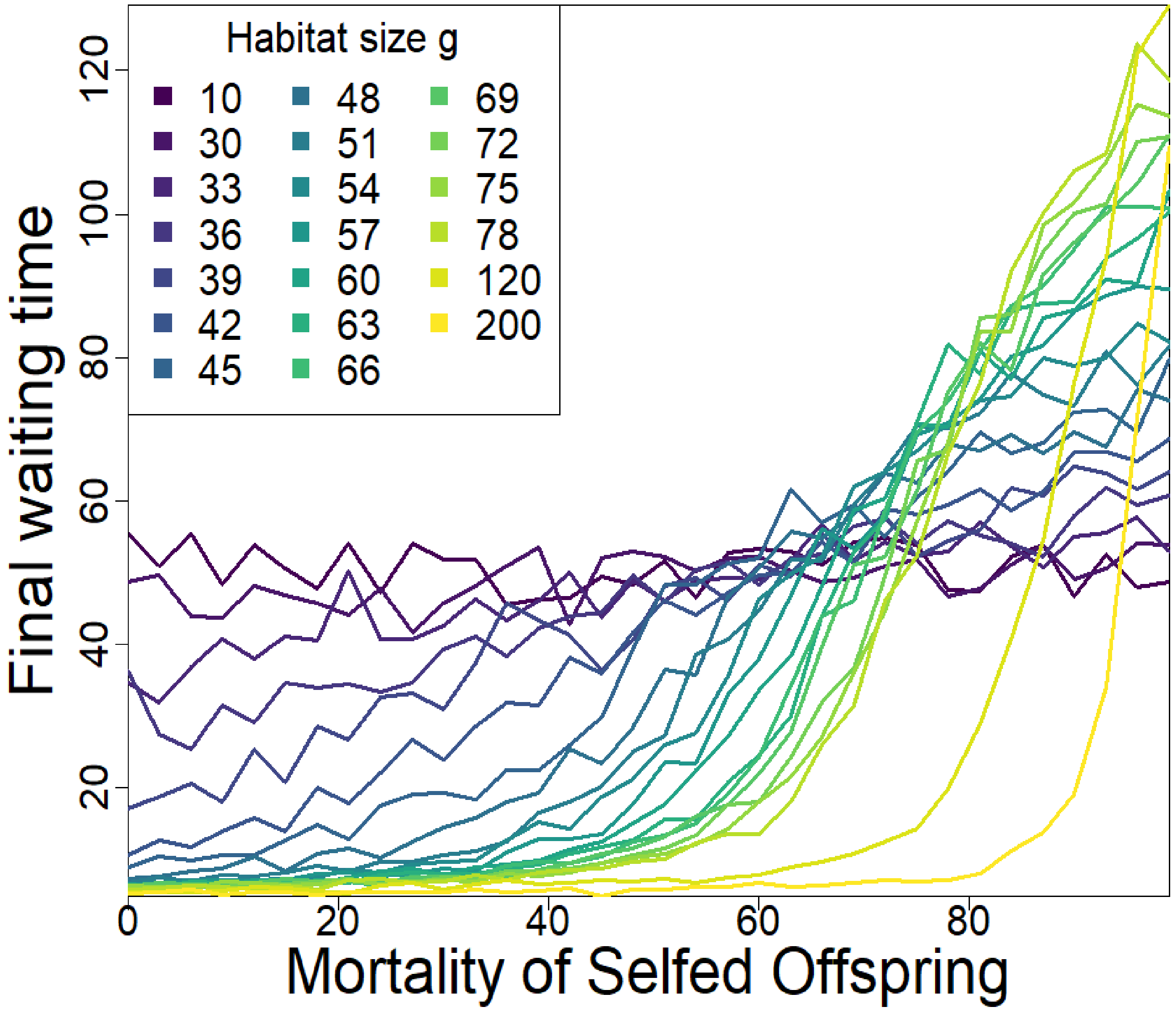


**Figure K: Average final (last day) waiting times with fixed mortality of selfed offspring after 30,000 days (number of days used in Figure 2). The dark (purple) lines indicate habitat size *g* where waiting time evolved due to drift alone. Therefore, the point where this line is crossed marks the minimum inbreeding depression needed to achieve an increase in waiting time within this number of days for a given density. Number of replicates: 50.**

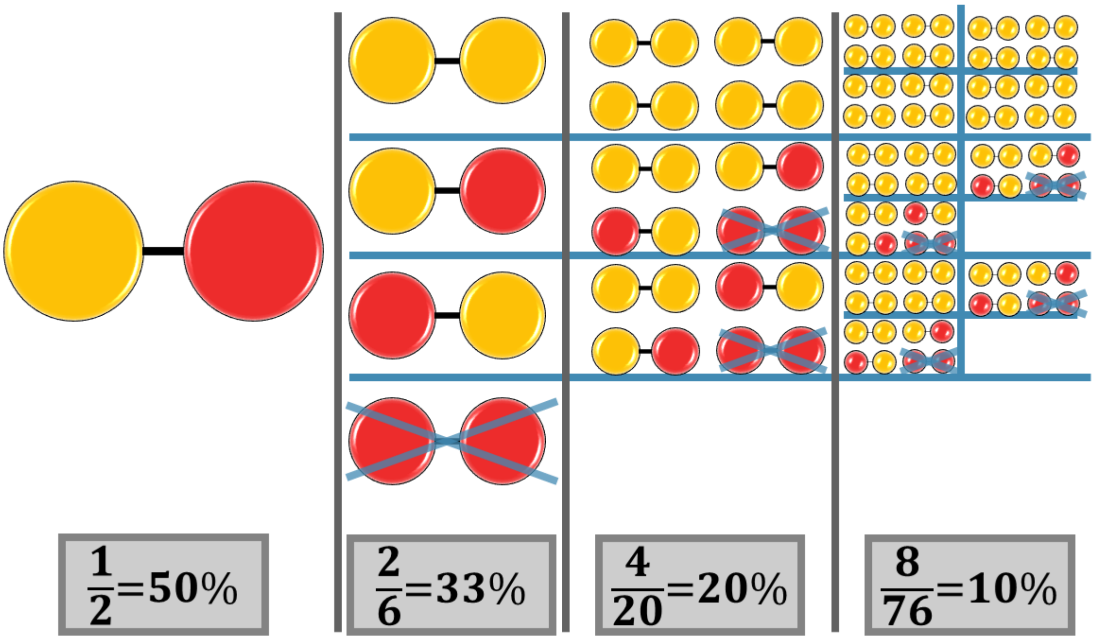


**Figure L: Example of purging of deleterious alleles when offspring is produced via selfing. The red circle represents a recessive lethal mutation, the yellow one the wild-type allele at one given locus. Vertical grey lines separate generations. Blue lines indicate which parent offspring belongs to. Crossed out loci indicate the individual has died, or will not reproduce due to being homozygous for the deleterious mutation. The grey boxes indicate proportion of alleles being deleterious in each generation.**

### 9. Allelic Diversity

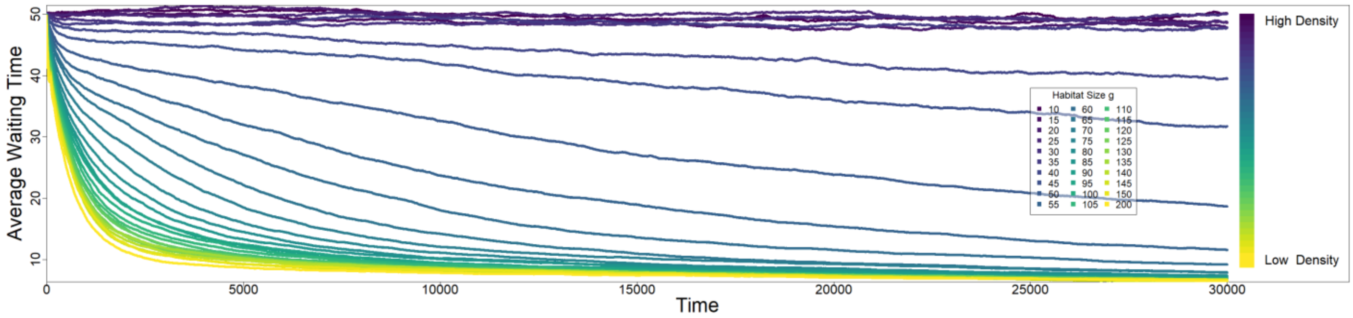


**Figure M: Waiting time evolution when allelic diversity *v*=2. Here we use only one possible type of deleterious mutation at the loci determining inbreeding depression. Colour bar added for clarity of how density relates to habitat size *g*. Number of replicates=1000.**

### 10. Robustness Analysis

In order to evaluate if any parameter had a strong influence on model results, we ran the model for each parameter being 5% larger or smaller than the default value shown in main text Table 1. We did this for the selection coefficient, sperm storage duration, the number of loci determining waiting time and inbreeding depression, respectively, as well as the corresponding mutation rates, the number of eggs produced per day, number of allele variants for inbreeding depression caring capacity and therefore number of individuals, and the death rate. Parameters with discrete values were manipulated to the closest value to 5% deviation from the default. Figure N and Figure O show the results under constant and fluctuating density, respectively.

No parameter drastically changed the pattern of the model output. All changes are within the expected direction.

Figure Q to Figure R show a different approach: to confirm that no specific parameter combination could lead to stable intermediate waiting times, we randomly drew all parameters from uniform distributions as shown in Table A. Discrete parameters were rounded to the nearest full number. We ran 7000 simulations with parameters drawn this way, 6% of which led to the extinction of the population. As a result, we got final waiting times and change in waiting time over the last iteration of the step function for 6577 parameter combinations.

We then chose parameter combinations where final waiting time was between 10% and 90% of the average lifespan of an individual ("intermediate waiting time"). Among those we chose the ones with the 10 smallest relative changes in waiting time within the last iteration of the step function. We compared the parameters for these replicates with all parameters (Figure P) and also manually looked at the parameter combinations. For 8 out of 10 replicates in this group, the mutation rate was lower than the default value ($u_w$ =0.05). Apparently, the populations therefore lost variation in waiting time, and then lacked the evolutionary potential to reach the endpoints by the end of the simulation. However, when we then replicated these 10 parameter combinations 100 times each, the replicates all appeared to reach different waiting times (Figure R). Then we ran 100 replicates each where after 30000 days all waiting time alleles in the population were either increased or decreased by 10% of the average lifespan. As seen in Figure Q, in no case there seems to be a systematic return to the previous value. In summary, these cases do not appear to represent stable intermediate waiting times but rather populations evolving very slowly because of low mutation rates.

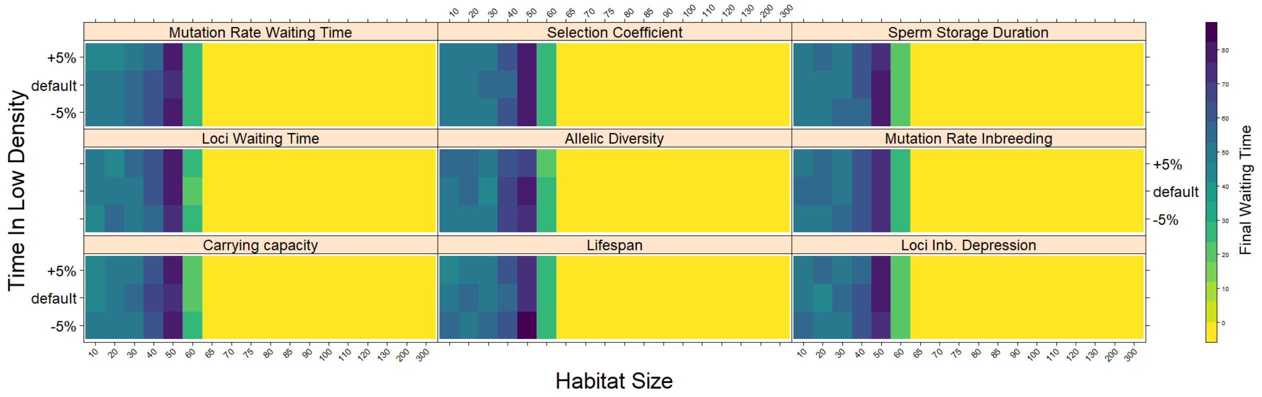


**Figure N: Robustness of the model with constant density. Each graph shows mean final waiting time over 100 replicates. Colors indicate final waiting time (over the last 5% of 30000 days). The parameters shown on the y-axis are altered by 5% or when discrete values were necessary the closest value to 5% deviation from the default. The most vulnerable densities are those with alternative stable states for waiting time**

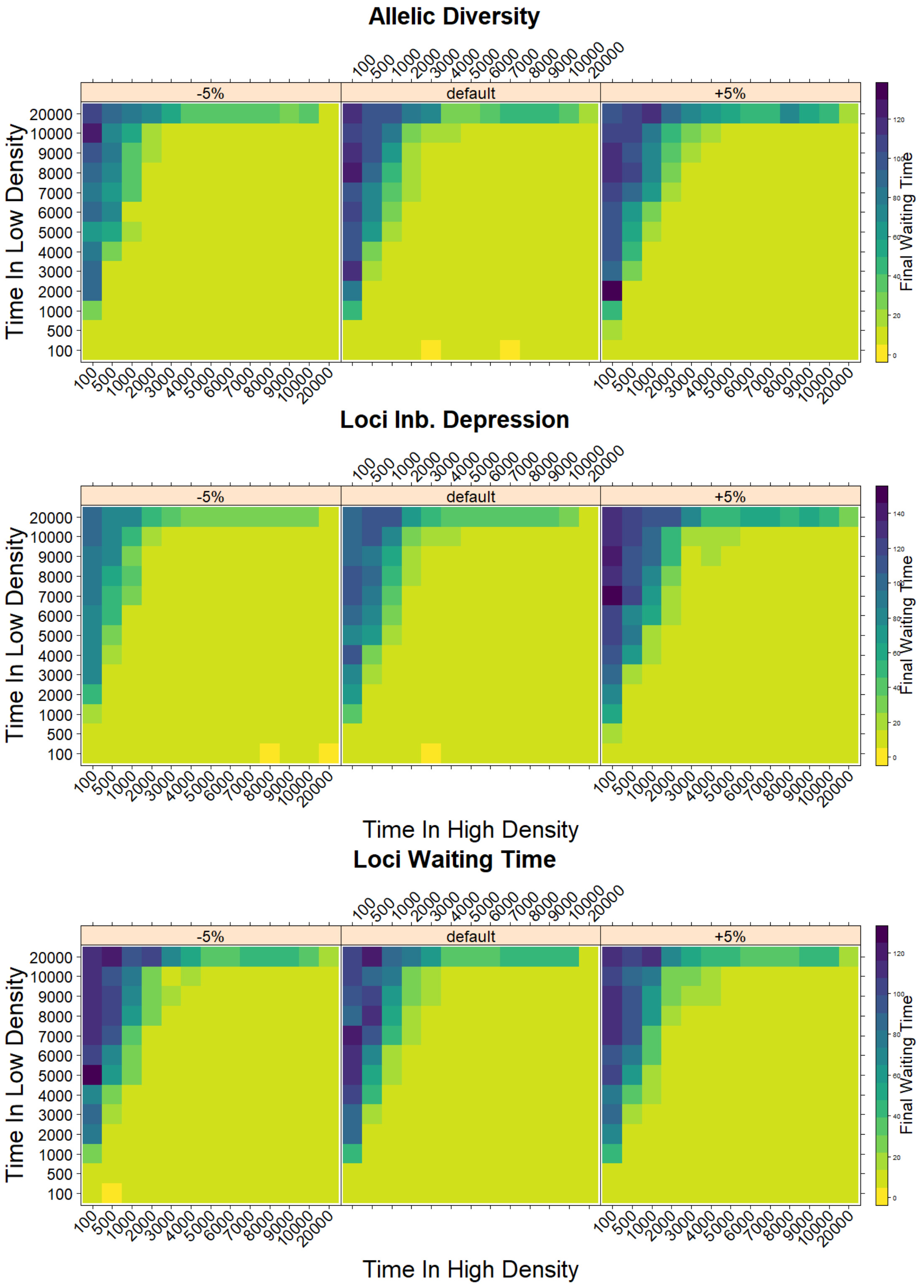


**Figure O: Robustness analysis with the method as in Figure N with fluctuating density. Colours indicate mean waiting time within the last 5% of 50000 days in the simulation. Each data point is averaged over 70 replicates. The default setting was simulated independently for each parameter. The default data was averaged and used in Figure 5.**

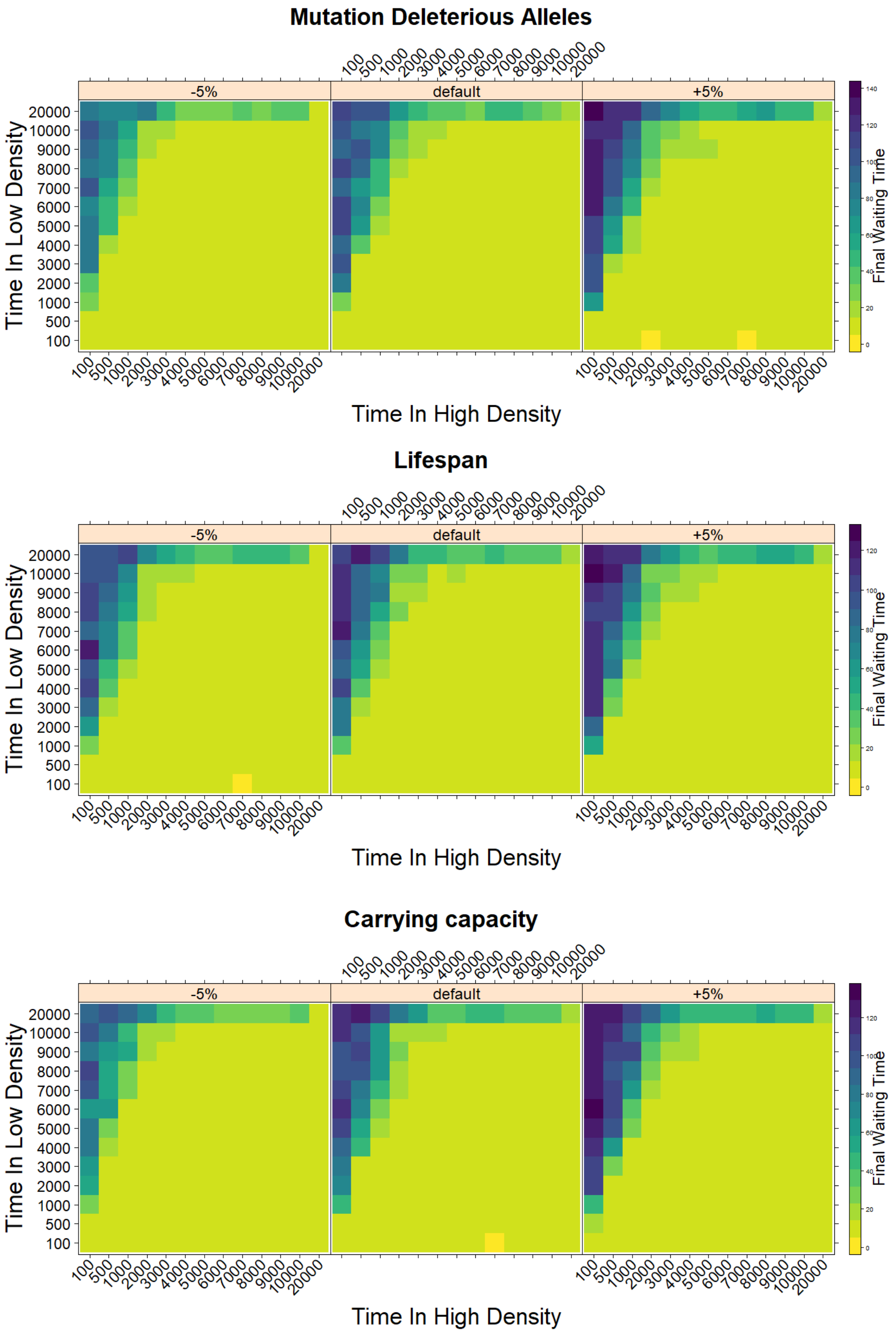


**Figure O continued**

### Selection Coefficient

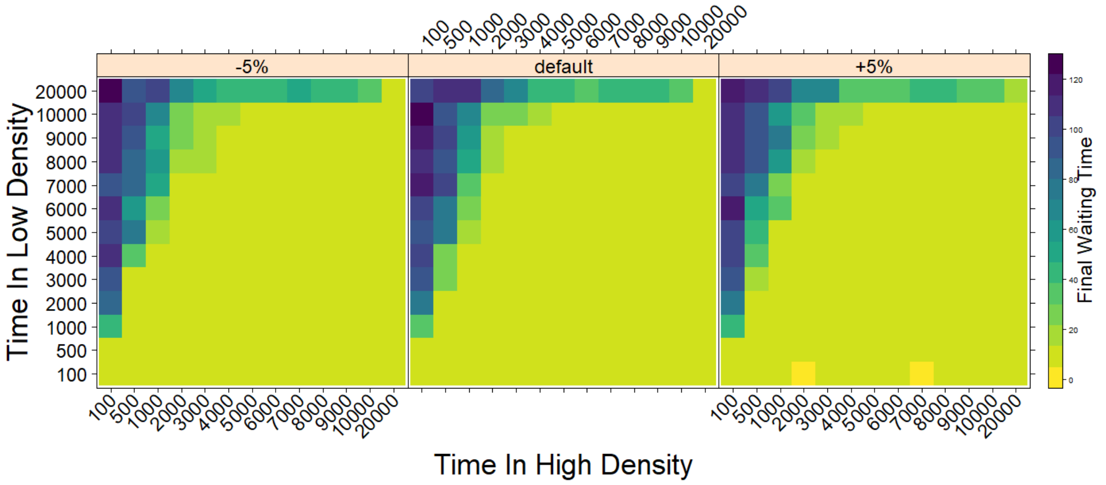


### Mutation Waiting Time

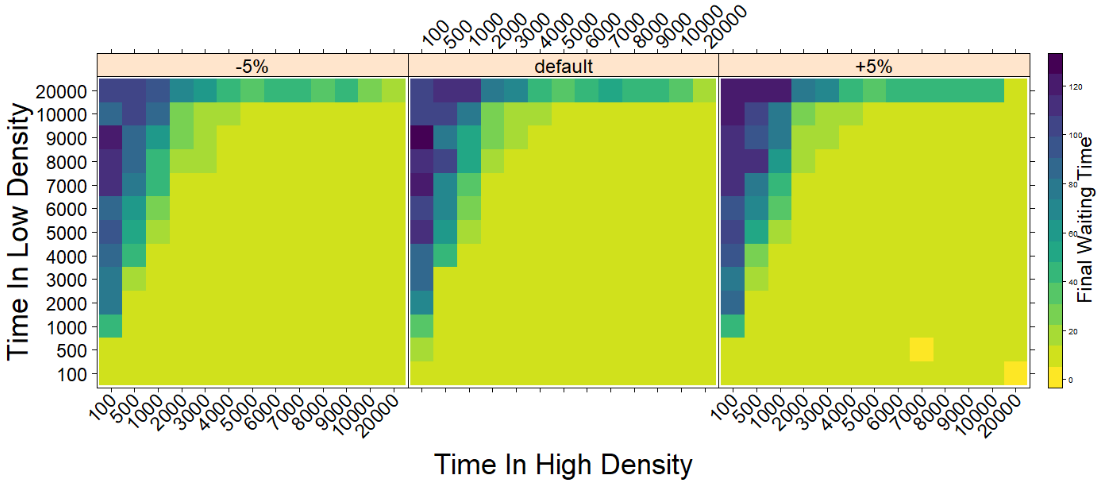


### Sperm Expiration Time

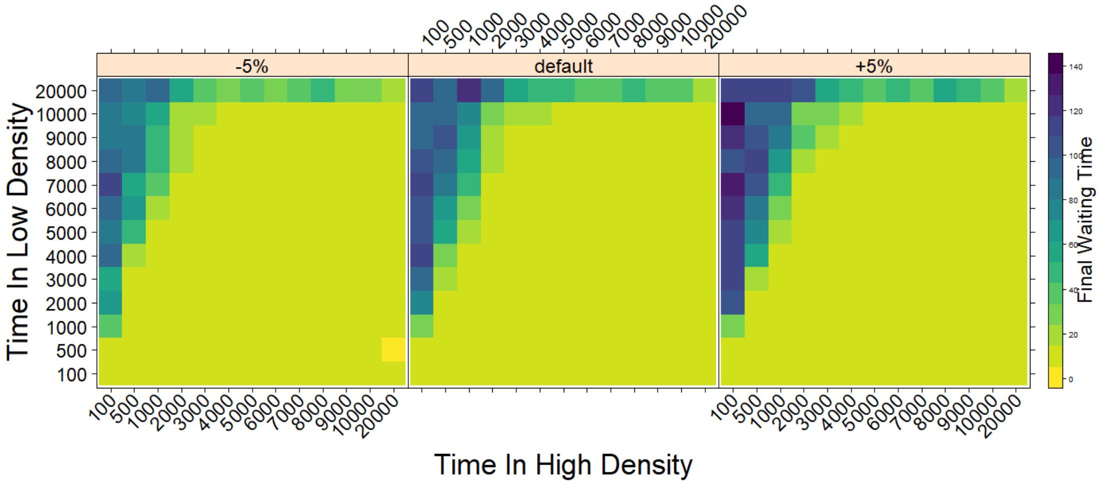


**Figure O continued**

**Table A: Ranges from which parameters were drawn from a uniform distribution.**

| Parameter | Symbol | Range |
|---|---|---|
| Average Lifespan | $d$ | 70-300 |
| Sperm storage duration | $o$ | 0-30 |
| Number of eggs produced per day | $c$ | 0.3-1.3 |
| Number of loci for waiting time | $W$ | 2-50 |
| Number of loci for inbreeding depression | $I$ | 2-50 |
| Standard deviation for mutation of waiting time loci | $u_w$ | 0.00005-0.01 |
| Mutation rate at inbreeding loci | $u_i$ | 0.00001-0.03 |
| "Allelic diversity" Number of alleles at inb. loci | $v$ | 3-300 |
| Selection coefficient | $s$ | 0-1 |
| Time in high density | | 50-12500 |
| Time in low density | | 50-12500 |

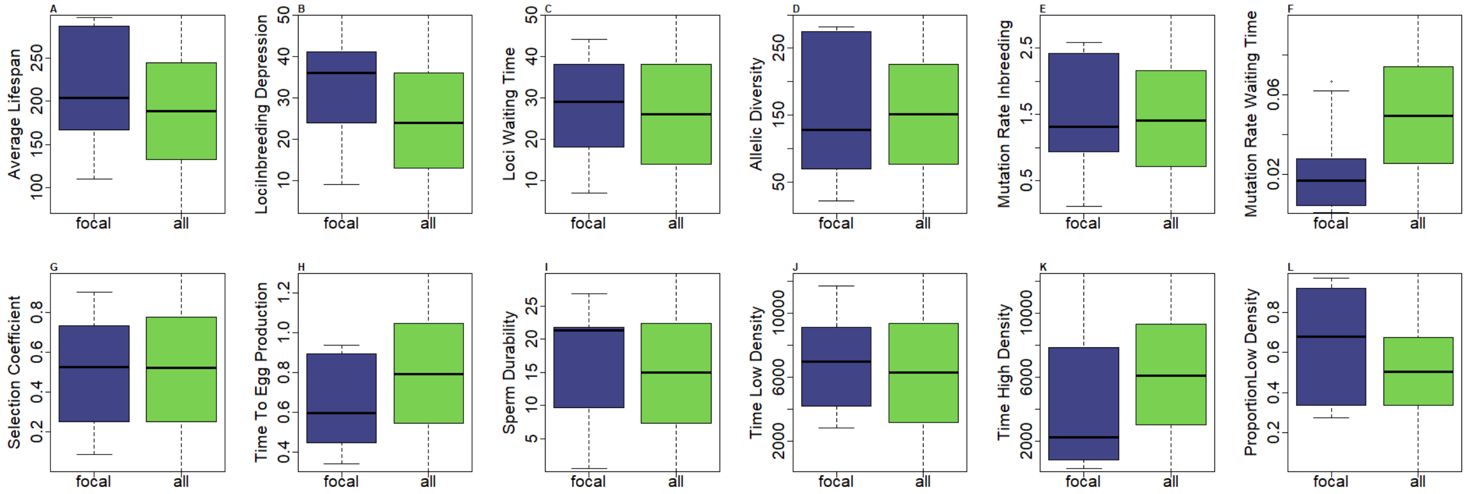


**Figure P: Comparison of random parameter values with those in the focal simulation. The right ("all") boxplot shows the distribution of randomly drawn parameter in 6577 simulations, while the left boxplot ("focal") shows the parameter distribution in 10 simulations, which we considered most likely to have evolved intermediate stable waiting time. To fulfil the first condition (intermediate final waiting time), the final waiting time (we used last day of the simulation, because different fluctuation patterns might have taken place during the final 5%) has to be between 0.1 and 0.9 of the average lifespan (which was randomly chosen for each simulation). For the second condition (stable) we searched for the 10 simulations with the smallest change in waiting time over the final iteration of the step function.**

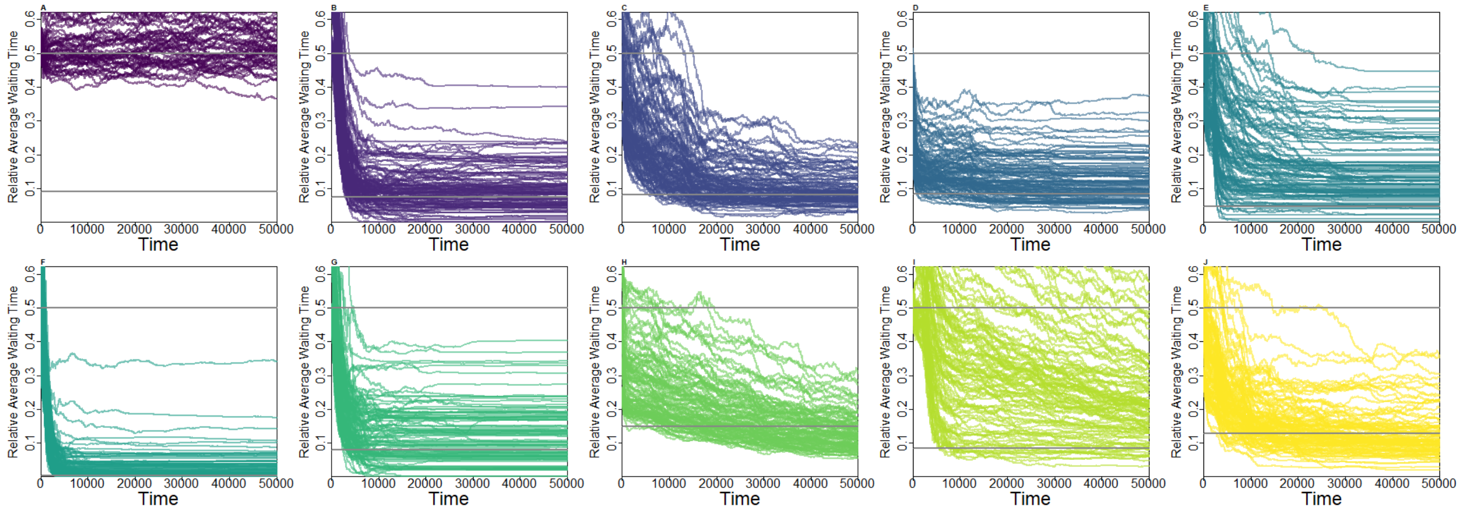


**Figure Q: Individual trajectories from 100 replicates with the parameters used in the focal populations chosen above. Gray lines indicate 50% of average lifespan and sperm expiration time.**

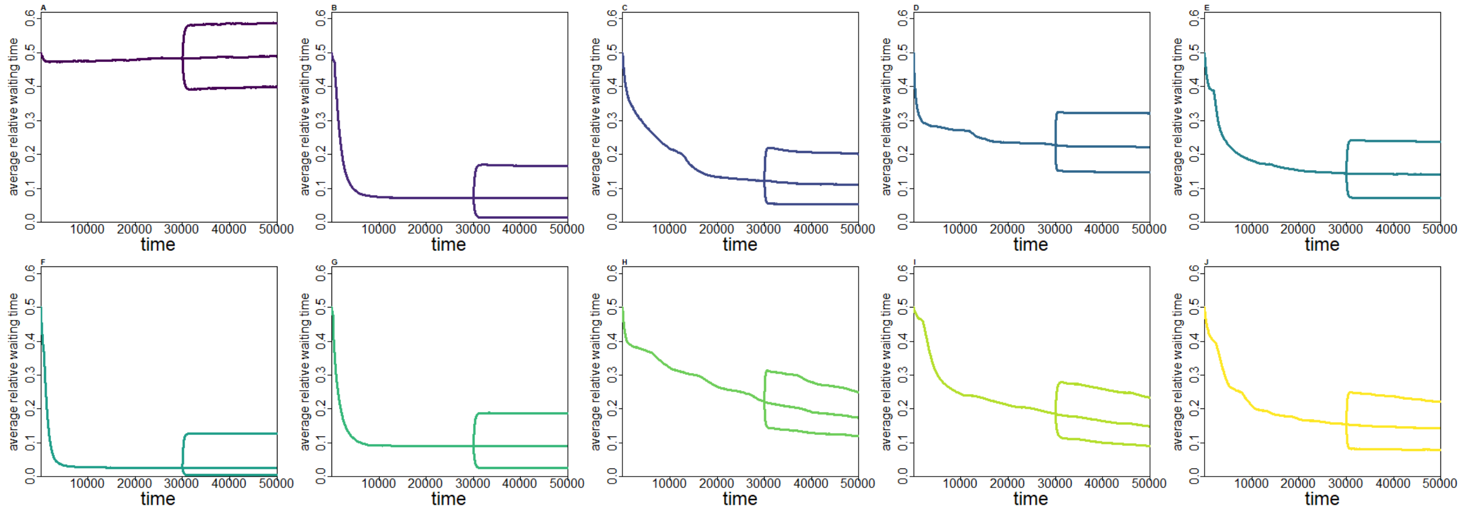


**Figure R: Averaged result from 100 replicates using the parameters from the focal populations and increasing or decreasing the waiting time alleles by 10% of average lifespan after 30000 days.**

### 11. Stochastic fluctuations

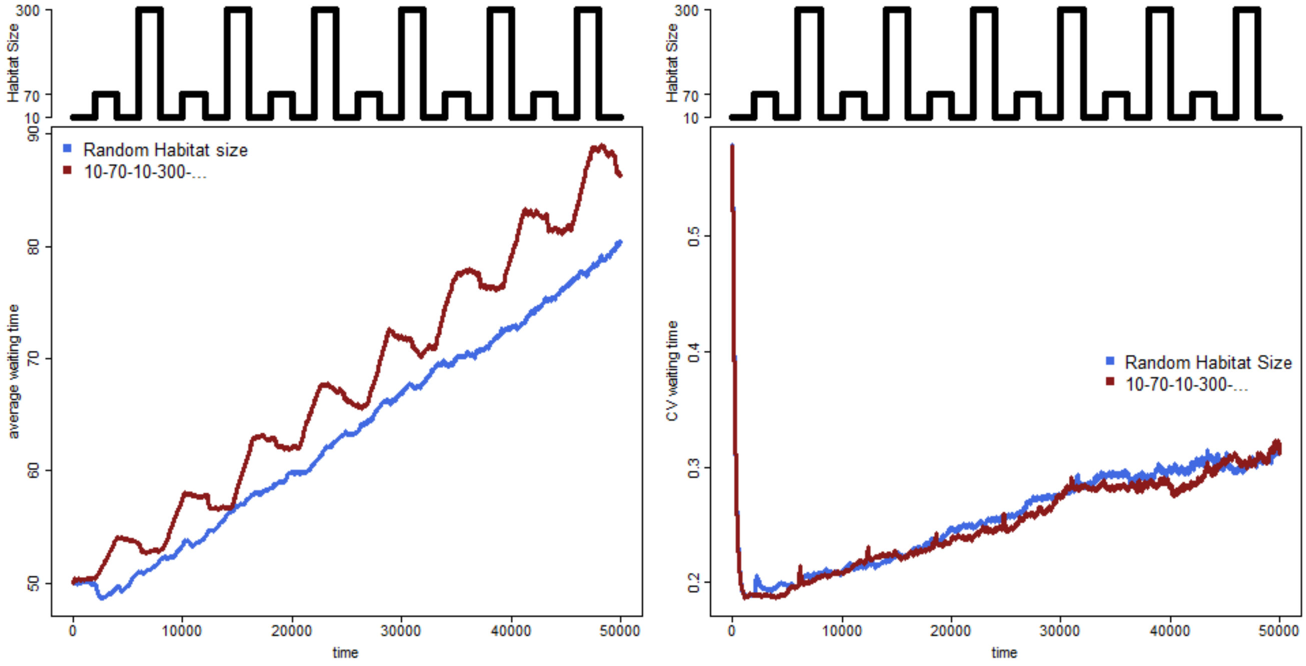


**Figure S: Alternating density with a pattern including 3 densities. The blue line represents random change between the three habitat sizes 10, 70, 300, lasting 2000 days after each change. For the red line, the pattern as shown above was used. The idea was that during high density phases (habitat size *g*=10) drift would increase variation and therefore increase evolutionary potential for the intermediate and low-density situation. Since this pattern was realistic for wild populations, we wanted to see whether it could increase variation in comparison to fluctuating density between two states. Strikingly, each time after a change in density towards low density (habitat size *g*=300) we see a sharp increase in coefficient of variation (CV). It is followed by a decrease in variation due to the strength of selection towards short waiting times. Number of replicates: 300.**
