## Supplementary material for "Waiting for love but not forever: modelling the evolution of waiting time to selfing in hermaphrodites": Simulation Scripts: AllFigures.R

```
#Plotting all the plots------------------
library(viridis)
library(lattice)
library(beepr)
setwd("D:/Arbeit/Flatworms/ThePaper/")
FigFol<-"Abb3/"
par(mfrow=c(1,2),mar=c(bottom=4.5,left=7,top=1.5,right=.4),cex.lab=3,cex.axis=2,mgp=c(3, .7, 0),xaxs='i',yaxs='i')

rel_legend_pos<-function(perc_x,perc_y){
 return(list(x=par("usr")[1]+diff(par("usr")[c(1,2)])*perc_x,y=par("usr")[3]+diff(par("usr")[c(3,4)])*perc_y)) 
}
#Fig1 Mathgraphs--------------
t<-seq(0,45,.001)
d<-.01
f<-.25
i<-.8
# g<-(f*(2+(f/d)+exp(-(d+f)*t)*(d*i+f*(i-1))*(1+t*(d+f))))/((d+f)^2)
# par(mfrow=c(1,2),mar=c(bottom=3,left=7,top=.45,right=.4),cex.lab=5,cex.axis=1.1,mgp=c(1.5, .5, 0))
# plot(t,g,type='l',lty=1,lwd=3,xlab=expression(tau),ylab=expression(paste(omega,"(",tau,")",sep="")))
# plot(t,exp(-(d+f)*t)*(f-i*(d+f))*t,type='l',lty=1,lwd=3,xlab=expression(tau),ylab=expression(paste(omega,"'(",tau,")",sep="")))
# idk<-diff(exp(-(d+f)*t)*(f-i*(d+f))*t)


t<-seq(0,500,.001)


fs<-seq(0.1,1,.1)/10
g<-matrix(nrow=length(fs),ncol=length(t))
g2<-g
for(f in fs){
  g[which(fs==f),]<-(f*(2+(f/d)+exp(-(d+f)*t)*(d*i+f*(i-1))*(1+t*(d+f))))/((d+f)^2)
  g2[which(fs==f),]<-exp(-(d+f)*t)*(f-i*(d+f))*t
}

{plotit<-function(){
legendcex<-7
lwds<-15
eeee<-.96
par(mfrow=c(1,2),mar=c(bottom=23,left=27,top=15,right=6),cex.lab=10,cex.axis=8,mgp=c(19, 2, 4),xaxs='i',yaxs='r',las=1)
matplot(t,t(g),type='l',lwd=lwds,lty=1,xlab=expression(tau),ylab=expression(paste(omega,"(",tau,")",sep="")),col=rev(viridis(length(fs),end=eeee)),xaxt='n',yaxt='n')
mtext("A",side=3,line=2,adj=0,cex=lwds)
box(lwd=10)
axis(side = 1, lwd =  lwds/3,tck=-0.02,mgp=c(15,8,0))
axis(side = 2, lwd =  lwds/3,tck=-0.02,mgp=c(12,5,0),las=2)
#legend(rel_legend_pos(0.6,0.7),as.character(fs),col=rev(viridis(length(fs))),pch=15,title="Meeting Probability",bty='n',cex=legendcex)
# print(par("usr"))
# print(c(par("usr")[1]+diff(par("usr")[c(1,2)])*.8,par("usr")[3]+diff(par("usr")[c(3,4)])*.4))
matplot(t,t(g2),type='l',lwd=lwds,lty=1,xlab=expression(tau),ylab=expression(paste(omega,"'(",tau,")",sep="")),col=rev(viridis(length(fs),end=eeee)),xaxt='n',yaxt='n')
mtext("B",side=3,line=2,adj=0,cex=lwds)
box(lwd=lwds/3*2)
axis(side = 1, lwd = lwds/3,tck=-0.02,mgp=c(15,8,0))
axis(side = 2, lwd =  lwds/3,tck=-0.02,mgp=c(12,5,0),las=2)
 legend(rel_legend_pos(0.6,0.7),as.character(fs),col=rev(viridis(length(fs),end=eeee)),pch=15,title="Meeting Probability",bty='n',cex=legendcex)
}
  
wid<-5000
hei<-2500
png(filename = paste0(FigFol,"F01_Mathgraphs.png"),width = wid, height = hei,pointsize=10);plotit();graphics.off()
pdf(file = paste0(FigFol,"F01_Mathgraphs.pdf"),width = wid, height = hei,pointsize=10);plotit();graphics.off()
}
#Fig2 F25-------------
load("FigureData/F25_ori_compressed.RData")
load("FigureData/F25_compressed.RData")
{plotit<-function(){
  legendcex<-7
  lwds<-15
  eeee<-.96
  col2<-adjustcolor("gray45",alpha.f=.65)
  par(mfrow=c(1,1),mar=c(bottom=23,left=27,top=3,right=6),cex.lab=10,cex.axis=8,mgp=c(19, 2, 4),xaxs='i',yaxs='i',las=1)
  matplot(waiter,col=viridis(dim(waiter)[2],end=eeee),type='l',lty=1,lwd=lwds,ylab="Average Waiting Time",xlab="Time",ylim=c(0,101),xlim=c(0,times+4000),xaxt='n',yaxt='n')
  lines(c(0,rel_legend_pos(0.9,.05)[[1]]-500),rep(10,2),col=col2,lwd=lwds/2)
  lines(c(0,rel_legend_pos(0.9,.05)[[1]]-500),rep(100,2),col=col2,lwd=lwds/2)
    box(lwd=lwds/3*2)
  axis(side = 1, lwd = lwds/3,tck=-0.02,mgp=c(15,8,0))
  axis(side = 2, lwd =  lwds/3,tck=-0.02,mgp=c(12,5,0),las=2)
  legend(rel_legend_pos(.05,.95),as.character(w1),cex=legendcex,col=viridis(dim(waiter)[2],end=eeee),pch=15,title=Faktor,ncol=4,bty='n')
range<-(max(waiter)*.95):(min(waiter)+(max(waiter)*.05))
points(rep(rel_legend_pos(0.9,.05)[[1]]+300,length(range)),seq(rel_legend_pos(0.9,.05)[[2]],rel_legend_pos(0.9,.95)[[2]],length.out=length(range)),col=rev(viridis(length(range),end=eeee)),pch=15,cex=10)
text(32300,max(range),"High \n Density",cex=legendcex)
text(32300,min(range),"Low  \n Density",cex=legendcex)


}
  wid<-5000
  hei<-2000
  png(filename = paste0(FigFol,"F02_F25.png"),width = wid, height = hei,pointsize=10);plotit();graphics.off()
  pdf(file = paste0(FigFol,"F02_F25.pdf"),width = wid, height = hei,pointsize=10);plotit();graphics.off()
}
#Fig3 DelA-----------------
load("D:/Arbeit/Flatworms/ThePaper/FigureData/delA_compressed.RData")
{plotit<-function(){
  legendcex<-7
  lwds<-15
  col2<-"gray53"
  par(mfrow=c(1,1),mar=c(bottom=23,left=27,top=3,right=6),cex.lab=10,cex.axis=8,mgp=c(19, 2, 4),xaxs='i',yaxs='i',las=1)
  
matplot(Wert1,defris,pch=21,lty=1,type='b',lwd=lwds,col="black",bg=c("firebrick2","royalblue"),xlab="Habitat Size",ylab="Frequency",ylim=c(0,1),cex=3.3,xaxt='n',yaxt='n')
box(lwd=lwds/3*2)
axis(side = 1, lwd = lwds/3,tck=-0.02,mgp=c(15,8,0))
axis(side = 2, lwd =  lwds/3,tck=-0.02,mgp=c(12,5,0),las=2)
for(i in 1:18){
  lines(rep(Wert1[i],2),rep(defris[i,2],2)+c(dsd[i],-dsd[i]),col=col2,lwd=lwds*.7)
  points(rep(Wert1[i],2),rep(defris[i,2],2)+c(dsd[i],-dsd[i]),cex=lwds*.7,pch='-',col=col2,lwd=lwds*.7)
  lines(rep(Wert1[i],2),rep(defris[i,1],2)+c(dsd2[i],-dsd2[i]),col=col2,lwd=lwds*.7)
  points(rep(Wert1[i],2),rep(defris[i,1],2)+c(dsd2[i],-dsd2[i]),cex=lwds*.7,pch='-',col=col2,lwd=lwds*.7)
}
matplot(Wert1,defris,pch=21,lty=1,type='b',lwd=5,col="black",bg=c("firebrick2","royalblue"),cex=lwds,add=T)
legend(rel_legend_pos(.7,.6),c("wildtype","deleterious"),col=c("royalblue","firebrick2"),pch=15,bg="white",cex=legendcex*2,bty='n')

}
  wid<-5000
  hei<-2000
  png(filename = paste0(FigFol,"F03_delA.png"),width = wid, height = hei,pointsize=10);plotit();graphics.off()
  pdf(file = paste0(FigFol,"F03_delA.pdf"),width = wid, height = hei,pointsize=10);plotit();graphics.off()
}
#Fig4 Multipanel------------
load("D:/Arbeit/Flatworms/ThePaper/FigureData/F25_compressed.RData")
h<-3 #h�he vier graphen
h2<- 3 #h�he quadrate
b1=9  #breite gesamt (muss durch drei teilbar sein)
g1<-matrix(1,nrow=h,ncol=b1)
g2<-g1+1
g3<-g2+1
g4<-g3+1
obereh�lfe<-cbind(g1,g2)
untereh�lfe<-matrix(ncol=b1*2,nrow=h2*3)
s<-1:(h2)
bab<-1:((b1/3)*2)
for(i in 5:13){
  # stop()
  untereh�lfe[s,bab]<-i
  bab<-bab+(b1/3)*2     
  if(i %in% c(7,10)){
    s<-s+h2
    bab<-1:((b1/3)*2)
  }}
png(file="D:/Arbeit/Flatworms/ThePaper/Abbildungen2/Multi.png",width=1200/1.7,height=1400/1.7)
# pdf(file="D:/Arbeit/Flatworms/ThePaper/Abbildungen2/Multi.pdf",width=1200/1.7,height=1400/1.7)
layout(rbind(obereh�lfe,untereh�lfe-2))
par(mar=c(bottom=3.1,left=6,top=1,right=1),cex.axis=2,mgp=c(2, .7, 0),cex.lab=2,yaxs='r',xaxs='r')
x<-1:times
cix=.8
colori<-viridis(dim(waiti_vari2)[1])
# matplot(x,t(waiti_vari2[,x]),lwd=3,type='l',lty=1,xlab="Time",ylab="Variance in Waiting Time",col=colori,xaxs="i",yaxs="i")
# legend(3000,810,as.character(w1),cex=cix,col=viridis(dim(waiti_vari2)[1]),pch=15,title=Faktor,ncol=5,bg="n",bty='n')
# matplot(x,t(waiti_CV2[,x]),lwd=3,type='l',lty=1,xlab="Time",ylab="CV Waiting Time",col=colori,xaxs="i",yaxs="i")
# legend(7500,.78,as.character(w1),cex=cix,col=colori,pch=15,title=Faktor,ncol=5,bg="n",bty='n')
bigi<-3
plot(w1,waiti_vari,lwd=2,type='b',pch=21,col='black',bg=colori,cex=1,lty=3,xlab=Faktor,ylab="Final Variance \n Waiting Time",ylim=c(min(c(rep(waiti_vari,2)+c(sd_vari,-sd_vari))),max(c(rep(waiti_vari,2)+c(sd_vari,-sd_vari)))))
title(main=LETTERS[1],adj=0)
for(i in 1:length(sd_vari)){
  lines(rep(w1[i],2),rep(waiti_vari[i],2)+c(sd_vari[i],-sd_vari[i]),lwd=2.5,col="grey50")
  points(rep(w1[i],2),rep(waiti_vari[i],2)+c(sd_vari[i],-sd_vari[i]),cex=1.7,pch='-',col="grey50")
}
points(w1,waiti_vari,pch=21,bg=colori,cex=bigi,col="black")

plot(w1,waiti_CV,lwd=2,type='b',pch=21,col='black',bg=colori,cex=1.7,lty=3,xlab=Faktor,ylab="Final CV \n Waiting Time",ylim=c(0,.5))
title(main=LETTERS[2],adj=0)
for(i in 1:length(sd_vari)){
  lines(rep(w1[i],2),rep(waiti_CV[i],2)+c(sd_CV[i],-sd_CV[i]),lwd=2.5,col="grey50")
  points(rep(w1[i],2),rep(waiti_CV[i],2)+c(sd_CV[i],-sd_CV[i]),cex=1.7,pch='-',col="grey50")
}
points(w1,waiti_CV,pch=21,col='black',bg=colori,cex=bigi)

vs<-c(10,50,60,70,75,80,85,90,120)

vv<-which(Wert1 %in% (vs))
W<-matrix(-1,dim(wiria)[2],dim(wiria)[3])
SW<-W
O<-W
SO<-W
for(x in 1:dim(wiria)[2]){
  for(n in 1:dim(wiria)[3]){
    p<-which(wiria[,x,n]!=-1)
    W[x,n]<-mean(wiria[p,x,n]*100)
    SW[x,n]<-sd(wiria[p,x,n]*100)
    O[x,n]<-mean(oria[p,x,n])
    SO[x,n]<-sd(oria[p,x,n])
  }}
# W<-W*100

for(i in vv){
  print(plot(W[i,],O[i,],type='p',pch=21,col='black',bg=viridis(length(Wert1))[i],xlab="Final Waiting Time",ylab="Final Risk",cex=3,xlim=c(min(W),max(W)),ylim=c(0,1)))
  symbols(W[i,],O[i,],rectangles=cbind(SW[i,]*2,SO[i,]*2),add=TRUE,inches=FALSE,bg="gray53",fg=NA)
    abline(h=.5,col="black")
    abline(v=100,col="black")
  print(matplot(W[i,],O[i,],type='p',pch=21,col='black',bg=viridis(length(Wert1))[i],cex=3,add=T))
  title(main=LETTERS[i+2],adj=0)
  text(200,.2,paste0("g=",Wert1[i]),cex=bigi)
}
graphics.off()
#Fig5 Fluctuating---------------------
load("D:/Arbeit/Flatworms/Ori/BT/WMUd.RData")
load("D:/Arbeit/Flatworms/Ori/BT/CVUd.RData")
load("D:/Arbeit/Flatworms/Ori/BT/WMSd.RData")
load("D:/Arbeit/Flatworms/Ori/BT/CVSd.RData")
load("D:/Arbeit/Flatworms/ThePaper/FigureData/quick_compressed.RData")
# load("D:/Arbeit/Flatworms/Ori/Q/compressed.RData")
f<-c(which(Wert1==70),which(Wert1==300))
library(lattice)
library(viridis)

times<-50000
Wert1<-c(100,500,seq(1000,10000,1000),20000)#low density time
Wert2<-c(100,500,seq(1000,10000,1000),20000)#high density time 

WM<-(WMUd*38+WMSd*32)/70
CV<-(CVUd*38+CVSd*32)/70

W2s<-c(20000,9000,7000,6000,5000,3000,500)
cc<-which(Wert2%in%W2s)
ocol<-c("firebrick3","slateblue2")
i<-which(Wert1==1000)

{plotit<-function(){
  legendcex<-7
  lwds<-15
  col2<-"gray53"
  par(mfrow=c(1,2),mar=c(bottom=23,left=27,top=15,right=6),cex.lab=10,cex.axis=8,mgp=c(19, 2, 4),xaxs='i',yaxs='i',las=1)
  
  
print(matplot(0:(times-1),WM[,i,cc],ylab="Average Waiting Time",type='l',lty=1,lwd=lwds,xlab="Time",col=rev(viridis(length(Wert2)))[cc],xaxt='n',yaxt='n'))
mtext("A",side=3,line=2,adj=0,cex=lwds)
box(lwd=lwds/3*2)
axis(side = 1, lwd = lwds/3,tck=-0.02,mgp=c(15,8,0))
axis(side = 2, lwd =  lwds/3,tck=-0.02,mgp=c(12,5,0),las=2)
print(matplot(waiter[,f],col=ocol,type='l',lty=3,lwd=lwds,ylab="Average Waiting Time",xlab="Time",add=T))
#legend(rel_legend_pos(.01,.91),title="Time In High Density",col=rev(viridis(length(Wert2)))[cc],legend=as.character(Wert2)[cc],ncol=3,pch=15,cex=legendcex,box.col=adjustcolor("white",.4),bg=adjustcolor("white",.4))
#legend(rel_legend_pos(.01,.995),c("High Density","Low Density"),cex=legendcex,col=ocol,pch=15,ncol=2,box.col=adjustcolor("white",.4),bg=adjustcolor("white",.4))

print(matplot(0:(times-1),CV[,i,cc],ylab="CV",xlab="Time",type='l',lty=1,lwd=lwds,col=rev(viridis(length(Wert2)))[cc],ylim=c(0,1),xaxt='n',yaxt='n'))
mtext("B",side=3,line=2,adj=0,cex=lwds)
box(lwd=lwds/3*2)
axis(side = 1, lwd = lwds/3,tck=-0.02,mgp=c(15,8,0))
axis(side = 2, lwd =  lwds/3,tck=-0.02,mgp=c(12,5,0),las=2)
print(matplot(t(waiti_CV2[f,]),col=ocol,type='l',lty=3,lwd=lwds,add=T))
legend(rel_legend_pos(.45,.8-(.995-.91)),title="Fluctuating \n Time In High Density",col=(viridis(length(Wert2)))[cc],legend=rev(as.character(Wert2)[cc]),ncol=3,cex=legendcex,pch=15,box.col=adjustcolor("white",.4),bg=adjustcolor("white",.4))
legend(rel_legend_pos(.45,.95),c("High Density","Low Density") ,title="Constant",cex=legendcex,col=ocol,pch=15,ncol=2,box.col=adjustcolor("white",.4),bg=adjustcolor("white",.4))

}
  wid<-5000
  hei<-2000
  png(filename = paste0(FigFol,"F05_FlucDen.png"),width = wid, height = hei,pointsize=10);plotit();graphics.off()
  pdf(file = paste0(FigFol,"F05_FlucDen.pdf"),width = wid, height = hei,pointsize=10);plotit();graphics.off()
}

#Fig6 Positive FeedbackLoops-----

#___________________----------------------------------------
#-----------------Supplement-------------------

#Fig12 General-----------

{plotit<-function(){
  legendcex<-7
  lwds<-15
  col2<-"gray53"
  par(mfrow=c(2,2),mar=c(bottom=23,left=32 ,top=15,right=6),cex.lab=10,cex.axis=8,mgp=c(19, 2, 4),xaxs='i',yaxs='i',las=1)
  
#layout(matrix(c(1,2,3,3), 2, 2, byrow = TRUE))
N<-200
g<-0:200
plot(g,1-(1-(1/(g^2)))^N,type='l',lwd=lwds,xlab="Habitat Size",ylab="Meeting Probability",xaxt='n',yaxt='n')
mtext("A",side=3,line=2,adj=0,cex=lwds)
box(lwd=lwds/3*2)
axis(side = 1, lwd = lwds/3,tck=-0.02,mgp=c(15,8,0))
axis(side = 2, lwd =  lwds/3,tck=-0.02,mgp=c(12,5,0),las=2)
print(matplot(t(waiti_CV2[f,]),col=ocol,type='l',lty=3,lwd=lwds,add=T))

dd<-1:500
plot(dd,1-pgeom(dd,.01),type='l',lwd=lwds,xlab="Time",ylab="Probability Of \n Being Alive",ylim=c(0,1),xaxt='n',yaxt='n')
mtext("B",side=3,line=2,adj=0,cex=lwds)
box(lwd=lwds/3*2)
axis(side = 1, lwd = lwds/3,tck=-0.02,mgp=c(15,8,0))
axis(side = 2, lwd =  lwds/3,tck=-0.02,mgp=c(12,5,0),las=2)

######## Code 1 - Graph for plotting expected time until meeting partner (y) vs. habitat size (x)

hab_size <- seq(0,200,1)

exp_meet <- 1/(1-(1-1/hab_size^2)^200)
exp_death <- 1/0.01
exp_mat <- 20

g_from_t <- function(t){ # function to calculate g backwards from expected time until first meeting
  sqrt(1/(1-(1-1/t)^(1/200)))
}

plot_in <- function(x,y){ # plotting dashed lines for a single point (x,y) (x and y should not be vectors)
  lines(c(-10,x,x),c(y,y,-10),lty=2,lwd=lwds)  
}

g_exp_death <- g_from_t(exp_death)
g_exp_mat <- g_from_t(exp_mat)

# plot(hab_size,exp_meet,type='l',xlab="habitat size, g",ylab="expected time until first meeting")
# plot_in(g_exp_death,exp_death)
# plot_in(g_exp_mat,exp_mat)

###### Code 2 - coefficient of variance vs. mean value
# this code uses two levels of sampling.
# * At the lower level, Nrep population trajectories are made.
# * for each of these, Nind individuals are allowed to drift around over time
# * at the end of the trajectory, the variance and mean across trait values of the individuals is calculated
# * once all Nrep populations have been run, the slope between the means and variances is calculated
# * This procedure is then repeated metarep times
# * This gives us metarep slopes, that are plotted in a histogram, the histogram, as expected is centered around zero
#   and both positive and negative relations occur.
z0 = 100
Nind = 30 # (not set to high, because we want some variation across the replicates)
Nt = 100 # ok, let's just say 100 time points
Nrep = 1e2 # many replicates should be better
metarep = 1e2 # how many times to repeat the overall sampling

fin_mean <- numeric(Nrep)
fin_var <- numeric(Nrep)
meta_slope <- numeric(metarep)
for(j in 1:metarep){
  for(i in 1:Nrep){
    fin_z <- z0 + rowSums(matrix(rnorm(Nind*Nt,0,1),nrow=Nind))
    fin_mean[i] <- mean(fin_z)
    fin_var[i] <- var(fin_z)
  }
  meta_slope[j] <- coef(lm(fin_var~fin_mean))['fin_mean']
}
# par(mfrow=c(1,2))
# plot(fin_mean,fin_var)
# abline(coef(lm(fin_var~fin_mean)))
# hist(meta_slope,n=30)


plot(hab_size,exp_meet,type='l',xlab="Habitat Size",ylab="Expected Time Until \n First Meeting",lwd=lwds,xaxt='n',yaxt='n')
mtext("C",side=3,line=2,adj=0,cex=lwds)
box(lwd=lwds/3*2)
axis(side = 1, lwd = lwds/3,tck=-0.02,mgp=c(15,8,0))
axis(side = 2, lwd =  lwds/3,tck=-0.02,mgp=c(12,5,0),las=2)
plot_in(g_exp_death,exp_death)

#plot_in(g_exp_mat,exp_mat)
pmeet<-1-(1-(1/(g^2)))^N
g<-1:200
d<-.01
# plot((1-(1-(1/(g^2)))^N)/(1-(1-(1/(g^2)))^N+(.01)),type='l',lwd=5,xlab="Habitat Size",ylab="Probability Encounter \n In Lifetime",ylim=c(0,1),col="black")
plot(pmeet/(pmeet+d),type='l',lwd=lwds,xlab="Habitat Size",ylab="Probability Encounter \n In Lifetime",ylim=c(0,1),col="black",xaxt='n',yaxt='n')
mtext("D",side=3,line=2,adj=0,cex=lwds)
box(lwd=lwds/3*2)
axis(side = 1, lwd = lwds/3,tck=-0.02,mgp=c(15,8,0))
axis(side = 2, lwd =  lwds/3,tck=-0.02,mgp=c(12,5,0),las=2)

}
wid<-5000
hei<-2000
png(filename = paste0(FigFol,"F09_General.png"),width = wid, height = hei,pointsize=10);plotit();graphics.off()
pdf(file = paste0(FigFol,"F09_General.pdf"),width = wid, height = hei,pointsize=10);plotit();graphics.off()
}
#Fig10 NoSotre---------------
# par(mar=c(bottom=5,left=6,top=1.5,right=.3),cex.axis=2,cex.lab=1.7,cex.main=2,mgp=c(3, 1, 0))
{plotit<-function(){
  legendcex<-7
  lwds<-15
  col2<-"gray53"
  par(mfrow=c(2,2),oma=c(bottom=1, left=1, top=0, right=3),mar=c(bottom=23,left=27,top=15,right=6),cex.lab=10,cex.axis=8,mgp=c(19, 2, 4),xaxs='i',yaxs='i',las=1,cex.main=lwds)
  layout(matrix(c(0,1,1,0,2,2,3,3), 2, 4, byrow = TRUE))
load("D:/Arbeit/Flatworms/ThePaper/FigureData/NoStoreBoth_compressed.RData")
print(matplot(waiter,col=viridis(dim(waiter)[2]),type='l',lty=1,lwd=lwds,ylab="average waiting time",xlab="time",main="No Sperm or Egg Storage",xaxt='n',yaxt='n'))
mtext("A",side=3,line=2,adj=0,cex=lwds)
box(lwd=lwds/3*2)
axis(side = 1, lwd = lwds/3,tck=-0.02,mgp=c(15,8,0))
axis(side = 2, lwd =  lwds/3,tck=-0.02,mgp=c(12,5,0),las=2)
legend(rel_legend_pos(.5,.8),as.character(w1),cex=legendcex,bty='n',col=viridis(dim(waiter)[2]),pch=15,title=Faktor,ncol=3,bg=adjustcolor("white",alpha.f=.4))
load("D:/Arbeit/Flatworms/ThePaper/FigureData/NoStoreEgg_compressed.RData")
print(matplot(waiter,col=viridis(dim(waiter)[2]),type='l',lty=1,lwd=lwds,ylab="average waiting time",xlab="time",main="No Egg Storage",xaxt='n',yaxt='n'))
mtext("B",side=3,line=2,adj=0,cex=lwds)
box(lwd=lwds/3*2)
axis(side = 1, lwd = lwds/3,tck=-0.02,mgp=c(15,8,0))
axis(side = 2, lwd =  lwds/3,tck=-0.02,mgp=c(12,5,0),las=2)
legend(rel_legend_pos(.3,.65),as.character(w1),cex=legendcex,bty='n',col=viridis(dim(waiter)[2]),pch=15,title=Faktor,ncol=3,bg=adjustcolor("white",alpha.f=.4))
load("D:/Arbeit/Flatworms/ThePaper/FigureData/NoStoreSperm_compressed.RData")
print(matplot(waiter,col=viridis(dim(waiter)[2]),type='l',lty=1,lwd=lwds,ylab="average waiting time",xlab="time",main="No Sperm Storage",xaxt='n',yaxt='n'))
mtext("C",side=3,line=2,adj=0,cex=lwds)
box(lwd=lwds/3*2)
axis(side = 1, lwd = lwds/3,tck=-0.02,mgp=c(15,8,0))
axis(side = 2, lwd =  lwds/3,tck=-0.02,mgp=c(12,5,0),las=2)
legend(rel_legend_pos(.7,.9),as.character(w1),cex=legendcex,bty='n',col=viridis(dim(waiter)[2]),pch=15,title=Faktor,ncol=3,bg=adjustcolor("white",alpha.f=.4))


}
wid<-5000
hei<-2000
png(filename = paste0(FigFol,"F10_Store.png"),width = wid, height = hei,pointsize=10);plotit();graphics.off()
pdf(file = paste0(FigFol,"F10_Store.pdf"),width = wid, height = hei,pointsize=10);plotit();graphics.off()
}
#Fig9 Long-------------

load("D:/Arbeit/Flatworms/ThePaper/FigureData/Long_compressed.RData")
xx<-seq(500,times*500,length.out=dim(waiter)[1]-1)
#par(mfrow=c(1,2),mar=c(bottom=5,left=5,top=.45,right=.4),cex.lab=2,cex.axis=1.4,mgp=c(3.6, 2, 0))
{plotit<-function(){
  legendcex<-7
  lwds<-15
  col2<-"gray53"
  par(mfrow=c(1,2),mar=c(bottom=23,left=29,top=15,right=6),cex.lab=10,cex.axis=8,mgp=c(21, 2, 4),xaxs='i',yaxs='i',las=1,cex.main=lwds)
  
print(matplot(xx,waiter[-dim(waiter)[1],],col=viridis(dim(waiter)[2]),type='l',lty=1,lwd=lwds,ylab="Average Waiting Time",xlab="Time",xlim=c(0,max(xx)+51000000),xaxt='n',yaxt='n'))
mtext("A",side=3,line=2,adj=0,cex=lwds)
box(lwd=lwds/3*2)
axis(side = 1, lwd = lwds/3,tck=-0.02,mgp=c(15,8,0))
axis(side = 2, lwd =  lwds/3,tck=-0.02,mgp=c(12,5,0),las=2)
legend("topleft",as.character(w1),cex=legendcex,col=viridis(dim(waiter)[2]),pch=15,title=Faktor,ncol=3,bg=adjustcolor("white",alpha.f=.4))
range<-(max(waiter)*.95):(min(waiter)+(max(waiter)*.05))
points(rep(max(xx)*1.03,length(range)),range,col=viridis(length(range)),pch=15,cex=lwds)
text(max(xx)*1.12,max(range)*.98,"High \nDensity",cex=legendcex)
text(max(xx)*1.12,min(range)+max(range)*.02,"Low  \nDensity",cex=legendcex)
print(matplot(xx,waiter[-dim(waiter)[1],],col=viridis(dim(waiter)[2]),type='l',lty=1,lwd=lwds,ylab="Average Waiting Time",xlab="Time",xlim=c(0,max(xx)+51000000),ylim=c(0,500),xaxt='n',yaxt='n'))
mtext("B",side=3,line=2,adj=0,cex=lwds)
box(lwd=lwds/3*2)
axis(side = 1, lwd = lwds/3,tck=-0.02,mgp=c(15,8,0))
axis(side = 2, lwd =  lwds/3,tck=-0.02,mgp=c(12,5,0),las=2)
legend("topleft",as.character(w1),cex=legendcex,col=viridis(dim(waiter)[2]),pch=15,title=Faktor,ncol=3,bg=adjustcolor("white",alpha.f=.4))
range<-(500*.95):(0+500*.05)
points(rep(max(xx)*1.03,length(range)),range,col=viridis(length(range)),pch=15,cex=lwds)
text(max(xx)*1.12,max(range)*.98,"High \nDensity",cex=legendcex)
text(max(xx)*1.12,min(range)+max(range)*.02,"Low  \nDensity",cex=legendcex)
}
wid<-5000
hei<-1700
png(filename = paste0(FigFol,"F11_Longterm.png"),width = wid, height = hei,pointsize=10);plotit();graphics.off()
pdf(file = paste0(FigFol,"F11_Longterm.pdf"),width = wid, height = hei,pointsize=10);plotit();graphics.off()
}

#Fig18 Frequi----------
load("D:/Arbeit/Flatworms/ThePaper/FigureData/Frequi_compressed.RData")
{plotit<-function(){
  legendcex<-7
  lwds<-15
  col2<-"gray53"
  par(mfrow=c(1,1),oma=c(bottom=1, left=1, top=0, right=3),mar=c(bottom=23,left=29,top=15,right=12),cex.lab=10,cex.axis=8,mgp=c(21, 2, 4),xaxs='i',yaxs='i',las=1,cex.main=lwds)
  matplot(fre[,1,],type='l',lty=1,lwd=lwds,xlab="Time",ylab="Frequency Of Wildtype Alleles",col=viridis(length(Wert1)),xaxt='n',yaxt='n')
  box(lwd=lwds/3*2)
  axis(side = 1, lwd = lwds/3,tck=-0.02,mgp=c(15,8,0))
  axis(side = 2, lwd =  lwds/3,tck=-0.02,mgp=c(12,5,0),las=2)
  legend(rel_legend_pos(.7,.7),as.character(Wert1),ncol=3,title=Faktor,pch=15,col=viridis(length(Wert1)),bg=adjustcolor("white",.6),box.col=adjustcolor("white",.6),cex=legendcex)
}
wid<-5000
hei<-2500
png(filename = paste0(FigFol,"F12_Wildtype.png"),width = wid, height = hei,pointsize=10);plotit();graphics.off()
pdf(file = paste0(FigFol,"F12_Wildtype.pdf"),width = wid, height = hei,pointsize=10);plotit();graphics.off()
}
#Fig19 AvFreq---------
load("D:/Arbeit/Flatworms/ThePaper/FigureData/F25_compressed.RData")
 vv<-which(Wert1 %in% (c(10,50,70,85,90,120)))
{plotit<-function(){
  legendcex<-7
  lwds<-15
  col2<-"gray53"
  par(mfrow=c(1,1),mar=c(bottom=23,left=29,top=15,right=6),cex.lab=10,cex.axis=8,mgp=c(21, 2, 4),xaxs='i',yaxs='i',las=1,cex.main=lwds)
 
   matplot(Wert1,t(erg2/40/replikate),col='black',type='b',pch=21,bg=plasma(3),xlab=Faktor,ylab="Average Frequency of Loci",cex=lwds,lwd=lwds,lty=1,ylim=c(0,.4),xaxt='n',yaxt='n')
   box(lwd=lwds/3*2)
   axis(side = 1, lwd = lwds/3,tck=-0.02,mgp=c(15,8,0))
   axis(side = 2, lwd =  lwds/3,tck=-0.02,mgp=c(12,5,0),las=2)
   legend("topright",c(rev(expression(paste('x'[i],'x'[i],sep=''),paste('x'[i],'x'[j],sep=''),paste('w','x'[i],sep=''))),'total'),pch=15,col=c(rev(plasma(3)),"forestgreen"),cex=legendcex*2,bty='n')
lines(Wert1,colSums(erg2/40/replikate),type='b',pch=21,col='black',bg="forestgreen",cex=lwds,lwd=lwds)

}
wid<-5000
hei<-2500
png(filename = paste0(FigFol,"F13_Types.png"),width = wid, height = hei,pointsize=10);plotit();graphics.off()
pdf(file = paste0(FigFol,"F13_Types.pdf"),width = wid, height = hei,pointsize=10);plotit();graphics.off()
}

 #Fig13 Individual Replicates----------------
 load("D:/Arbeit/Flatworms/ThePaper/FigureData/Traj.RData")
 vv<-which(Wert1 %in% (c(10,50,70,85,90,120)))
 trajs<-300
 hi<-matrix(NA,length(vv),trajs)
 ho<-numeric(length(vv))
 for (i in 1:length(vv)){
   hi[i,]<-sample(1:1000,trajs)
   ho[i]<-max(rq[,hi[i,],i])
 }
 ho<-max(ho)
 
 {plotit<-function(){
   legendcex<-7
   lwds<-15
   col2<-"gray53"
   par(mfrow=c(2,3),oma=c(bottom=1, left=1, top=0, right=3),mar=c(bottom=23,left=29,top=15,right=6),cex.lab=10,cex.axis=8,mgp=c(21, 2, 4),xaxs='i',yaxs='i',las=1,cex.main=lwds)
   
 #par(mfrow=c(2,length(vv)/2),mar=c(bottom=5,left=6,top=3,right=.4),cex.lab=3,cex.axis=2,cex.main=4,mgp=c(3, .9, 0),xaxs='i')
 for (i in 1:length(vv)){
   matplot(rq[,hi[i,],i],lwd=lwds/3,type='l',lty=1,col=adjustcolor(viridis(30)[vv[i]],.3),main=paste(Faktor,"=",Wert1[vv[i]],sep=""),xlab="Time (Days)",ylab="Mean Waiting Time (Days)",ylim=c(0,ho),xaxt='n',yaxt='n')
   mtext(LETTERS[i],side=3,line=2,adj=0,cex=lwds)
   
   box(lwd=lwds/3*2)
   axis(side = 1, lwd = lwds/3,tck=-0.02,mgp=c(15,8,0))
   axis(side = 2, lwd =  lwds/3,tck=-0.02,mgp=c(12,5,0),las=2)
   }

 }
   wid<-5000
   hei<-2500
   png(filename = paste0(FigFol,"F14_IndTraj.png"),width = wid, height = hei,pointsize=10);plotit();graphics.off()
   pdf(file = paste0(FigFol,"F14_IndTraj.pdf"),width = wid, height = hei,pointsize=10);plotit();graphics.off()
 }
 #Fig14 Step Vari time---------------------------
 load("D:/Arbeit/Flatworms/ThePaper/FigureData/F25_compressed.RData")
 # par(mfrow=c(1,2),mar=c(bottom=2.4,left=2,top=.1,right=.8),cex.axis=.7,mgp=c(1, 0.1, 0),cex.lab=1)
 {plotit<-function(){
   legendcex<-7
   lwds<-15
   col2<-"gray53"
   par(mfrow=c(1,2),mar=c(bottom=23,left=29,top=15,right=6),cex.lab=10,cex.axis=8, oma=c(bottom=0, left=0, top=0, right=10),mgp=c(21, 2, 4),xaxs='i',yaxs='i',las=1,cex.main=lwds)
   x<-1:times
 cix=legendcex
 colori<-viridis(dim(waiti_vari2)[1])
 matplot(x,t(waiti_vari2[,x]),lwd=lwds,type='l',lty=1,xlab="Time",ylab="Variance in Waiting Time",col=colori,ylim=c(0,max(waiti_vari2[,x])),xaxt='n',yaxt='n')
 box(lwd=lwds/3*2)
 axis(side = 1, lwd = lwds/3,tck=-0.02,mgp=c(15,8,0))
 axis(side = 2, lwd =  lwds/3,tck=-0.02,mgp=c(12,5,0),las=2)
 legend(3000,810,as.character(w1),cex=cix,col=viridis(dim(waiti_vari2)[1]),pch=15,title=Faktor,ncol=5,bg="n",bty='n')
 mtext("A",side=3,line=2,adj=0,cex=lwds)
 
 matplot(x,t(waiti_CV2[,x]),lwd=lwds,type='l',lty=1,xlab="Time",ylab="CV Waiting Time",col=colori,ylim=c(0,1),xaxt='n',yaxt='n')
 box(lwd=lwds/3*2)
 axis(side = 1, lwd = lwds/3,tck=-0.02,mgp=c(15,8,0))
 axis(side = 2, lwd =  lwds/3,tck=-0.02,mgp=c(12,5,0),las=2)
 legend(7500,.9,as.character(w1),cex=cix,col=colori,pch=15,title=Faktor,ncol=5,bg="n",bty='n')
 mtext("B",side=3,line=2,adj=0,cex=lwds)
 
 }
   wid<-5000
   hei<-2000
   png(filename = paste0(FigFol,"F15_Vari.png"),width = wid, height = hei,pointsize=10);plotit();graphics.off()
   pdf(file = paste0(FigFol,"F15_Vari.pdf"),width = wid, height = hei,pointsize=10);plotit();graphics.off()
 }


#Fig7 Population size---------------
load("D:/Arbeit/Flatworms/ThePaper/FigureData/F25_compressed.RData")
nun<-t(popi[which(Wert1%in% c(20,50,70,85,110,300)),seq(1,dim(popi)[2],300)])
{plotit<-function(){
  legendcex<-7
  lwds<-15
  col2<-"gray53"
  par(mfrow=c(1,2),mar=c(bottom=23,left=29,top=15,right=6),cex.lab=10,cex.axis=8, oma=c(bottom=0, left=0, top=0, right=8),mgp=c(21, 2, 4),xaxs='i',yaxs='i',las=1,cex.main=lwds)
  matplot(nun,type='l',col=adjustcolor(rev(viridis(dim(nun)[2])),alpha.f=1),lty=1,xlab="Time",ylab="Average Population Size",lwd=lwds,xaxt='n',yaxt='n')
  box(lwd=lwds/3*2)
  axis(side = 1, lwd = lwds/3,tck=-0.02,mgp=c(15,8,0))
  axis(side = 2, lwd =  lwds/3,tck=-0.02,mgp=c(12,5,0),las=2)
  legend("topright",as.character(c(20,50,70,85,110,300)),cex=legendcex,col=(viridis(dim(nun)[2])),pch=15,title=Faktor,bg="white",ncol=3,bty='n')
  mtext("A",side=3,line=2,adj=0,cex=lwds)
  tutu<-rowMeans(popi)
plot(w1,tutu,xlab=Faktor,ylab="",pch=16,cex=lwds,xaxt='n',yaxt='n')
box(lwd=lwds/3*2)
axis(side = 1, lwd = lwds/3,tck=-0.02,mgp=c(15,8,0))
axis(side = 2, lwd =  lwds/3,tck=-0.02,mgp=c(12,5,0),las=2)
lines(w1,tutu,lty=1,col="forestgreen",lwd=lwds)
points(w1,tutu,pch=19,cex=lwds)
mtext("B",side=3,line=2,adj=0,cex=lwds)
}
wid<-5000
hei<-2000
png(filename = paste0(FigFol,"F16_PopSize.png"),width = wid, height = hei,pointsize=10);plotit();graphics.off()
pdf(file = paste0(FigFol,"F16_PopSize.pdf"),width = wid, height = hei,pointsize=10);plotit();graphics.off()
}


#Fig10 FIXI-------------

load("D:/Arbeit/Flatworms/ThePaper/FigureData/FIXI_compressed.RData")
{plotit<-function(){
  legendcex<-7
  lwds<-15
  col2<-"gray53"
  par(mfrow=c(1,1),mar=c(bottom=23,left=29,top=15,right=6),cex.lab=10,cex.axis=8, oma=c(bottom=0, left=0, top=0, right=8),mgp=c(21, 2, 4),xaxs='i',yaxs='i',las=1,cex.main=lwds)

print(matplot(w2,t(fin),type='l',lwd=lwds,col=viridis(length(w1)),xlab=Faktor2,ylab="Final waiting time",lty=1,xaxt='n',yaxt='n'))
box(lwd=lwds/3*2)
axis(side = 1, lwd = lwds/3,tck=-0.02,mgp=c(15,8,0))
axis(side = 2, lwd =  lwds/3,tck=-0.02,mgp=c(12,5,0),las=2)
legend("topleft",as.character(w1),cex=legendcex*3/2,ncol=3,col=viridis(length(w1)),pch=15,title=Faktor1,bg=adjustcolor("white",.7),bty='n')
}
wid<-5000
hei<-5000
png(filename = paste0(FigFol,"F17_FIXI.png"),width = wid, height = hei,pointsize=10);plotit();graphics.off()
pdf(file = paste0(FigFol,"F17_FIXI.pdf"),width = wid, height = hei,pointsize=10);plotit();graphics.off()
}


#Fig8 Diversity2-------------------
load("D:/Arbeit/Flatworms/ThePaper/FigureData/DIV2_compressed.RData")
#par(mfrow=c(1,1),mar=c(bottom=5,left=5,top=.45,right=.4),cex.lab=2,cex.axis=1.4,mgp=c(3.6, 2, 0))
{plotit<-function(){
  legendcex<-7
  lwds<-15
  col2<-"gray53"
  par(mfrow=c(1,1),mar=c(bottom=23,left=29,top=15,right=6),cex.lab=10,cex.axis=8, oma=c(bottom=0, left=0, top=0, right=8),mgp=c(21, 2, 4),xaxs='i',yaxs='i',las=1,cex.main=lwds)
  
print(matplot(waiter,col=viridis(dim(waiter)[2]),type='l',lty=1,lwd=lwds,ylab="Average Waiting Time",xlab="Time",xlim=c(0,times+3100),xaxt='n',yaxt='n'))
box(lwd=lwds/3*2)
axis(side = 1, lwd = lwds/3,tck=-0.02,mgp=c(15,8,0))
axis(side = 2, lwd =  lwds/3,tck=-0.02,mgp=c(12,5,0),las=2)
legend(rel_legend_pos(.6,.9)[1],rel_legend_pos(.7,.9)[2],as.character(w1),cex=legendcex,col=viridis(dim(waiter)[2]),bg=adjustcolor("white",.7),pch=15,title=Faktor,ncol=4)
range<-(max(waiter)*.95):(min(waiter)+(max(waiter)*.05))
points(rep(rel_legend_pos(0.915,.05)[[1]],length(range)),seq(rel_legend_pos(0.915,.05)[[2]],rel_legend_pos(0.915,.95)[[2]],length.out=length(range)),col=viridis(length(range)),pch=15,cex=lwds)
text(rel_legend_pos(0.96,.05)[1],max(range),"High Density",cex=legendcex-2)
text(rel_legend_pos(0.96,.05)[1],min(range),"Low  Density",cex=legendcex-2)

}
  wid<-5000
  hei<-2500
  png(filename = paste0(FigFol,"F18_Div.png"),width = wid, height = hei,pointsize=10);plotit();graphics.off()
  pdf(file = paste0(FigFol,"F18_Div.pdf"),width = wid, height = hei,pointsize=10);plotit();graphics.off()
}


# load("D:/Arbeit/Flatworms/ThePaper/FigureData/FIXIg_compressed.RData")
# print(matplot(w2,t(fin),type='l',lwd=3,col=viridis(length(w1)),xlab=Faktor2,ylab="Final waiting time",lty=1))
# legend("topleft",as.character(w1),cex=2,ncol=3,col=viridis(length(w1)),pch=15,title=Faktor1,bg=coli)
# load("D:/Arbeit/Flatworms/ThePaper/FigureData/FIXI_Detail_compressed.RData")
# print(matplot(w2,t(fin),type='l',lwd=3,col=viridis(length(w1)),xlab=Faktor2,ylab="Final waiting time",lty=1))
# legend("bottomright",as.character(w1),cex=2,ncol=3,col=viridis(length(w1)),pch=15,title=Faktor1,bg=coli)

#Fig11 Mathcases----------


#Fig15 Other fluctuations--------------------
load("D:/Arbeit/Flatworms/ThePaper/FigureData/Randi_compressed.RData")
x<-1:times
png(filename = paste0(FigFol,"F12_Other.png"),width = wid/5, height = hei/5,pointsize=10)
par(mar=c(bottom=0,left=6,top=.5,right=.3),oma=c(bottom=1, left=1, top=0, right=3),cex.axis=1.7,cex.lab=1.7,mgp=c(3, 1, 0))
layout(matrix(c(1,1,rep(3,9),2, 2,rep(4,9)), 11,2, byrow = F))
plot(x,KK[,2],lwd=5,type='l',lty=1,col="black",bty='n',xaxt='n',xlab="",ylab="Habitat Size",yaxt='n')
axis(2,at=c(10,70,300),labels=c(10,70,300),las=2)
plot(x,KK[,2],lwd=5,type='l',lty=1,col="black",bty='n',xaxt='n',xlab="",ylab="Habitat Size",yaxt='n')
axis(2,at=c(10,70,300),labels=c(10,70,300),las=2)
par(mar=c(bottom=5,left=6,top=.5,right=.3))
#par(mar=c(bottom=4,left=6,top=.5,right=.3),oma=c(bottom=1, left=1, top=0, right=3),cex.axis=1.7,cex.lab=1.7,mgp=c(3, 1, 0))
print(matplot(waiter,col=c("royalblue","firebrick4"),type='l',lty=1,lwd=3,ylab="average waiting time",xlab="time"))
legend("topleft",c("Random Habitat size","10-70-10-300-..."),cex=2,col=c("royalblue","firebrick4"),pch=15,ncol=1,bg="white",bty='n')
matplot(x,t(waiti_CV2[,x]),lwd=3,type='l',lty=1,xlab="time",ylab="CV waiting time",col=c("royalblue","firebrick4"))
legend("right",c("Random Habitat Size","10-70-10-300-..."),cex=2,col=c("royalblue","firebrick4"),pch=15,ncol=1,bg="white",bty='n')
graphics.off()

#Fig16 Robustness------------------
par(mfrow=c(1,2),mar=c(bottom=4.5,left=7,top=1.5,right=.4),cex.lab=3,cex.axis=2,mgp=c(3, .7, 0),xaxs='i',yaxs='i')
load("D:/Arbeit/Flatworms/ThePaper/FigureData/Robust_compressed.RData")
paras2<-c("Carrying capacity","Lifespan","Loci Inb. Depression","Loci Waiting Time","Allelic Diversity","Mutation Rate Inbreeding","Mutation Rate Waiting Time","Selection Coefficient","Sperm Storage Duration")
dimnames(WMf)<-list(Wert1,c("-5%","default","+5%"),paras2)

png(file=paste0("D:/Arbeit/Flatworms/ThePaper/Abb3/","Robust",".png"),width=2200,height=662)
print(levelplot(WMf,col.regions=rev(viridis(100)),par.strip.text=list(cex=2),ylab=list(label="Time In Low Density",cex=3),xlab=list(label="Habitat Size",cex=3),cex.lab=5,scales=list(x=list(cex=1.2,rot=45),y=list(cex=2))))
library(rasterVis)
library(grid)
grid::grid.text(label='Final Waiting Time', rot=90, y=unit(0.5, "npc"),x=unit(.96, "npc"),gp=gpar(fontsize=25))
graphics.off()
#Fig17 NoSotre---------------
# par(mar=c(bottom=5,left=6,top=1.5,right=.3),cex.axis=2,cex.lab=1.7,cex.main=2,mgp=c(3, 1, 0))
par(mfrow=c(1,2),mar=c(bottom=4.5,left=7,top=3,right=.4),cex.lab=3,cex.axis=2,mgp=c(3, .7, 0),xaxs='i',yaxs='i')
layout(matrix(c(0,1,1,0,2,2,3,3), 2, 4, byrow = TRUE))
load("D:/Arbeit/Flatworms/ThePaper/FigureData/NoStoreBoth_compressed.RData")
print(matplot(waiter,col=viridis(dim(waiter)[2]),type='l',lty=1,lwd=6,ylab="average waiting time",xlab="time",main="No Sperm or Egg Storage"))
legend(25000,38,as.character(w1),cex=1.5,col=viridis(dim(waiter)[2]),pch=15,title=Faktor,ncol=3,bg=adjustcolor("white",alpha.f=.4))
load("D:/Arbeit/Flatworms/ThePaper/FigureData/NoStoreEgg_compressed.RData")
print(matplot(waiter,col=viridis(dim(waiter)[2]),type='l',lty=1,lwd=6,ylab="average waiting time",xlab="time",main="No Egg Storage"))
legend(25000,28,as.character(w1),cex=1.5,col=viridis(dim(waiter)[2]),pch=15,title=Faktor,ncol=3,bg=adjustcolor("white",alpha.f=.4))
load("D:/Arbeit/Flatworms/ThePaper/FigureData/NoStoreSperm_compressed.RData")
print(matplot(waiter,col=viridis(dim(waiter)[2]),type='l',lty=1,lwd=6,ylab="average waiting time",xlab="time",main="No Sperm Storage"))
legend(25000,38,as.character(w1),cex=1.5,col=viridis(dim(waiter)[2]),pch=15,title=Faktor,ncol=3,bg=adjustcolor("white",alpha.f=.4))


#Fig20 Reciprocity---------------
load("D:/Arbeit/Flatworms/ThePaper/FigureData/Reciprocity_compressed.RData")
# par(mfrow=c(1,1),mar=c(bottom=7,left=8,top=3,right=.4),cex.axis=1.6,mgp=c(5, 2, 0))
rem<-apply(recip,2,mean)
res<-apply(recip,2,sd)
plot(Wert1,rem,cex.lab=3,cex.axis=2, xlab="Habitat Size", ylab= "Recoprocity",pch=21,type='b',lty=3,lwd=3,bg=viridis(length(Wert1)),ylim=c(0,max(c(rem+res,1))))
abline(h=c(1,0),lwd=3,col="gray53")
for(i in 1:length(res)){
  lines(rep(Wert1[i],2),rep(rem[i],2)+c(res[i],-res[i]),lwd=3,col="grey50")
  points(rep(Wert1[i],2),rep(rem[i],2)+c(res[i],-res[i]),cex=2,pch='-',col="grey50")
}
points(Wert1,rem,pch=21,cex=3,type='b',lty=3,lwd=3,bg=viridis(length(Wert1)))

#Fig21 Pertubation------------
load("D:/Arbeit/Flatworms/ThePaper/FigureData/Step_compressed.RData")
png(file="D:/Arbeit/Flatworms/ThePaper/Abbildungen2/Pertubation1.png",width=1905,height=1200)
par(mfrow=c(3,2),mar=c(bottom=4.5,left=8.5,top=1.5,right=.4),cex.lab=3,cex.axis=2,mgp=c(3, .7, 0),xaxs='i',yaxs='i')
for(i in 1:3){
  print(matplot(WMA[,,i],col=viridis(dim(WMA)[2]),type='l',lty=1,lwd=3,ylab="Average \n Waiting Time",xlab="Time",ylim=c(0,100)))
  abline(v=30000,lty=3,col="gray53",lwd=5)
  print(matplot(WMA[,,i],col=viridis(dim(WMA)[2]),type='l',lty=1,lwd=3,add=T))
  legend("topleft",as.character(Wert2),cex=2,col=viridis(dim(WMA)[2]),pch=15,title=Faktor1,bg=adjustcolor("white",alpha.f=.7),ncol=3,box.col=NA)
  print(matplot(CVA[,,i],col=viridis(dim(CVA)[2]),type='l',lty=1,lwd=3,ylab="CV \n Waiting Time",xlab="Time",ylim=c(0,1)))
  abline(v=30000,lty=3,col="gray53",lwd=5)
  print(matplot(CVA[,,i],col=viridis(dim(CVA)[2]),type='l',lty=1,lwd=3,add=T))
  legend("topright",as.character(Wert2),cex=2,col=viridis(dim(WMA)[2]),pch=15,title=Faktor1,bg=adjustcolor("white",alpha.f=.4),ncol=3,box.col=adjustcolor("white",alpha.f=.4))
  
}
graphics.off()
# Step Robustness------------------
load("D:/Arbeit/Flatworms/Ori/BT/WMS.RData")
WMS<-aWM*32
load("D:/Arbeit/Flatworms/Ori/BT/WMU.RData")
WMU<-aWM*38
WMa<-(WMS+WMU)/70
wma<-WMa
wma[,,,1]<-WMa[,,,3]
wma[,,,3]<-WMa[,,,1]
WMa<-wma
library(viridis)
library(lattice)
paras<-c("N","lifespan","genesINB","genesSELF","allelicDivers","inbMUT","selfMUT","selection","spermEXP")
paras2<-c("Number Of Individuals","Lifespan","Loci Inb. Depression","Loci Waiting Time","Allelic Diversity","Mutation Deleterious Alleles","Mutation Waiting Time","Selection Coefficient","Sperm Expiration Time")
Wert1<-c(100,500,seq(1000,10000,1000),20000)#low density time
Wert2<-c(100,500,seq(1000,10000,1000),20000)#high density time 
Wert3=1:9
replikate<-38
W1<-rep(Wert1,each=length(Wert2)*length(Wert3)*replikate)
W2<-rep(Wert2,length(Wert1)*length(Wert3)*replikate)
W3<-rep(rep(Wert3,each=length(Wert2)),length(Wert1)*replikate)
dimnames(WMa)<-list(Wert1,Wert2,paras,c("-5%","default","+5%"))
paras2<-c("Carrying capacity","Lifespan","Loci Inb. Depression","Loci Waiting Time","Allelic Diversity","Mutation Rate Inbreeding","Mutation Rate Waiting Time","Selection Coefficient","Sperm Storage Duration")
paras2<-c("Carrying capacity","Lifespan","Loci Inb. Depression","Loci Waiting Time","Allelic Diversity","Mutation Deleterious Alleles","Mutation Waiting Time","Selection Coefficient","Sperm Expiration Time")

dimnames(WMa)<-list(Wert1,c("-5%","default","+5%"),paras2)
for(iii in 1:length(Wert3)){
  png(filename=paste("D:/Arbeit/Flatworms/ThePaper/Abb3/",paras[iii],".png"),width=1400,height=662)
 print(levelplot(WMa[,,iii,],col.regions=rev(viridis(100)),main=list(paras2[iii],cex=3),par.strip.text=list(cex=2),ylab=list(label="Time In Low Density",cex=3),xlab=list(label="Time In High Density",cex=3),cex.lab=5,main.text.cex=5,scales=list(x=list(cex=2,rot=45),y=list(cex=2))))
 library(rasterVis)
 library(grid)
 grid::grid.text('Final Waiting Time', rot=90, y=unit(0.5, "npc"),x=unit(.985, "npc"),gp=gpar(fontsize=25))
  graphics.off()
}

#focals----------
load("D:/Arbeit/Flatworms/Ori/SR/compressed2.RData")
absch<-abs(change_wait)

relativeENDwt<-wait_end/avli
relativeCIR<-change_wait/avli
relativeABSCH<-abs(relativeCIR)
pars<-c("avli","gpm","gps","div","imut","smut","sco","eg","spex","ooo","uuu","verh")
pn<-c("Average Lifespan","LociInbreeding Depression","Loci Waiting Time","Allelic Diversity","Mutation Rate Inbreeding","Mutation Rate Waiting Time","Selection Coefficient","Time To Egg Production","Sperm Durability","Time Low Density","Time High Density","ProportionLow Density")

par(mfrow=c(1,1),mar=c(bottom=4.5,left=7,top=1.5,right=1),cex.lab=2,cex.axis=2,mgp=c(3, .7, 0),xaxs='i',yaxs='i')

focal3<-which(relativeENDwt>.10& relativeENDwt<.90&relativeABSCH<=sort(relativeABSCH[relativeENDwt>.10& relativeENDwt<.90])[10]) #10s smalles change in last circle
par(mfrow=c(2,6))
for(i in 1:length(pars)){
  boxplot(list(focal=get(pars[i])[focal3],all=get(pars[i])),ylab=pn[i],col=c(viridis(6)[c(2,5)]))
  title(main=LETTERS[i],adj=0,cex=5)
}


load("D:/Arbeit/Flatworms/ThePaper/FigureData/focal_compressed.RData")
par(mfrow=c(1,1),mar=c(bottom=4.5,left=7,top=1.5,right=.4),cex.lab=3,cex.axis=2,mgp=c(3, .7, 0),xaxs='i',yaxs='i')
print(matplot(waiter_normal,col=viridis(dim(waiter_normal)[2]),type='l',lty=1,lwd=3,ylab="average relative waiting time",xlab="time",main="Waiting time",ylim=c(0,.62)))
matplot(waiter_normal[,smut[focal3]>0.05],type='l',lty=2,col="black",add=T,lwd=3)
print(matplot(waiter_plus,col=viridis(dim(waiter_plus)[2]),type='l',lty=1,lwd=3,add=T))
print(matplot(waiter_plus[,smut[focal3]>0.05],type='l',lty=2,col="black",add=T,lwd=3))
print(matplot(waiter_minus,col=viridis(dim(waiter_minus)[2]),type='l',lty=1,lwd=3,add=T))
print(matplot(waiter_minus[,smut[focal3]>0.05],type='l',lty=2,col="black",add=T,lwd=3))

png(file="D:/Arbeit/Flatworms/ThePaper/Abbildungen2/PertubationFocal.png",width=1905,height=662)
par(mfrow=c(2,5),oma=c(bottom=1, left=1, top=0, right=3),mar=c(bottom=4.5,left=7,top=1.5,right=.4),cex.lab=3,cex.axis=2,mgp=c(3, .7, 0),xaxs='i',yaxs='i')
for(i in 1:10){
  plot(waiter_normal[,i],col=viridis(10)[i],type='l',lty=1,lwd=3,ylab=list("average relative waiting time",cex=2),xlab="time",ylim=c(0,.62))
  lines(waiter_plus[,i],col=viridis(10)[i],type='l',lty=1,lwd=3)
  lines(waiter_minus[,i],col=viridis(10)[i],type='l',lty=1,lwd=3)
  title(main=LETTERS[i],adj=0,cex=3)
  }
graphics.off()

load("D:/Arbeit/Flatworms/SR/FOCAL/Einzel.RData")
png(file="D:/Arbeit/Flatworms/ThePaper/Abbildungen2/Einzel.png",width=1905,height=662)
par(mfrow=c(2,5),oma=c(bottom=1, left=1, top=0, right=3),mar=c(bottom=4.5,left=7,top=1.5,right=.4),cex.lab=3,cex.axis=2,mgp=c(3, .7, 0),xaxs='i',yaxs='i')
for(i in 1:10){
  print(matplot(seq(0,50000,length.out=2000),t(Einzel[i,,]/avli[focal3]),xlab="Time",ylab=list("Relative Average Waiting Time",cex=2),ylim=c(min(Einzel/avli[focal3]),max(Einzel/avli[focal3])),type='l',lty=1,lwd=2,col=adjustcolor(viridis(10)[i],.6)) )
  abline(h=.5,col="gray53",lwd=2)
  abline(h=(spex[focal3]/avli[focal3])[i],col="gray53",lwd=2)
  title(main=LETTERS[i],adj=0,cex=5)
}
graphics.off()

#gall-----------------

load("FigureData/F25_compressed.RData")
load("FigureData/gall.RData")

galli<-gall[,-1]

# matplot(galli,type='l',lwd=3,xlab="Time",ylab="Freuency")
# mean(apply(galli[which(W==Wert1[i]),-10],1,sd))

for (i in 1:nrow(galli)){
  galli[i,]<-galli[i,]/sum(galli[i,])
}
library(ggplot2)
# test<-data.frame(data=as.vector(galli),Wert=as.factor(W),Allele=as.factor(rep(1:10,each=length(Wert1)*replikate)))
# ggplot(test, aes(x=Wert,y=data,fill=Allele)) + geom_violin()+ scale_fill_manual(values=adjustcolor(c("royalblue",rep("firebrick3",10)),alpha.f=.4))                                               
#    
test<-data.frame(Frequency=as.vector(galli),HabitatSize=as.factor(W),Allele=as.factor(c(rep(1,each=length(Wert1)*replikate),rep(2,each=length(Wert1)*replikate*9))))
# ggplot(test, aes(x=HabitatSize,y=Frequency,fill=Allele)) + geom_violin(trim=FALSE,scale="count",show.legend=T ,position="identity",width=3)+ scale_fill_manual(values=c("royalblue","firebrick3"))+theme_classic()                                           

galli2<-cbind(galli[,1],rowSums(galli[,-1])/9)
test<-data.frame(Frequency=c(rep(NA,length((min(Wert1):max(Wert1))[!((min(Wert1):max(Wert1))%in%Wert1)])),as.vector(galli2)),HabitatSize=as.factor(c((min(Wert1):max(Wert1))[!((min(Wert1):max(Wert1))%in%Wert1)],rep(W,2))),Allele=as.factor(c(rep(NA,length((min(Wert1):max(Wert1))[!((min(Wert1):max(Wert1))%in%Wert1)])),rep(c("wildtype","mutation"),each=length(Wert1)*replikate))))

galli3<-matrix(ncol=2,nrow=length(Wert1));for(i in 1:length(Wert1)){
 galli3[i,]<-colMeans(galli2[which(W==Wert1[i]),] )
}
test2<-data.frame(Frequency=as.vector(galli3),HabitatSize=rep(Wert1,2),Allele=as.factor(rep(c("wildtype","mutation"),each=length(Wert1))))

ggplot(test2, aes(x=HabitatSize,y=Frequency,fill=Allele)) + geom_violin(data=test, aes(x=HabitatSize,y=Frequency,fill=Allele),trim=FALSE,scale="width",show.legend=T ,position="identity",alpha=.6)+ scale_fill_manual(values=rev(c("royalblue","firebrick3")))+theme_classic()+geom_line(aes(x=as.numeric(HabitatSize)-10),lwd=1.3)+  geom_violin(data=test, aes(x=HabitatSize,y=Frequency,fill=Allele),trim=FALSE,scale="width",show.legend=T ,position="identity",alpha=.6) # +geom_point(pch=21,cex=3)       


# test<-data.frame(Frequency=as.vector(galli),HabitatSize=as.factor(W),Allele=as.factor(rep(1:10,each=length(Wert1)*replikate)))
# ggplot(test, aes(x=HabitatSize,y=Frequency,fill=Allele)) + geom_violin(trim=FALSE,scale="count",show.legend=T ,position="identity",width=1)+ scale_fill_manual(values=c("royalblue",adjustcolor(viridisLite::plasma(9),alpha.f=.4)))+theme_classic()                                           
rem<-c(300,200)
# ggplot(test, aes(x=HabitatSize,y=Frequency,fill=Allele)) + geom_violin(trim=FALSE,scale="count",show.legend=T ,position="identity",width=3)+ scale_fill_manual(values=c("royalblue","firebrick3"))+theme_classic()                                           

galli2<-cbind(galli[-which(W%in%rem),1],rowSums(galli[-which(W%in%rem),-1])/9)
V<-(min(Wert1[-which(Wert1%in%rem)]):max(Wert1[-which(Wert1%in%rem)]))[!((min(Wert1[-which(Wert1%in%rem)]):max(Wert1[-which(Wert1%in%rem)]))%in%Wert1[-which(Wert1%in%rem)])]
test<-data.frame(Frequency=c(rep(NA,length(V)),as.vector(galli2)),HabitatSize=as.factor(c(V,rep(W[-which(W%in%rem)],2))),Allele=as.factor(c(rep(NA,length(V)),rep(c("wildtype","deleterious"),each=length(Wert1[-which(Wert1%in%rem)])*replikate))))

galli3<-matrix(ncol=2,nrow=length(Wert1[-which(Wert1%in%rem)]));for(i in 1:length(Wert1[-which(Wert1%in%rem)])){
  galli3[i,]<-colMeans(galli2[which(W[-which(W%in%rem)]==Wert1[-which(Wert1%in%rem)][i]),] )
}
test2<-data.frame(Frequency=as.vector(galli3),HabitatSize=rep(Wert1[-which(Wert1%in%rem)],2),Allele=as.factor(rep(c("wildtype","deleterious"),each=length(Wert1[-which(Wert1%in%rem)]))))
#----------
png(file="D:/Arbeit/Flatworms/ThePaper/Abb3/alt_DelA.png",width=1500,height=1000)
XXXX<-paste(Wert1)
XXXX[XXXX%in%c(65,75,85)]<-""
ggplot(test2, aes(x=HabitatSize,y=Frequency,fill=Allele))  + 
  geom_violin(data=test, aes(x=HabitatSize,y=Frequency,fill=Allele),lwd=2,trim=FALSE,scale="width",show.legend=T ,position="identity",alpha=.6)+ 
  scale_fill_manual(values=rev(c("royalblue","firebrick3")))+
  theme_classic()+
  geom_line(aes(x=as.numeric(HabitatSize)-10),lwd=2)+  
  geom_violin(data=test, aes(x=HabitatSize,y=Frequency,fill=Allele),lwd=2,trim=FALSE,scale="width",show.legend=T ,position="identity",alpha=.6)  +  
  scale_x_discrete(breaks=as.character(Wert1),labels=XXXX)+ 
  theme(text=element_text(size=50),legend.key.size=unit(1.5, "cm"))+xlab("Habitat Size")
graphics.off()
```
