## Supplementary material for "Waiting for love but not forever: modelling the evolution of waiting time to selfing in hermaphrodites": Simulation Scripts: Clean_MW.R

```
Wert1<-c(seq(10,60,10),seq(65,85,5),seq(90,130,10),200,300) #Values for the habitat size g
# Wert1<-c(10,50,80,200)
replikate<-2

W1=rep(Wert1,replikate)
#print(paste( "Number of simulations: ",length(W1)))
seddi<-c(1:length(W1))#random number seeds

W3<-rbind(W1,seddi)
W3<-W3[,sort(W1,index.return=TRUE)$ix]
W1<-W3[1,]
seddi<-W3[2,]

idx1 <- as.numeric(Sys.getenv('SLURM_PROCID'))+1 # get the processID

iterationnum <- as.numeric(commandArgs(trailingOnly=T))[1] #retrieve the parameter you want to use in this simulation.
cores_to_use <- as.numeric(commandArgs(trailingOnly=T))[2] #retrieve the parameter you want to use in this simulation.

idx2<-((1:cores_to_use)+(cores_to_use*(iterationnum-1)))[idx1]
if(!is.na(idx2)){ # use this to make sure, thare are parameters left for this simulation.
  W1<-W1[idx2]
  seddi<-seddi[idx2]


  set.seed(seddi)
  # cat(noquote(paste("\n Replikatnummer: ",seddi,"  Simulationen: ",length(Wert1)*replikate,"\n ",seddi/length(seddi)*100,"%",sep="")))
  # Define Parameters-------------------------------------------------------------------------------------------------------
  #_________________________________________Define Parameters___________________________________________________
  
  N<-200               # number of individuals at t0
  K<-N #carrying capacity
  grid<-W1#sqrt(N/D), g in the paper
  times<-30000          # number of steps observed
  
  av_lifespann<-100    # average number of steps an individual will live before dying of natural causes. In optimal conditions. 1/d
  
  genesper_maturity<-20#8 # how many genes should determine reaching maturity, inbreeding loci I
  genesper_selfprop<-5 # how many genes should determine selfing propencity, waiting time loci W
  diversity<-10 # number of different alleles per locus
  mutation<-3#/(genesper_maturity*2)#7*10^-6*100#.005 #probability of alleles mutating in %
  smutation<-.05#.05 #standard deviation for normal distribution that pulls mutations on selfing alleles
  domcoef<-0 #dominance coeficient if NA, coeficient is random
  selcoef<-.2 # 1 minus the selection coeficient; if NA, coefficient is random
  FIXI<-NA #can be used to fix the mortality of selfed offspring; if NA it is not fixed
  divers<- FALSE # should the population start with uniform mix of alleles at the inbreeding loci, or just All Allele 1
  refuge<-TRUE#should there be a "default gene" that is not selected against

  egg_grow<-1 #number of rounds until 1 egg is produced
  sperm_expire<-10 #how long should sperm last? put NA for eternal sperm durability
    #___________________________________________________________________________________________________________________
  
  Modus<-"Bottleneck" #bottleneck for bottleneck simulation, Function for K_func simulation
  #Bottleneck with no strength is used for constant densities. This script is the "basic" one doing different constant habitat sizes. 
  
  # Bottleneck
  timetobot<-1 #when should the first bottleneck occur
  bottleneck_freq<-50 #every how many round should a bottlenack occur
  bottleneck_strength<-0 #how many percent of the population should die in bottleneck
  
  # Function
  
  Kfuni<- "Bottleneck" #which function to use in generatin K, options: "Sinus", "Step", "Slope"
 #This is Bottleneck, although that's not an option in order to avoid confusion. It doesn't harm and avoids for sure that the later conditions become true. 
  
   gr_max<-300#grid^2*0.02           # Max N in habitat (Capacity) 
  gr_min<-50     # Min Capacity "bottleneck
  # "Sinus"
  freq_sin<-8000/2
  # "Step"
  oben<-3000 #how many rounds should the Capacity be Kmax
  unten<-3000 #how many rounds should the Capacity be Kmin
  startpos<-"unten" #at what condition should K start
  # "Slope"
  slopetype<-"negative" #should there be a negative or positive slope between Kmin & Kmax
  #_____________________________________________________________________________________________________________
  # Variable Prep ----------------------------------------------------------------------------------------------
  
  reserve<-K*1.5 #reserve is very important to make the code run smoothly. It helps creating "empty" space in vectors, so that vectors don't have to change size. 

  gr<-grid
  nu=N # Starting value in counting Vektor nu
  waitmean<-c() #prep vector for mean waittimes per round
  wantthat<-waitmean
  wata<-matrix(-1,nrow=times,ncol=3)
  ID<-seq(1:N) # set starting ID for each individual
  id=max(ID) #max ID number makes new ID for offspring easier
  d<-100/av_lifespann #deathrate in %
  #Tracking birth and death caues
  Bself<-rep(0,times)
  Bself_sperm<-rep(0,times)
  Bmate<-rep(0,times)
  Bmate_sperm<-rep(0,times)
  
  Dinb_mate<-rep(0,times)
  Dinb_self<-rep(0,times)
  Dnat<-rep(0,times)

  
  KK<-c() #vector for Capacity over time
  bottleneck_time<-seq(timetobot,times-1,bottleneck_freq) #time points for bottlenecks
  kkillcount<-c() #vector to contain how many individuals are killed in each bottleneck


  delfreq<-0
  

  reciproc<-0
  mcounter<-0
  
  waitvari<-numeric(times)-1
  waitmean<-rep(-1,times) #tracking mean waiting time per round
  wira<-matrix(NA,nrow=times/av_lifespann,ncol=reserve)
  ori<-matrix(NA,nrow=times/av_lifespann,ncol=reserve)
  
  # eggs available to starting population 

    offnum<-c(rpois(length(ID),av_lifespann/10),rep(-1,K))
  blocki<-rep(-1,reserve) #storing the information who has mated yet
  blocki[1:N]<-0
  eggi<-1/egg_grow# eggs added per round
  sincemate<-c(rep(0,length(ID)),rep(-1,K))# vector of rounds since mating for each individual, important to find the time for selfing
  
  #generate values for Capacity
  if (Modus=="Function"){ 
    if (Kfuni=="Step"){
      Stepi<-c()
      while (length(Stepi)<times){
        if (startpos=="unten"){ 
          Stepi<-c(Stepi,rep(gr_min,unten),rep(gr_max,oben))
        }else if(startpos=="oben"){ 
          Stepi<-c(Stepi,rep(gr_max,oben),rep(gr_min,unten))
        }}
      Stepi<-Stepi[1:times]}
    
    if (Kfuni=="Slope"){
      if (slopetype=="positive"){
        slope<-(gr_max-gr_min)/(times)
      }else if (slopetype=="negative"){
        slope<-(gr_min-gr_max)/(times)
      }}}
  
  # make infertile list-------------------------------------------------------------------------
 
  # Function Prep----------------------------------------------------------
  
  # genetrack <- function (genetics,type,numdiff,ID){
  #   # gives the number of times each dominant/recessive/intermediate gene is in the population
  #   
  #   # genetlist - list of genotypes as generated in firstgenes/gene_init/genemate
  #   # type - information on type of expression, "dri" for dominant/recessive/intermediate genes, "adi" for aditive genes
  #   # numdiff - information on how many differend types of Aleeles are on each locus, information is not acessed for "adi" just give NA
  #   # ID - positions in genetics for all living individuals
  #   

  
  genemate<- function (IDmother,IDfather,genetics,h,mut,muti){
    # generates a new genotype for offspring of a mating between genotype IDmother and genotype IDfather
    
    # IDmother/ID father - position in genetics of genetic information of mating individual
    # genetics - list with genotypes of the population
    # type - information on type of expression, "dri" for dominant/recessive/intermediate genes, "adi" for aditive genes
    # h - list of matrices of dominance coeficients for alleles. give NA if adi
    
    
    genesmother<-as.vector(genetics[,c(1,2),IDmother])
    genesfather<-as.vector(genetics[,c(1,2),IDfather])
    gennum<- c(dim(genetics[,c(1,2),IDmother])[1])
    baby<-matrix(0,nrow=gennum,ncol=3)
    
    ## stop("in genmate")
    frompa<-sample(c(1,2),gennum,replace = TRUE)
    fromma<-sample(c(1,2),gennum,replace = TRUE)
    fromma2<-fromma
    frompa2<-frompa
    
    fromma[fromma2==1]<-(1:gennum)[fromma2==1]
    fromma[fromma2==2]<-(1:gennum)[fromma2==2]+gennum
    frompa[frompa2==1]<-(1:gennum)[frompa2==1]
    frompa[frompa2==2]<-(1:gennum)[frompa2==2]+gennum
    
    baby[,c(1,2)]<-cbind(genesmother[fromma],genesfather[frompa])
    #print(baby)
    

      inds1 <- which(runif(genesper_maturity*2)<mut/100)
      t1<-genesper_maturity


      inds2<-c()
      t2<-0
    inds<-c(inds1,inds2)
    tt<-(1:(t1+t2))+genesper_selfprop
    if(length(inds)  > 0){
      
      baby[tt,1:2][inds] <- ((baby[tt,1:2][inds] - 1) + ceiling(runif(length(inds),0,diversity-1))) %% diversity + 1
      
    }
        
  
    #Method1
    baby[1:genesper_selfprop,c(1,2)]<-exp(log(baby[1:genesper_selfprop,c(1,2)])+log(rgamma(genesper_selfprop*2,shape =1^2/muti^2 , scale =muti^2/1 )))
    
   
    if (is.matrix(baby[1:genesper_selfprop,c(1,2)])){
      baby[1:genesper_selfprop,3]<-rowMeans(baby[1:genesper_selfprop,c(1,2)])
    }else{
      baby[1,3]<-mean(baby[1,c(1,2)])
    }
    
    
    baby[(1:genesper_maturity)+genesper_selfprop,3]<-h[cbind(baby[(1:genesper_maturity)+genesper_selfprop,1],baby[(1:genesper_maturity)+genesper_selfprop,2],1:genesper_maturity)]
    
    
    
    return(baby)
  }
  

  firstgenes<-function (genetics,ID,numdiff,type,h,av_lifespann,divers){
    # matix - list of gene matrix, having 3 collums col1=allele1, col2=allele2, col3=phenotype
    # nimdiff - how many differend alleles there are as starting condition for each locus
    # type -"dri" for dominant/recessiv or intermediate alleles, "adi" for aditive Alleles
    # h - list of matrices of dominance coeficients for alleles. give NA if adi
    n<-1:length(ID)
    for (i in 1:dim(genetics)[1]){
      for(n in 1:length(ID)){
        if (type[i]=="dri"){
          
          if (!divers){
            genetics[i,1,n]<-1
            genetics[i,2,n]<-1
          } else{
            genetics[i,1,n]<-sample(seq(1,numdiff[i],1),1,replace=TRUE)
            genetics[i,2,n]<-sample(seq(1,numdiff[i],1),1,replace=TRUE)
          }
          
          genetics[i,3,n]<-h[genetics[i,1,n],genetics[i,2,n],i-genesper_selfprop]
        } else if (type[i]=="adi"){
  
          genetics[i,1,n]<-seq(0.001,1,length.out = length(ID))[n]
          genetics[i,2,n]<-seq(0.001,1,length.out = length(ID))[n]
          genetics[i,3,n]<-mean(c(genetics[i,1,n],genetics[i,2,n]))
        }}}
    
    return(genetics)
  }
  offspringrisk <- function(ID,genetics,selcoef){
    #Input:
    #ID: vector with the individual IDs, providing the number of living individuals.
    #genetics: a 3-dimensional array with dimension 1 containing 25 different loci, dimension 2 representing chromosome 1, 2 and the resulting effect, and dimension 3 for each individual.
    #In dimension 1, the last 20 loci are needed, as they store the potentially deleterious alleles.
    #In dimension 2, column 1 and 2 are compared, and for each individual, Nhomd (the number of homozygous loci with deleterious alleles), Nhetd (the number of hterozygous loci with deleterious alleles), and Nhetr (the number of heterozygous loci with one refuge allele) are determined
    #From Nhomd,Nhetd and Nhetr, the offspring risk is calculated
    #selcoef: Selection coefficient influencing the offspring risk
    #Output:
    #A vector with numerical values, containing the offspring risk for each individual
    if(refuge){ #if refuge==T, allele nr. 1 is not selected against.
      Nxixi <- colSums(genetics[-1:-genesper_selfprop,1,1:length(ID)] == genetics[-1:-genesper_selfprop,2,1:length(ID)]&(genetics[-1:-genesper_selfprop,1,1:length(ID)]!=1&genetics[-1:-genesper_selfprop,2,1:length(ID)]!=1)) #The locus is deleteriously homozygous if both alleles are deleterious and col. 1 and 2 are equal.
      Nxixj <- colSums(genetics[-1:-genesper_selfprop,1,1:length(ID)] != genetics[-1:-genesper_selfprop,2,1:length(ID)]&(genetics[-1:-genesper_selfprop,1,1:length(ID)]!=1&genetics[-1:-genesper_selfprop,2,1:length(ID)]!=1)) #The locus is heterozygous with deleterious alleles if the columns differ and no column is the refuge allele.
      Nwxi <- colSums(genetics[-1:-genesper_selfprop,1,1:length(ID)] != genetics[-1:-genesper_selfprop,2,1:length(ID)]&(genetics[-1:-genesper_selfprop,1,1:length(ID)]==1|genetics[-1:-genesper_selfprop,2,1:length(ID)]==1)) #The locus is heterozygous with one refuge allele if the columns differ and one column is the refuge allele.
      # offspringrisk <-(1-selcoef)^Nhomd*(((1-selcoef)*0.5))^Nhetd*((1-selcoef)*0.25)^Nhetr #calculating expected offspring survivability
      # offspringrisk <-1- (selcoef)^Nhomd*((selcoef*0.5))^Nhetd*(selcoef*0.25)^Nhetr #calculating expected offspring survivability
      s<-1-selcoef
      offspringrisk <-1-(1-s)^Nxixi*(1-s*.5)^Nxixj*(1-s*.25)^Nwxi
    }else{
      Nhomd <- colSums(genetics[-1:-genesper_selfprop,1,1:length(ID)] == genetics[-1:-genesper_selfprop,2,1:length(ID)]&(genetics[-1:-genesper_selfprop,1,1:length(ID)]!=1&genetics[-1:-genesper_selfprop,2,1:length(ID)]!=1)) #The locus is deleteriously homozygous if both alleles are deleterious and col. 1 and 2 are equal.
      Nhetd <- colSums(genetics[-1:-genesper_selfprop,1,1:length(ID)] != genetics[-1:-genesper_selfprop,2,1:length(ID)]&(genetics[-1:-genesper_selfprop,1,1:length(ID)]!=1&genetics[-1:-genesper_selfprop,2,1:length(ID)]!=1)) #The locus is heterozygous with deleterious alleles if the columns differ and no column is the refuge allele.
      offspringrisk <- (selcoef)^Nhomd*((selcoef*0.5))^Nhetd #calculating expected offspring survivability
    }
    return(offspringrisk) #vector for offspring risk of all individuals
  }
  
  gnum<-genesper_maturity+genesper_selfprop
  kill<-function(z){
    if(length(z)>0){
    replacer<-rep(-1,length(z))
    ID<<-ID[-z]
    offnum<<-c(offnum[-z],replacer)
    sincemate<<-c(sincemate[-z],replacer)
    
    wait_time<<-c(wait_time[-z],replacer)
    blocki<<-c(blocki[-z],replacer)

    genetics<<-array(c(genetics[,,-z],numeric(length(z)*3*(gnum))-1),dim=c(gnum,3,reserve))
    sperm<<-array(c(sperm[,,-z],numeric(length(z)*3*(gnum))-1),dim=c(gnum,3,reserve))

}
  }
  
  # Genes Prep------------------------------------------------
  # # 
  # creating a list of genmatrices, where each row represents 1 locus
  # column1= Allel 1
  # column2=Allel2
  # collum3=Genotype (mean of additive genes, or 
  # result of dominance-selection matrix for gene combination)
  # Matrices are added to the list at each birth, but the list is never shortened.
    genetics<-array(-1,dim=c((genesper_maturity+genesper_selfprop),3,(reserve)))
    ggg<-genesper_maturity+1
  sperm<-genetics
  #creating dominance-selection Matrices by random.
  numdiff<-c(rep(NA,genesper_selfprop),rep(diversity,ggg-1)) #5 is the number of differend Alleles that can be dominant or recessive
  fibonacci<-0
  for (i in 1:diversity-1){
    fibonacci<- fibonacci+i}#number of entries needet to fill triangular matrix
  domi<-list()
  sele<-list()
  type<-c(rep("adi",genesper_selfprop),rep("dri",genesper_maturity))
  h<-array(-1,dim=c(diversity,diversity,genesper_maturity))
  if(is.na(domcoef) | is.na(selcoef)){
    for (i in (genesper_selfprop+1):length(type)){
      hd<-diag(1,nrow=diversity) #dominance coefficient
      if (is.na(domcoef)){
        hd[lower.tri(hd)]<-runif(fibonacci,0,1)
      }else{
        hd[lower.tri(hd)]<-rep(domcoef,fibonacci)
      }
      rs<-rep(1:diversity,(1:diversity)-1)
      cs<-c()
      for (cc in 2:diversity){
        cs<-c(cs,1:(cc-1))
      }
      for (j in 1:fibonacci){
        hd[upper.tri(hd)][j]<-hd[rs[j],cs[j]]
      }
      if(refuge){
        hd[1]<-1#Genotype 1/1 is always "basic genotype" with dominance 1 and selection 0
      }
      domi[[i]]<-hd
      if (is.na(selcoef)){
        sss<-diag(c(runif((diversity),0.5,1)),nrow=diversity) # selection coefficient
        sss[lower.tri(sss)]<-runif(fibonacci,0,1)
        rs<-rep(1:diversity,(1:diversity)-1)
        cs<-c()
        for (cc in 2:diversity){
          cs<-c(cs,1:(cc-1))
        }
        for (j in 1:fibonacci){
          sss[upper.tri(sss)][j]<-sss[rs[j],cs[j]]
        }

      }else{
        sss<-diag(selcoef,nrow=diversity) # selection coefficient
        sss[lower.tri(sss)]<-rep(selcoef,fibonacci)
        rs<-rep(1:diversity,(1:diversity)-1)
        cs<-c()
        for (cc in 2:diversity){
          cs<-c(cs,1:(cc-1))
        }
        for (j in 1:fibonacci){
          sss[upper.tri(sss)][j]<-sss[rs[j],cs[j]]
        }
      }
      
      #quasinormal genes are less selected against
      #_____________________
      if(refuge){
        sss[1]<-0#Genotype 1/1 is always "basic genotype" with dominance 1 and selection 0
      }
      sele[[i]]<-sss
      h[,,i]<-1-(hd*sss)
    }
  }else {
    xh<-matrix(1-selcoef*domcoef,diversity,diversity)
    diag(xh)<-selcoef
    if(refuge){
      xh[1]<-1
    }
    h<-array(xh,dim=c(diversity,diversity,genesper_maturity))
  }
  genetics<-firstgenes(genetics,ID,numdiff,type,h,av_lifespann,divers)#generation of starting genotypes
  # Waiting time--------------------
  wait_time<-apply(genetics[1:genesper_selfprop,1:2,],3,mean)*av_lifespann
  #>----------------------------------
  # Main Loop ---------------------------------------------------------------
  
  gr=grid
  #____________________________________________Start of main loop__________________________________________________________
  
  for (i in 1:times){ #rahmenfunktion. repeat steps t times
    if (length(ID)>0){ #make it stop at extinction
      if (Modus=="Bottleneck"&bottleneck_strength>0&i %in% bottleneck_time){ #bottlenecking population if desired
            killcount<-round(length(ID)*(bottleneck_strength/100))# calculates hor many individuals will be killed
            if (killcount>0){
              necks1<-sample(ID,killcount)#samples random IDs to kil
              necks<-which(ID %in% necks1)#gets indicee of killed individual
              kkillcount<-c(kkillcount,killcount)
              kill(necks)
            }
      }else if (Modus=="Function"){ #sets up current capacity if K is a function of time
        if (Kfuni=="Sinus"){
          amp<-(gr_max-gr_min)/2
          b<-(2*pi)/freq_sin
          gr<-amp*sin(b*i)+mean(c(gr_min,gr_max))
        }else if (Kfuni=="Step"){
          gr=Stepi[i]
        }else if (Kfuni=="Slope"){
          if (slopetype=="positive"){
            gr<-slope*i+gr_min
          }else if (slopetype=="negative"){
            gr<-slope*i+gr_max
          }}
        KK[i]<-gr
      }
      grid<-gr
      
      # Die ---------------------------------------------------------------------
        reaper<-which(runif(length(ID),0,100)<=d)
kill(reaper)
    }
    if(length(ID)>0){
        offnum[1:length(ID)]<-offnum[1:length(ID)]+eggi
      # sincemate---------------------------time since last mating, to check if waiting time was reached
        sincemate[sincemate>=0]<-sincemate[sincemate>=0]+1 
      
      if (!is.na(sperm_expire)){
        sperm[,,which(sincemate>sperm_expire)]<- -1
        blocki[which(sincemate[1:length(ID)]>sperm_expire & blocki[1:length(ID)]==ID)]<-0 #blocki records the previous partner. This way even during sperm usage it is possible to find out if offspring is produced from selfing or outcrossing sperm. This allows to track: degree of reciprocity and outcrossing/selfing mortality rates, and moreover enables us to run the experiments with fixed selfing mortality. 
      }
      
      # Mating detection----------------
      eprob<-(1/(grid^2))
      numpairs <- choose(length(ID),2)
      pairs <- cbind(rep(ID,rep(length(ID),length(ID))),rep(ID,length(ID)))
      pairs <- subset(pairs,pairs[,1]<pairs[,2])
      numencounters <- rbinom(1,numpairs,eprob)
      epairids <- sample(numpairs,numencounters)#gives the ids in random order, so no need for shuffling
      
      spermsi<-ID[which(offnum[1:length(ID)]>=1 & sperm[1,1,1:length(ID)]>=0)]
      selfer<-ID[which(wait_time[1:length(ID)]<sincemate[1:length(ID)] & sperm[1,1,1:length(ID)]==-1)]
      mater <- c(pairs[epairids,1],selfer,spermsi)
      partner <- c(pairs[epairids,2],selfer,rep(0,length(spermsi)))
      
      #>-------------------  
  
      # Mating Event ------------------------------------------------------------
      if (length(mater)>=1) { #when matings occur == collisions detected
              psuc<- max(min((K-length(ID))/sum(offnum[which(ID %in% unique(c(partner,mater))[!unique(c(partner,mater))==0])]),1),0) #sucess depends on capacity
        for (f in 1:length(mater)){ #loops though matings == positions  in indiceees by 2
            p<-which(ID==partner[f])#indcee for partner comonly used
            m<-which(ID==mater[f])#indcee for mater comonly used
              #generating litter size, saving sperm, saving information on previous encounters and settingtime since mating back
              if(mater[f]!=partner[f] & partner[f]!=0){ #mating
                if(max(c(offnum[m],offnum[p]))>0){
                  reciproc<-reciproc+(min(c(offnum[m],offnum[p]))/max(c(offnum[m],offnum[p])))
                }
                mcounter<-mcounter+1
                wurfgrose<-rbinom(1,floor(offnum[m])+floor(offnum[p]),psuc)
                blocki[m]<-partner[f]
                blocki[p]<-mater[f]
                sperm[,,m]<-genetics[,,p]
                sperm[,,p]<-genetics[,,m]
                sincemate[m]<-0 
                sincemate[p]<-0
                offnum[m]<-offnum[m]-floor(offnum[m])  #set back egg number
                offnum[p]<-offnum[p]-floor(offnum[p])
              }else if (mater[f]==partner[f]){#selfing
                wurfgrose<-rbinom(1,floor(offnum[m]),psuc) 
                blocki[m]<-mater[f]
                sperm[,,m]<-genetics[,,m]
                sincemate[m]<-0
                offnum[m]<-offnum[m]-floor(offnum[m])  #set back egg number
              } else if(partner[f]==0){#sperm
                if(blocki[m]!=mater[f]){#if sperm is from other individual, not from selfing
                  if(length(which(ID==blocki[m]))==1){# if the ex partner the sperm is from is still alive
                    if(blocki[which(ID==blocki[m])]==mater[f]){ #and he has the maters sperm still
                      reciproc<-reciproc+1
                      mcounter<-mcounter+1
                    }
                  }else{
                    reciproc<-reciproc+0
                    mcounter<-mcounter+1 
                  }
                }
                wurfgrose<-rbinom(1,floor(offnum[m]),psuc) 
                offnum[m]<-offnum[m]-floor(offnum[m])  #set back egg number
              }
              if (wurfgrose>0){
                for (w in 1:wurfgrose){ #generates litter size as in wurfgr
                  if(mater[f]!=partner[f] & partner[f]!=0){ # case: mating
                    Bmate[i]<-Bmate[i]+1
                    selfing<-FALSE
                  }else if (mater[f]==partner[f] ){ #case: selfing
                    Bself[i]<-Bself[i]+1
                    selfing<-TRUE
                  } else if (partner[f]==0 & blocki[m]==mater[f]){ #case: selfing sperm
                    Bself_sperm[i]<-Bself_sperm[i]+1
                    selfing<-TRUE
                  }else if (partner[f]==0 & blocki[m]!=mater[f]){ #case: mating sperm
                    Bmate_sperm[i]<-Bmate_sperm[i]+1
                    selfing<-FALSE
                  }
                  # Inbreeding Depression-------------------------------------------
                  if (partner[f]!=0){
                    babyworm<-genemate(m,p,genetics,h,mutation,smutation) #generates genotype (mating/selfing)
                  }else if (partner[f]==0){
                    thisone<-array(-1,dim=c(dim(genetics[,,m])[1],dim(genetics[,,m])[2],2))
                    thisone[,,1]<-genetics[,,m]
                    thisone[,,2]<-sperm[,,m]
                    babyworm<-genemate(1,2,thisone,h,mutation,smutation) #generates genotype (sperm)
                  } 
                  if(!is.na(FIXI)){
                    if(selfing){
                      phenin<-1-FIXI
                    }else if(!selfing){
                      phenin<-1
                    }
                  }else{
                    phenin<- prod(babyworm[((genesper_selfprop+1):gnum),3]) #survival prob
                    }
                  if (runif(1,0,1)<=(1-phenin)){ #if baby dies from inbreeding, record that death
                    if(blocki[m]!=mater[f]){
                      Dinb_mate[i]<-Dinb_mate[i]+1 
                      }else {
                        Dinb_self[i]<-Dinb_self[i]+1
                    }
                  }else{ #if baby survives, generate new individual and its properties, also track data
                    ID<-c(ID,id+1) # erstenllen einer neuen ID fr den neuen Punkt
                    id<-id+1
                    genetics[,,length(ID)] <-babyworm
                    sperm[,,length(ID)]<-c(-1)
                    blocki[length(ID)]<-0
                    wait_time[length(ID)]<-mean(babyworm[1:genesper_selfprop,c(1,2)])*av_lifespann
                    # phen<-phentrack(phen,babyworm,genesper_selfprop,hkats,"live")
                    # track<-genetrack(genetics,length(ID),"birth",track)
                    offnum[length(ID)]<-0
                    sincemate[length(ID)]<-0
                  }}
                }}}}
      

      #>------------------------- 
      # Grafik & Tracking ------------------------------------------------------------------
      waitvari[i]<-var(wait_time[1:length(ID)])
      #________________________________________________Grafik & Verarbeitung____________________________________________________________

      delfreq<-delfreq+sum(genetics[(1:genesper_maturity)+genesper_selfprop,c(1,2),1:length(ID)]!=1)/(length(ID)*genesper_maturity*2)
        #Grafic Output
        nu=c(nu,length(ID))
      waitmean[i]<-mean(wait_time[1:length(ID)])
      wantthat[i]<-sum(genetics[(genesper_selfprop+1):dim(genetics)[1],c(1,2),1:length(ID)]!=1)/length(ID)
      Nhomd <- colSums(genetics[-1:-genesper_selfprop,1,1:length(ID)] == genetics[-1:-genesper_selfprop,2,1:length(ID)]&(genetics[-1:-genesper_selfprop,1,1:length(ID)]!=1&genetics[-1:-genesper_selfprop,2,1:length(ID)]!=1)) #The locus is deleteriously homozygous if both alleles are deleterious and col. 1 and 2 are equal.
      Nhetd <- colSums(genetics[-1:-genesper_selfprop,1,1:length(ID)] != genetics[-1:-genesper_selfprop,2,1:length(ID)]&(genetics[-1:-genesper_selfprop,1,1:length(ID)]!=1&genetics[-1:-genesper_selfprop,2,1:length(ID)]!=1)) #The locus is heterozygous with deleterious alleles if the columns differ and no column is the refuge allele.
      Nhetr <- colSums(genetics[-1:-genesper_selfprop,1,1:length(ID)] != genetics[-1:-genesper_selfprop,2,1:length(ID)]&(genetics[-1:-genesper_selfprop,1,1:length(ID)]==1|genetics[-1:-genesper_selfprop,2,1:length(ID)]==1)) #The locus is heterozygous with one refuge allele if the columns differ and one column is the refuge allele.
      wata[i,]<-c(mean(Nhomd),mean(Nhetd),mean(Nhetr))
    
      
      if (i%%av_lifespann==0){
        wira[i/av_lifespann,]<-colMeans(genetics[1:genesper_selfprop,3,])
        ori[i/av_lifespann,1:length(ID)]<-offspringrisk(ID,genetics,selcoef)
      }
    }#ende von     for (i in 1:times){  &  if (length(ID)>0){ 
  #>---------------------
  # End Main loop---------------------------------------------

  defr<-delfreq/times
  fitself<-(Dinb_self/(Bself+Bself_sperm))
  fitmate<-(Dinb_mate/(Bmate+Bmate_sperm))
  sesult<-sum(Bself+Bself_sperm)/sum(Bself+Bself_sperm+Bmate+Bmate_sperm)
  
  Wert1<-Wert1
  replikate<-replikate
  reciprociality<-reciproc/mcounter
#   
  save(list=ls(),file=paste("clean_",sprintf("%05d", seddi),".RData",sep=""))
}
```
