## Supplementary material for "Waiting for love but not forever: modelling the evolution of waiting time to selfing in hermaphrodites": Simulation Scripts: CSolv.R

```
rm(list=ls())

daten<-dir(pattern="*RData")
Faktor<-"Habitat size"

times<-30000 
extinction<-c()
W<-c()


Wert1<-c(seq(10,60,10),seq(65,85,5),seq(90,130,10),200)
replikate<-1000


# W=rep(Wert1,replikate)

# W1<-c()
# ind<-c()
# for (xi in 1:length(daten)){
#   load(daten[xi])
#   if (!is.na(W1) & W1==grid){
#     ind<-c(ind,xi)
#     rm(list=ls())

extinction<-c()
W<-numeric(length(daten))


#   }}

fits<-c()
load(daten[1])
popi<-matrix(0,nrow=length(Wert1), ncol=times+1)
waiti<-matrix(0,nrow=length(Wert1), ncol=times)
waiti_vari<-numeric(length(Wert1))
waiti_vari2<-waiti
# self1<-matrix(NA,length(daten), ncol=times)
# mate1<-matrix(NA,length(daten), ncol=times)
# fm<-matrix(NA,length(daten), ncol=times)
# fs<-matrix(NA,length(daten), ncol=times)
# fit1<-matrix(NA,length(daten), ncol=101)
# fit2<-matrix(NA,length(daten), ncol=101)
# fitfin<-matrix(NA,length(daten), ncol=101)
pp<-numeric(length(Wert1))
delis<-matrix(0,length(Wert1), ncol=length(defr))
# waitgen<-array(NA,dim=c(11,times,length(daten)))
wiria<-array(dim=c(ncol(wira),length(Wert1),replikate))
oria<-wiria
jup<-numeric(length(Wert1))
# erg<-array(-1,dim=c(times,3,length(daten)))
erg2<-matrix(-1,nrow=3,ncol=length(Wert1))
erg3<-matrix(-1,nrow=replikate,ncol=length(Wert1))

for (ol in (1:length(daten))){
  load(daten[ol])
  if(ol %in% round(seq(1,length(daten),length.out=11))){# shows that thing is actually doing somthing.... and how many % are done
    print(noquote(paste(round(ol/length(daten)*100)," % done"))) 
  }
  
  
  # erg[,,ol]<-wata
  
  # fitself[which(is.na(fitself))]<-0
  # fitmate[which(is.na(fitmate))]<-0
  # fm[ol,1:length(fitmate)]<-fitmate
  # fs[ol,1:length(fitself)]<-fitself
  # fit1[ol,]<-tifo#fits
  # fit2[ol,]<-fits#tifo_late#fitti
  delis[which(Wert1==W1),]<-delis[which(Wert1==W1),]+defr
  # waitgen[,,ol]<-apply(varii,2:3,mean)
  # fitfin[ol,]<-shots
  popi[which(Wert1==W1),1:length(nu)]<-popi[which(Wert1==W1),1:length(nu)]+nu
  waiti[which(Wert1==W1),1:length(waitmean)]<-waiti[which(Wert1==W1),1:length(waitmean)]+waitmean
  waiti_vari[which(Wert1==W1)]<-waiti_vari[which(Wert1==W1)]+waitvari[length(waitvari)]
  waiti_vari2[which(Wert1==W1),1:length(waitvari)]<-waiti_vari2[which(Wert1==W1),1:length(waitvari)]+waitvari
  # waiti_vari[which(Wert1==W1),1:length(waitvari)]<-waiti[which(Wert1==W1),1:length(waitvari)]+waitvari
  
  pp[which(Wert1==W1)]<-pp[which(Wert1==W1)]+sesult
  # if (length(nu)>=times){
  #   extinction[ol]<-NA
  # }else {
  #   extinction[ol]<-length(nu)}
  W[ol]<-W1
  jup[which(Wert1==W1)]<-jup[which(Wert1==W1)]+1
  wiria[,which(Wert1==W1),jup[which(Wert1==W1)]]<-wira[dim(wira)[1],]
  oria[,which(Wert1==W1),jup[which(Wert1==W1)]]<-ori[dim(ori)[1],]
  # if (!(is.null(Bself) & is.null(Bmate))){
  #   self1[ol,]<-Bself
  #   mate1[ol,]<-Bmate
  # }
  
  erg2[,which(Wert1==W1)]<- erg2[,which(Wert1==W1)]+colMeans(wata[(dim(wata)[1]*.95):dim(wata)[1],])
  erg3[min(which(erg3[,which(Wert1==W1)]==-1)),which(Wert1==W1)]<-mean(wantthat[(length(wantthat)*.95):length(wantthat)])
  
  
  rm(list=setdiff(ls(), c("o","k","a","aa","b","bb","erg","erg2","erg3","QQ","ffff","jup","oria","waiti_vari2","waiti_vari","nAme","Faktor","wiria","delis","waitgen","pp","times","fitfin","ind","fit1","fit2","fm","fs","self1","mate1","W","extinction","waiti","popi","Wert1","replikate","times","daten","ol")))
}
extinction[which(extinction>=times)]=NA
w1<-sort(unique(W))

rotate <- function(x) t(apply(x, 2, rev))
# for (u in 1:length(daten)){
# load(daten[u]); print(max(result[["Birth and death times"]][["Times of Birth by Selfing"]]))}
library("viridis")
real<-numeric(length(w1))
for (i in 1:length(w1)){
  real[i]<-sum(W==w1[i])
}

delis<-delis/real
popi<-popi/real
waiti<-waiti/real
pp<-pp/real
waiti_vari<-waiti_vari/real
waiti_CV<-sqrt(waiti_vari)/waiti[,dim(waiti)[2]]

waiti_vari2<-waiti_vari2/real
waiti_CV2<-sqrt(waiti_vari2)/waiti

sd_vari<-sqrt(waiti_vari/sqrt(real))
sd_vari<-sqrt(waiti_vari)/sqrt(real)
sd_CV<-sqrt(waiti_CV)/sqrt(real)

#-----------------Viability Alleles---------------------------------
png(filename = "Graphs%03d.png",width=1905,height=662)

defris<-cbind((1-delis),delis/10)
matplot(w1,defris,pch=19,lty=1,type='l',lwd=3,col=c("forestgreen","firebrick2"),xlab=Faktor,ylab="Frequency",ylim=c(0,1))
matplot(w1,defris,pch=19,lwd=3,col=c("green3","tomato1"),xlab=Faktor,ylab="Frequency",add=TRUE)         
legend("topleft",c("wildtype","deleterious"),col=c("green3","tomato1"),pch=15,bg="white",cex=1.5)

#waiting time and final----------------
waiter<-t(waiti)

fin<-numeric(length(w1))


fin<-rowMeans(waiti[,(times-(times*.05)):times])

print(matplot(waiter,col=viridis(dim(waiter)[2]),type='l',lty=1,lwd=3,ylab="average waiting time",xlab="time",main="Waiting time"))
legend("topleft",as.character(w1),cex=1,col=viridis(dim(waiter)[2]),pch=15,title=Faktor,ncol=3,bg="white")


plot(w1,t(fin), xlab=Faktor, ylab= "final waiting time",pch=19,cex=1.3,type='b',lty=3)
abline(h=mean(waiti[,1]))


graphics.off()
save(list=ls(),file="compressed.RData")
```
